# The genome and transcriptome of the snail *Biomphalaria sudanica s.l.*: Immune gene diversification and highly polymorphic genomic regions in an important African vector of *Schistosoma mansoni*

**DOI:** 10.1101/2023.11.01.565203

**Authors:** Tom Pennance, Javier Calvelo, Jacob A. Tennessen, Ryan Burd, Jared Cayton, Stephanie R. Bollmann, Michael S. Blouin, Johannie M. Spaan, Federico G Hoffmann, George Ogara, Fredrick Rawago, Kennedy Andiego, Boaz Mulonga, Meredith Odhiambo, Eric S. Loker, Martina R. Laidemitt, Lijun Lu, Andrés Iriarte, Maurice Odiere, Michelle L. Steinauer

## Abstract

**Background:** Control and elimination of schistosomiasis is an arduous task, with current strategies proving inadequate to break transmission. Exploration of genetic approaches to interrupt *Schistosoma mansoni* transmission, the causative agent for human intestinal schistosomiasis in sub-Saharan Africa and South America, has led to genomic research of the snail vector hosts of the genus *Biomphalaria*. Few complete genomic resources exist, with African *Biomphalaria* species being particularly underrepresented despite this being where the majority of *S. mansoni* infections occur. Here we generate and annotate the first genome assembly of *Biomphalaria sudanica* sensu lato, a species responsible for *S. mansoni* transmission in lake and marsh habitats of the African Rift Valley. Supported by whole-genome diversity data among five inbred lines, we describe orthologs of immune-relevant gene regions in the South American vector *B. glabrata* and present a bioinformatic pipeline to identify candidate novel pathogen recognition receptors (PRRs).

**Results:** *De novo* genome and transcriptome assembly of inbred *B. sudanica* originating from the shoreline of Lake Victoria (Kisumu, Kenya) resulted in a haploid genome size of ∼944.2 Mb (6732 fragments, N50=1.067 Mb), comprising 23,598 genes (BUSCO=93.6% complete). The *B. sudanica* genome contains orthologues to all described immune genes/regions tied to protection against *S. mansoni* in *B. glabrata*. The *B. sudanica PTC2* candidate immune genomic region contained many PRR-like genes across a much wider genomic region than has been shown in *B. glabrata*, as well as a large inversion between species. High levels of intra-species nucleotide diversity were seen in *PTC2*, as well as in regions linked to *PTC1* and *RADres* orthologues. Immune related and putative PRR gene families were significantly over-represented in the sub-set of *B. sudanica* genes determined as hyperdiverse, including high extracellular diversity in transmembrane genes, which could be under pathogen-mediated balancing selection. However, no overall expansion in immunity related genes were seen in African compared to South American lineages.

**Conclusions:** The *B. sudanica* genome and analyses presented here will facilitate future research in vector immune defense mechanisms against pathogens. This genomic/transcriptomic resource provides necessary data for the future development of molecular snail vector control/surveillance tools, facilitating schistosome transmission interruption mechanisms in Africa.

## Background

Freshwater pulmonate snails of the genus *Biomphalaria* are intermediate hosts for a diversity of trematode parasites but are most notorious for their role in the transmission of *Schistosoma mansoni*, to humans. Schistosomiasis is a chronic and inflammatory disease with devastating impacts to human health, which likely are underestimated due to schistosomiasis associated morbidities and mortalities being attributed to non-communicable diseases for which the symptoms are similar (1). Despite the widely recognized toll of schistosomiasis on human health, there are few effective and implementable options for controlling transmission of the parasite (2).

The need for novel interventions to interrupt *S. mansoni* transmission has spurred on investigatory genomic research of the *Biomphalaria* snail vector hosts (3). Genomic resources for *Biomphalaria* will facilitate the discovery of genetic resistance mechanisms to schistosome infection, which could be manipulated to block transmission to snails, and thus humans (4). Currently, three *Biomphalaria* species represent the extent of genome resources, two South American species: *Biomphalaria glabrata* (5,6); *Biomphalaria straminea* (7); and one sub-Saharan African species *Biomphalaria pfeifferi* (8). African *Biomphalaria* species are therefore underrepresented in terms of complete genomic information, even though African species contribute to the vast majority of global *S. mansoni* transmission since approximately 90% of human infections occur in Africa. Thus, genome-wide analysis of African *Biomphalaria* species would facilitate the development of genetic based vector control in areas where it is highly relevant to transmission. As more *Biomphalaria* genomes become available, evolutionary analysis of immunity of these major vectors will be possible, whilst putting it into the context of species divergence across the African continent after being introduced from South America sometime between 1.8 and 5 MYA (8–11).

The African species *B. sudanica* was originally described in 1870 from Djur and Rek tributaries of the White Nile in the Bahr el Ghazal region of Southern Sudan (12). It is distributed throughout the Nile Basin in marsh and lacustrine habitats in Uganda, Kenya, Sudan, Tanzania and Ethiopia (13–18). *Biomphalaria sudanica* distribution in East Africa corresponds to the geographic region where the genetic diversity of *S. mansoni* is the greatest (19,20). Most research regarding *B. sudanica* has been focused on populations from Lake Victoria, where *S. mansoni* remains highly endemic even following repeated and widespread mass drug administration of schistosomiasis preventative chemotherapy (21–23). Although *B. sudanica* inhabits the marshy fringes and nearshore shallow waters of Lake Victoria where human-freshwater contact takes place (24), another snail vector of *S. mansoni*, described as *B. choanomphala* (25), occurs in deep water habitats of Lake Victoria (13,26,27). The taxonomic status of these two species of Lake Victoria snails is in question, as DNA divergence of mitochondrial genes (and the few nuclear genes that have been sequenced) between these species suggests they may represent ecomorphs of a single species (28–30). However, distinct morphologies, habitats, and schistosome susceptibility profiles (31), make the distinction of these two forms critical in the context of a genome report. Thus, we follow conventional use of the species name, or *Biomphalaria sudanica* sensu lato. Genomic data of these species will facilitate future population genomic analyses aimed at better understanding the relationship between these taxa. Snail based schistosomiasis control cannot even be imagined without understanding these basics.

Experimental infections have shown that *B. sudanica* displays the greatest natural resistance to schistosome infection relative to *B. choanomphala*, and another closely related species, *B. pfeifferi* (27,31), and thus offers an excellent target for the discovery of immune relevant genes in an African *Biomphalaria* species. While some immune genes can be characterized by conserved domains as a result of positive selection (32), others such as those involved in host-pathogen interactions are rapidly evolving under balancing selection due to the simultaneous arms races occurring between the host and its pathogens (33). Indeed, immune loci are among the most diverse in many genomes, including the classic example of the vertebrate major histocompatibility complex (MHC) (34,35); human innate immunity genes (36); R genes in plants (37,38) and more recently shown in immune genes of invertebrate organisms such as *Caenorhabditis elegans* (39). Virtually nothing is known regarding the *B. sudanica* immune defense except for what can be inferred from orthologous gene searching strategies related to experimental work with the South American congener, *B. glabrata* (3). The identification of *B. glabrata* loci associated with resistance to schistosomes is an active research field with approaches such as Quantitative Trait Locus (QTL) analysis providing valuable new insights (6).

In this paper, we present the first description of the genome of *B. sudanica*. Our novel annotated genome of *B. sudanica* 111 (Bs111), an inbred line maintained at Western University of Health Sciences which originates from the Kisumu region (Kenya) of Lake Victoria, comprises PacBio HiFi long-read DNA and RNA sequence data, as well as Illumina short-read RNA sequence data. This combination of genomic and transcriptomic data provides a confident annotation of functional gene boundaries, exon-intron structure, and isoforms for the representative genome of this species. Here, we focus on identifying and describing gene regions orthologous to those involved in immunity of the South American vector *B. glabrata* to *S. mansoni*, such as the *PTC1* and *PTC2* genomic regions (40,41) and fibrinogen-related proteins (FREPs) (42,43). We also used a new analysis pipeline to find novel pathogen recognition receptors (PRRs). Following the hypothesis that PRRs are under balancing selection as with other highly polymorphic immune loci, we searched in the most hyperdiverse genome regions through the genomic comparison of five genetic lines of *B. sudanica* for signatures of candidate PRRs. This allows us to identify immune related genes that do not maintain detectable sequence similarity with known gene families and are not only specific to schistosome immunity. The description of key features of the *B. sudanica* genome provides multiple exciting avenues for future research into this important vector of *S. mansoni*.

## Results

### Biomphalaria sudanica 111 line genome assembly and nuclear genome annotation

The PacBio assembled *B. sudanica* (Bs111) haploid genome size is ∼944.2 Mb, comprising 6732 contigs and scaffolds with an N50 of 1.067 Mb, and a mean sequencing coverage of ∼23x (Supplementary Table 1). The estimated size of the *B. sudanica* genome is somewhat larger than those of *B. pfeifferi* (∼771.8 Mb (8)) and *B. glabrata* (iM line: ∼871.0 Mb (6)), but smaller than that of *B. straminea* (∼1,004.7 Mb (7)).

PacBio and Illumina RNA sequence data were obtained from Bs111 snails to aid in annotation of the assembled *B. sudanica* genome. Pooled RNA was processed following a standard PacBio IsoSeq procedure, which yielded 335.0 Gbases (N50 of ∼115.5 Kbases) of long-read transcript data. Of the original 6,945,781 circular consensus sequencing (ccs) reads, 3,798,283 (54.68%) passed the Q20 quality threshold determined in the longQC software (44), of which 1,708,667 (44.99%) were identified as potentially complete isoforms (i.e. they bear both 5’ and 3’ adapter sequences) using Lima (github.com/PacificBiosciences/barcoding) (Supplementary Table 1). To supplement the RNA transcript long-read data, Illumina paired-end 150 short-read sequence data yielded 45,125,478 paired reads of which 44,008,723 (97.53%) passed the trimming process conducted in Trimommatic (45) (Supplementary Table 1). Overall mapping rate of long-read transcripts using minimap2 (46,47) and Illumina RNA short-reads using STAR (48) to the assembled genome was close to 100%. Transcript characterization using StringTie2 v2.2.1 (49) identified 25,847 individual genes, of which 23,598 in TransDecoder.Predict v5.5 (50) had an assigned open reading frame (ORF) (Supplementary File 1 and Supplementary File 2). InterProScan v5.56-89.0 (51) identified at least one protein domain signature on 19,945 of the 23,598 genes (Supplementary File 3). BUSCO (52) completeness analysis shows that the latter set (23,598 genes) represents a close to complete genome annotation relative to mollusca_odb10 (of 5295 BUSCO groups, 93.6% complete, 1.4% fragments and 5.0% missing).

The *B. sudanica* genome contains orthologues to at least 18 candidate immune loci of *B. glabrata* that function in protection against *S. mansoni* (Supplementary Table 2). These include key genes/gene clusters coding for FREP2 and FREP3, *B. glabrata toll-like receptor* (*BgTLR*), *Phox*, Guadeloupe Resistance Complex 1 genes (*GRC*) – referred to from here as the polymorphic transmembrane cluster 1 (*PTC*1), polymorphic transmembrane cluster 2 (*PTC*2), *RADres*, heat shock protein 90 (*HSP*90), *Granulin* (*GRN*), *BgTEP*, Catalase (*cat*), *Biomphalysin*, *Glabralysin*, OPM-04 (Knight marker), superoxide dismutase 1 (*sod*1), *Peroxiredoxin* (*prx*4), qRS-5.1 and qRS-2.1 (Supplementary Table 2).

A total of 919 tRNA and 107 rRNA genes were predicted in the *B. sudanica* nuclear genome (Supplementary Table 3, Supplementary Table 4 and Supplementary File 1). The number of tRNA genes identified in the *B. sudanica* genome is comparably higher than that observed in the genomes of *B. pfeifferi* (n=514 (8)) and *B. glabrata* (n=510 (6)). As is the case for *B. pfeifferi*, one selenocysteinyl tRNA (tRNA-SeC) is present in the genome of *B. sudanica* (Supplementary Table 3 and Supplementary File 1), meaning this species is capable of synthesizing selenocysteine containing polypeptides, or selenoproteins (53,54). The tRNA-Sec gene has not been identified in *B. glabrata* (8). Overall, fewer rRNA genes were predicted in this genome assembly of *B. sudanica* compared to that of *B. pfeifferi* (107 and 757, respectively), which could be a result of some rRNA genes being misassembled in *B. sudanica*.

As with other *Biomphalaria*, repetitive elements composed a large proportion of the *B. sudanica* genome (40.3%) (Supplementary Figure 1). About 87% of protein coding genes overlap with at least one annotated repeated element in their gene model. The overlap is primarily within introns and untranslated regions (UTRs); however, 1576 genes have repeat elements within their predicted coding sequence (CDS) (Supplementary Table 5). The repeat regions largely comprise unknown repeat elements, in addition to an abundance of unclassified long interspersed nuclear elements (LINE), LINE/retrotransposable element Bovine B (RTE-BovB) and unclassified DNA transposons (Supplementary Figure 1, Supplementary Table 6).

### Mitochondrial genome annotation and trimming processes

The mitochondrial genome comprises the same gene content (13 genes, 3 rRNA, 22 tRNA), and synteny as its congeners (55) (Supplementary Table 7). The mitochondrial genomes of gastropods are unique in the fact that they have acquired transcriptional processes during their evolutionary history that are not often observed in vertebrates (56). Therefore, while an annotated mitochondrial genome of *B. sudanica* has been previously published (55), our novel contribution here is to validate gene boundaries and explore the transcription processes using long-read transcriptomic data.

Raw q20 PacBio IsoSeq reads were mapped to the mitochondrial genome with minimap2 (46,47). To explore the trimming process of the primary mitochondrial transcript, the intermediary pre-mRNA mitochondrial transcripts, i.e., that cover multiple features within the mitochondrial genome, were recovered and counted (Supplementary Figure 2). In line with the tRNA punctuation model (57), we established that pre-mRNAs of the *B. sudanica* mitochondrial genome are trimmed at the tRNA genes (with the potential exception of *atp6*/*atp8*), with the minus strand being processed 3’-to-5’ while the plus strand shows an odd mixture of both directions.

In *B. sudanica*, there are three mitochondrial gene re-arrangements in comparison to a typical animal mitochondrial genome that affect the transcription processes (see (58)). First, it is typical for *atp6* and *atp8* to be adjacent and remain together in the mature mRNA; however, in *Biomphalaria*, including *B. sudanica*, these genes are separated by a tRNA gene (trnN, see Fragment 5; Supplementary Figure 2 and (55)). In the case of *atp6*/*atp8*, monocistronic transcription of each of these genes following the cleavage of trnN is expected. Despite this, only a few monocistronic transcripts of each *atp6* (n=16) and *atp8* (n=1) were identified from the raw RNA reads, in contrast with the far more abundant untrimmed intermediaries (n=145; Supplementary Figure 2). Considering that the ancestral condition is the translation of both proteins from a bicistronic mRNA, it is a tempting hypothesis that this is still the case in *Biomphalaria* despite the extra trimming points.

Second, it appears that, *nad4l* is either a non-functional pseudogene in the *B. sudanica* mitochondrial genome or is only expressed in very low levels. The adjacency and bicistronic transcription of *nad4*/*nad4l*, is well conserved in invertebrate and vertebrate mitochondrial genomes, yet in many molluscan lineages, including *B. sudanica*, these genes are nonadjacent (Supplementary Figure 2) (56). Additionally, the transcript data demonstrates that *nad4* terminates on an abbreviated stop codon (T--) as was experimentally supported in its South American sister species *B. glabrata* (59), and was abundantly represented in the transcriptome (n=83 monocistronic transcripts from Fragment 3; Supplementary Figure 2). On the other hand, *nad4l* transcripts were rare, represented by only four intermediary reads that were attached to transcripts for neighboring gene Cytochrome B (*cob*) (Fragment 1, Supplementary Figure 2).

Lastly, *nad6*/*nad5*/*nad1* are found adjacent to one another and seem to be translated as a polycistronic mRNA (Fragment 1, Supplementary Figure 2), since their gene boundaries predicted by MITOS2 (60) overlap; there are no clear cuts in the read coverage; and pure monocistronic reads were only recovered in small numbers for *nad5* (n=16) and *nad1* (n=2). Given the lack of a clear trimming point in this region, the observed monocistronic and bicistronic reads (e.g. *nad6*/*nad5* or *nad5*/*nad1*) are likely the product of partial RNA degradation, implying that the three genes are translated into proteins from the same mRNA molecule.

### Location signals and transmembrane domains

Location signals for exportation, i.e. the signal peptides or mitochondrial targeting peptides, were identified with SignalP v6.0 (61) and TargetP v2.0 (62) in 3,339 genes (5,016 isoforms) and 69 genes (111 isoforms), respectively (Supplementary Table 8). Transmembrane domains were predicted in 4,922 genes (8,728 isoforms), and the location signals analysis suggests that 835 of these were firmly anchored to the plasma membrane, organellar membranes or vesicles, and seven to the mitochondria (Supplementary File 4).

An additional 82 proteins (146 isoforms) that showed no location signals in SignalP and TargetP were identified by SecretomeP (63) as potentially secreted through an alternative pathway. However, since 34 of these had at least one transmembrane domain, we suspect many of these are false negatives for either the signal or the mitochondrial targeting peptide. Furthermore four genes have signal peptides predicted in some but not all their isoforms (genes BSUD.7093, BSUD.10729, BSUD.12693 and BSUD.24440). This might represent cases of functional isoforms generated through alternative splicing that have different locations in the cell, as observed in other species (64).

Two identical secreted proteins worth pointing out, BSUD.4529 (contig 217) and BSUD.14556 (contig 559), were identified as orthologs to the precursor protein of peptide P12 in *B. glabrata* (BGLB027975), which has been shown to trigger behavior modifications in *S. mansoni* miracidia, and thus is potentially an attractant (65,66). Compared to the *B. glabrata* P12, the ortholog in *B. sudanica* contains a non-synonymous change within the 13 aa region, changing the 5^th^ amino acid from Glycine to Valine (DITS<u>V</u>LDPEVADD).

### Gene family evolution in Biomphalaria species

The evolutionary dynamics of *Biomphalaria* protein families among *B. sudanica, B. glabrata*, *B. pfeifferi*, *B. straminea*, *Bulinus truncatus* and *Elysia marginata* were estimated by first identifying orthology in Phylogenetic Hierarchical Orthogroups (HOG) with Orthofinder v2.5.4 (67). Orthofinder clustered ∼87% of identified genes into 29,664 orthogroups with representatives in at least 2 species. Orthogroups were distributed across 31,723 HOGs (Supplementary Table 9). HOGs were taken as an approximate estimation of protein families found on these genomes, and significant changes in their size estimated with CAFE 5 (68). The phylogeny utilized for the estimation was the species tree generated by Orthofinder for the HOG definition, including 3541 single copy orthogroups with representatives in all the genomes, and the root calibrated to be 20 Million Years based on the appearance of *Bulinus* in the fossil record 19-20 MYA (69) (see Figure 1). Gene Ontology (GO) terms were assigned to each HOG with eggNOG-mapper v2.0 (70,71) assuming that a GO assigned to one member applied to the whole HOG (Supplementary Table 10), significantly enriched GO terms among the expanded and contracted HOGs on each node were identified with topGO (72) (Supplementary Table 11) and then summarized with REVIGO (73) to allow further interpretation (Supplementary Table 12 Supplementary File 5).

**Figure 1.**
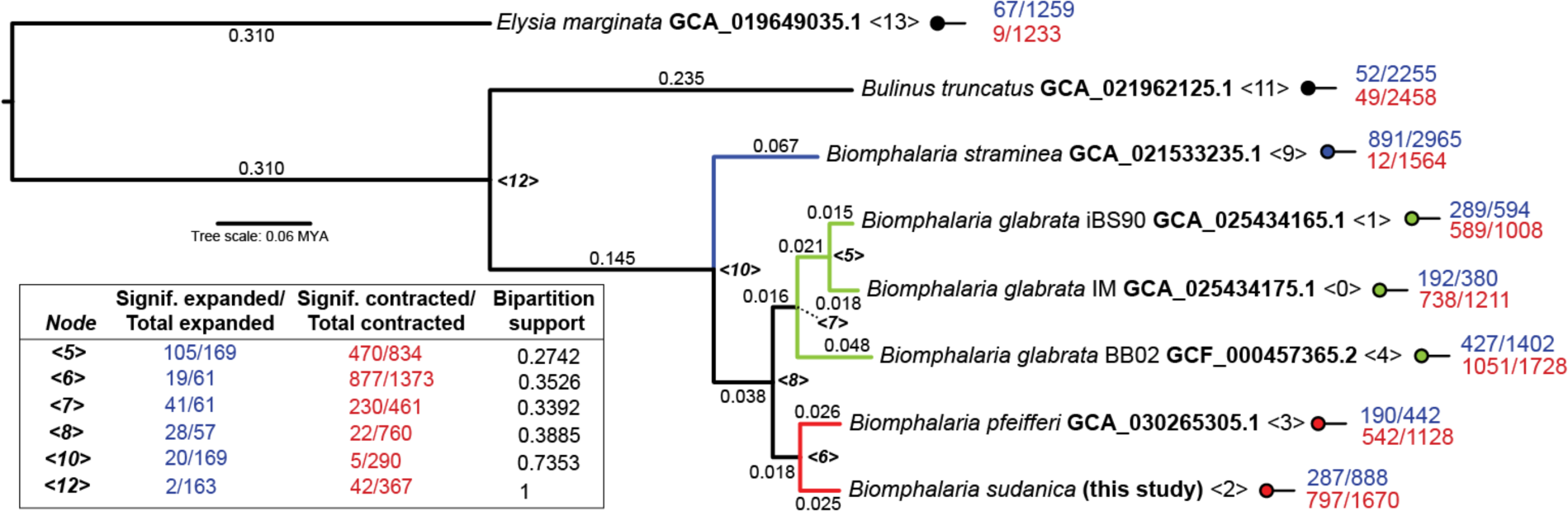
Species tree generated in Orthofinder using the Species Tree of All Genes (STAG) algorithm (67,74). Root is time calibrated to be 20 Million Years Ago based on appearance of *Bulinus* in the fossil record (69). Node support values represent the bipartition proportions in each of the individual species tree estimates. Branch lengths represent the average number of substitutions per site across all the individual trees inferred from each gene family. The number of (significant/total) gene families expanded (blue) and contracted (red) in the ancestral populations of the *Biomphalaria* species, and outgroups *Elysia marginata* and *Bulinus truncatus* as determined in CAFE 5 (68) are shown for each internal and terminal node (<0> to <13>).

Significant gene family expansions/contractions were identified throughout the analyzed phylogeny. Lineage-specific trends were identified within the *Biomphalaria* genus, in comparison to outgroups, between South American and African *Biomphalaria* species, within the African lineages (*B. sudanica* and *B. pfeifferi*), and between lines of *B. glabrata* (Figure 1, Supplementary Table 12 and Supplementary File 5). Expansions in immune related gene families were detected in the common ancestor of all *Biomphalaria* species relative to the previous node representing the split from *Bulinus truncatus*, including those involved in acute inflammatory response, glomerular filtration and regulation of cell adhesion (see node 10, Figure 1 and Supplementary File 5 and Supplementary Table 12).

In the branch leading to the common ancestor of the African species, *B. sudanica* and *B. pfeifferi*, we identified substantially more gene family contractions than expansions (see node 6, Figure 1). Within this lineage we found the expansion or contraction of gene families associated with the circulatory system (e.g. GO:0001525, GO:0001987, GO:0007512 and GO:0061337), compound transport (e.g. GO:0098660, GO:0006811, GO:0035459 and GO:0006855), protein maturation (e.g. GO:0006508, GO:0036211 and GO:0016579), metabolic pathways and regulation (e.g. GO:1901293, GO:0006210, GO:0006212 and GO:0006164), development and growth (e.g. GO:0050767, GO:0001558 and GO:0036342), and protection towards environmental stressors including chemical stressors (e.g. GO:0009620, GO:0009636, GO:0010033 and GO:0010035) (Supplementary Table 12 for full details). Several genes associated with chemotaxis (e.g. GO:0006935, GO:0009410 and GO:0071310), which are often associated with immunity, were contracted in the African *Biomphalaria* (Supplementary Table 12). Regarding the *B. sudanica* lineage, gene families belonging to 287 GO terms were significantly expanded, including genes associated with the regulation of hippo signaling (See Node 2, Group 4 in Supplementary File 5 and Supplementary Table 12) that account for multiple immune response processes. However, many GO terms associated with the immune response were also identified in contracted gene families in *B. sudanica*, including defense responses and regulation of immune system processes suggesting that a complex evolutionary trend took place in this species.

### Identification and phylogeny of variable immunoglobulin and lectin domain-containing molecules (VIgLs): FREPs and CREPs

Genes putatively belonging to FREP, C-type lectin-related protein (CREP) or galectin-related protein (GREP) families were predicted based on the presence of a secretion signal and in conjunction with either fibrinogen (FBD), C-type lectin, or Galectin as predicted by InterProScan v5.56-89.0 (51) (Supplementary File 3) and hmmsearch v3.3.2 (75) searches with custom IgSF profiles (as described in (43)).

Following this selection pipeline, 246 genes distributed among 140 HOGs were determined to have key domains that made them FREP, CREP or GREP candidates (Supplementary Table 13). Upon cross-checking our candidate protein domains, none of the seven initial GREP candidates (i.e. Galectin domain-containing proteins) were identified to contain putative IgSF or other immunoglobin domains (Supplementary Table 13).

Unlike the FBD-containing proteins (i.e. candidate FREPs), proteins bearing C-type lectin domains showed a considerable structural diversity (i.e. displaying several domains non-related with the CREP family). Some of these genes may participate in the snail immune response, as they often had signatures compatible with IgSF domains that were identified by InterPro (accessions: IPR003598, IPR003599, IPR007110, IPR013098, IPR013783 and IPR036179), but were not determined as members of the CREP family following previously described criteria (76). Considering the hallmarks of a CREP or FREP gene (signal peptide present plus one or two IgSF domains), a total of 10 potential CREP and 57 FREP (Supplementary Table 14) genes were identified (Table 1 and Supplementary Table 13).

**Table 1.** Summary of variable immunoglobulin and lectin domain-containing molecules (VIgLs), fibrinogen-related proteins (FREPs), C-type lectin-related protein (CREPs) and galectin-related proteins (GREPs) identified in *Biomphalaria sudanica* (this study)*, B. glabrata* and *B. pfeifferi* (8). No GREPs were identified in either of the African *Biomphalaria* species *B. sudanica* or *B. pfeifferi*. *One FREP in *B. sudanica* contained only a partial fibrinogen domain, and is considered truncated (see BSUD.19120.1). Summaries of *B. sudanica* FREP and CREP gene compositions can be found in Supplementary Table 14.

| Complete VIgLs | *B. sudanica* | *B. glabrata* | *B. pfeifferi* |
| --- | --- | --- | --- |
| Fibrinogen-related proteins (FREPs) | 57* | 39 | 55 |
| C-type lectin-related proteins (CREPs) | 10 | 4 | 11 |
| Galectin-related proteins (GREPs) | 0 | 1 | 0 |

The selected full CREP and FREP genes were aligned with a set of reference sequences from *B. glabrata* (Supplementary Table 15), their phylogeny relations estimated by maximum likelihood and annotated based on their position in the phylogeny (Figure 2 and Figure 3). Thirty-nine of these reference sequences were reported and curated previously (43), while 17 were annotated as members of these gene families (FREP/CREP) and available on the National Center for Biotechnology Information (NCBI) (Supplementary Table 15). The CREP family is divided into two main monophyletic groups, one of which includes all the reference sequences (Figure 2). In this group containing reference sequences, the HOGs associated with CREPs identified for *B. sudanica* were N0.HOG0002779 and N0.HOG0016618, which are closely related with the references CREP1 and CREP3, respectively. The other monophyletic group, which comprise five HOGs, is formed exclusively by divergent *B. sudanica* CREPs, possibly representing new subtypes within the CREP family.

**Figure 2.**
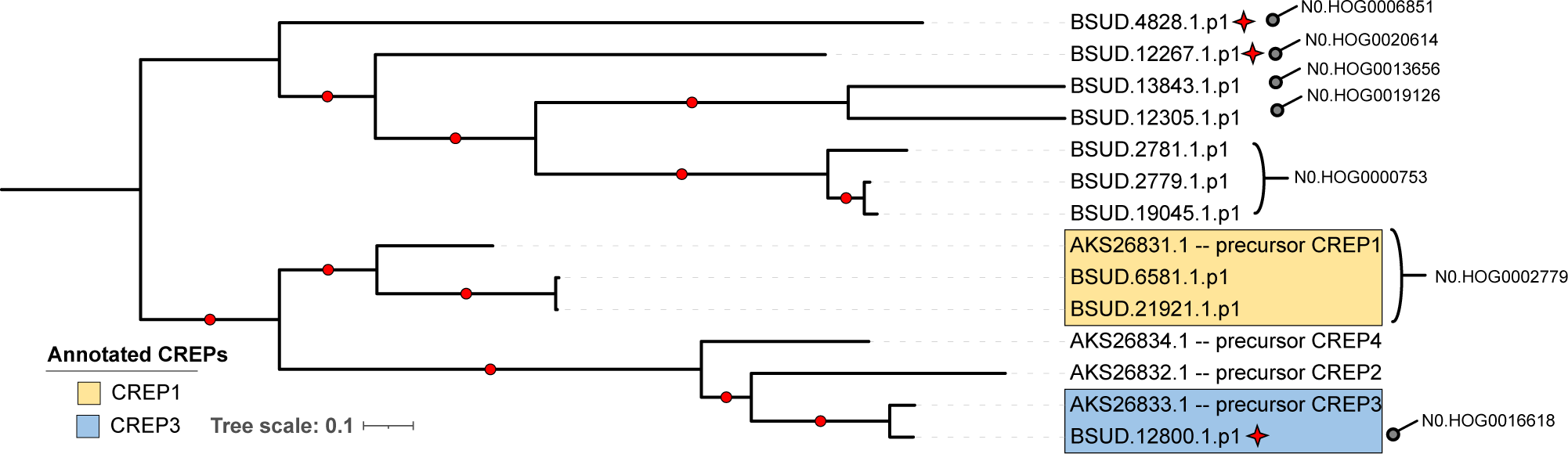
Maximum likelihood tree of C-type lectin-related proteins (CREPs) identified from *Biomphalaria sudanica* in the current study (see Supplementary Table 14) and four CREPs identified previously from *B. glabrata* (see Supplementary Table 15) organized within hierarchical orthogroups (HOGs) as determined by Orthofinder (67). Branch lengths represent the number of substitutions per site. Nodes with bootstrap values >75 (estimated with 1000 replicates of non-parametric bootstrap) are signified by a red dot on the branch before bipartition. Red stars indicate three CREPs with unusual features, namely that they include weak hits for secondary immunoglobulin domains, which may overlap with C-lectin domains as well as containing a large interdomain region (see Supplementary Table 14).

**Figure 3.**
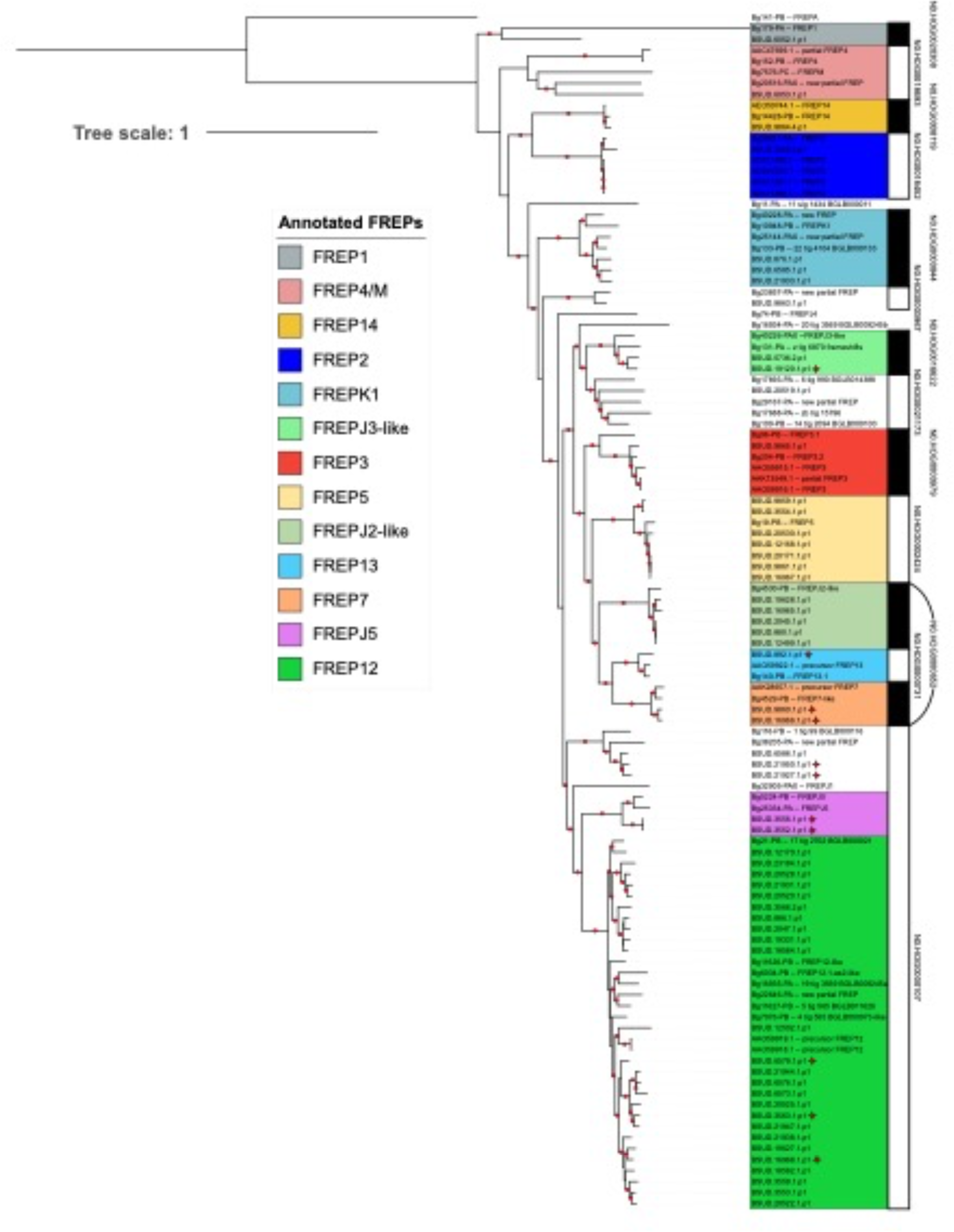
Maximum likelihood tree of the 57 fibrinogen-related proteins (FREPs) identified from *Biomphalaria sudanica* in the current study (see Supplementary Table 14) amongst reference sequences for FREPs identified previously from *B. glabrata* (see Supplementary Table 15) organized within hierarchical orthogroups (HOGs) as determined by Orthofinder (67). Branch lengths represent the number of substitutions per site. Nodes with bootstrap values >75 (estimated with 1000 replicates of non-parametric bootstrap) are signified by a red dot on the branch before bipartition. Red stars indicate FREPs with unusual features according to our annotation summarized in Supplementary Table 14, including those containing weak hits for additional immunoglobulin (IgSF) domains (e.g. BSUD.16968), IgSF rearrangements (e.g. BSUD.21927) or containing partial fibrinogen domains (e.g. BSUD.19120).

FREP genes were grouped in thirteen HOGs, eleven of which are closely related to classified reference sequences (Figure 3). Thus, most FREPs in *B. sudanica* could be classified as a known subtype within the family, except for FREP genes belonging to N0.HOG0000107, N0.HOG0000652, N0.HOG0021173 and N0.HOG0003967. Genes from the HOG N0.HOG0000107 is by far the largest FREP subgroup in *B. sudanica* identified in this work, and it is subdivided in three monophyletic lineages, the largest lineage comprising 25 genes is associated with several reference sequences identified as FREP12. The sister clade to the FREP12 group is the smaller monophyletic FREPJ5 group and basal to these are a group of three genes from *B. sudanica* and two unclassified *B. glabrata* reference sequences, representing an undescribed FREP lineage. Another HOG containing FREPJ2 and FREP7 (N0.HOG0000652) is remarkable since it is paraphyletic (Figure 3). HOGs N0.HOG0021173 and N0.HOG0003967, each comprise a single FREP gene of *B. sudanica*, and were not associated with FREP genes previously assigned to a subgroup within the family. Lastly, HOG N0.HOG0018693 possesses reference genes annotated for FREP4 and FREPM, yet unlike N0.HOG0000107 or N0.HOG0000652, they do not form clear monophyletic groups with strong bootstrap support.

When comparing the numbers of CREP and FREP genes assigned to HOGs across the eight analyzed genomes (*Biomphalaria* spp., *Bulinus truncatus*, *Elysia marginata*), it is evident that FREPs are not only more numerous, but also their HOGs, at least for *B. sudanica*, contain more genes that did not meet our selection criteria (i.e. containing well supported functional domains, see Methods) to be considered for phylogenetic comparison (Supplementary Table 16). For example, 12 of the 42 candidate genes initially identified in N0.HOG0000107 (FREPJ5/FREP12) were rejected because of not containing complete FREP characters, suggesting these are not typical FREP genes. Secondly, the number of FREP genes assigned to each HOG varies considerably across *B. glabrata* and the African *Biomphalaria* lineages. For instance, N0.HOG0000107 is relatively abundant in the African species *B. sudanica* (n=30) and *B. pfeifferi* (n=30), but this number varies considerably between *B. glabrata* lines iM (n=6), BB02 (n=0), and iBs90 (n=19) (Supplementary Table 16). In addition, only a single copy of a FREP13 classified gene (N0.HOG0000731) is present in all *Biomphalaria* genomes except for in *B. glabrata* iBS90 where it is absent, and in *B. glabrata* BB02 where it comprises 27 different genes. A full study of both CREP and FREP diversity and evolution is beyond the scope of this study, but these differences suggest that the FREP genes are under a complex dynamic of duplication and replacement, while the CREPs are more stable, at least among the laboratory strains for which there are genomes available.

### *Intraspecific diversity of the* B. sudanica *genome: comparing four inbred lines*

In addition to the Bs111 reference genome, we conducted Illumina whole genome sequencing of four other inbred lines of *B. sudanica* and aligned them to the reference (Table 2). The line Bs163 was the most divergent, with an average genomic divergence value of 0.44% from the four other inbred lines (Bs111, Bs110, BsKEMRI and Bs5-2). The Bs163 line is also the most resistant to *S. mansoni* infection in the laboratory setting (77). Among the remaining four, pairwise divergences were similar and ranged between 0.28% and 0.41% with one exception: Bs110 and Bs111 were only 0.16% divergent showing high similarity. Among all five lines, median nucleotide diversity was 0.32% (95% CI = 0.06-0.83%). The heterozygosity per inbred line ranges from 0.11% to 0.21% (Table 2).

**Table 2.**
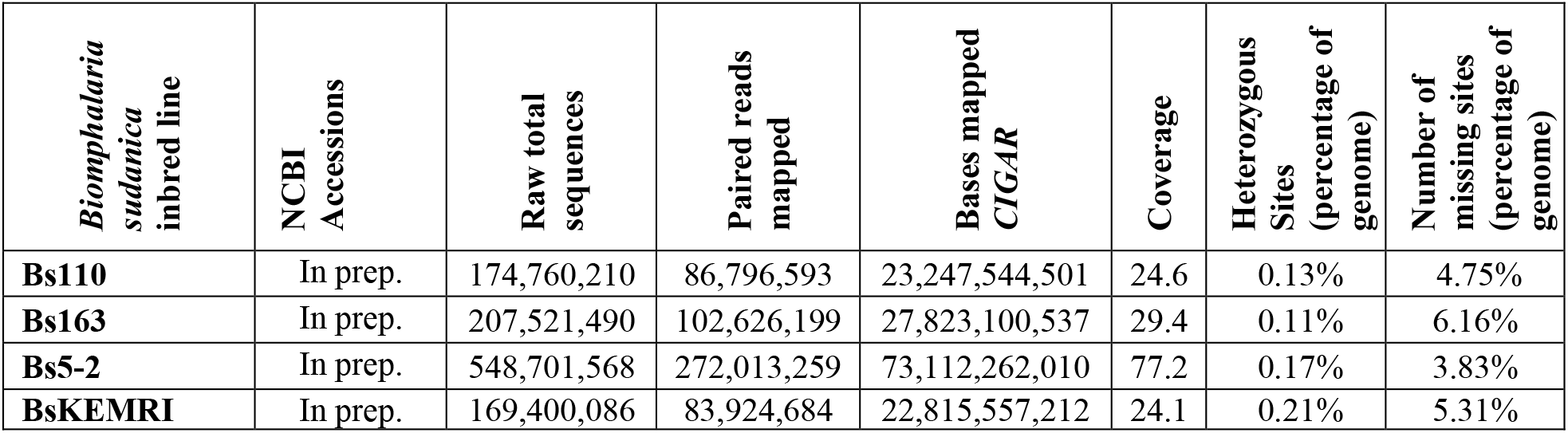
Genome statistics from Illumina sequencing (paired end 150 bp) for four *Biomphalaria sudanica* inbred lines (Bs110, Bs163, Bs5-2 and BsKEMRI) aligned to the ∼944.2 Mb *Biomphalaria sudanica* 111 reference genome.

### Characterization and categorization of highly diverse genes and genomic regions in *Biomphalaria sudanica* genome

We followed a novel bioinformatic pipeline (see Methods section: *Assessment of highly diverse genes and genome regions for novel pathogen recognition receptors*) to identify putative immune related genes (not just those involved in schistosome immunity) in the *B. sudanica* genome that may be under balancing selection, further narrowing these down to genes coding for membrane-associated proteins that could represent PRRs. We first identified genes in genomic regions that showed high divergence between the inbred lines of *B. sudanica*, hyperdiverse gene selection being based on the highest nucleotide diversity across both the entire coding and/or noncoding gene regions, and genes that were contained within the top 0.1-1% of the most nucleotide diverse sliding windows between 10-100 kb. We found multiple gene clusters that were of notably high diversity (Figure 4 and Figure 5). Following the bioinformatic pipeline, 1047 hyperdiverse genes (4.4% of all genes), from 184 different contigs/scaffolds were selected for further analysis (Supplementary Table 17).

**Figure 4.**
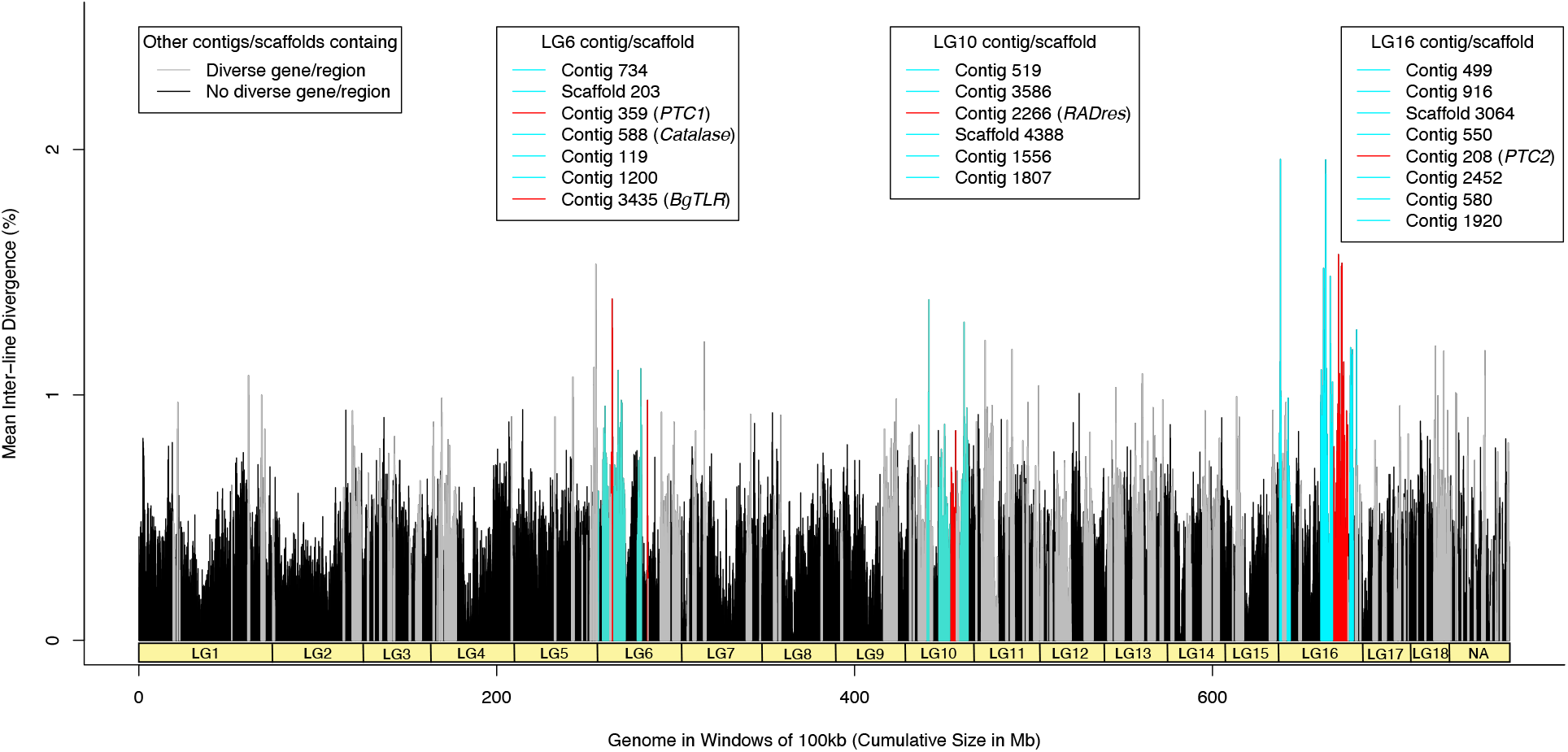
Mean nucleotide diversity of five *Biomphalaria sudanica* inbred line genomes in windows of 100 kb (0 kb stagger). Linkage groups (LG1-LG18) for *B. sudanica* are inferred from *B. glabrata* (6), showing the hypothesized chromosomal position of contigs in *B. sudanica*. Notable clusters of highly diverse genomic regions and genes were seen in *B. sudanica* linkage groups LG6, LG10 and LG16 (highlighted in turquoise and red peaks). Peaks in red represent regions containing candidate immune loci that are orthologous to some of those previously associated with *Schistosoma mansoni* resistance in *B. glabrata* (*PTC1*, *PTC2*, *Catalase*, *BgTLR*, *RADres*, see Supplementary Table 2).

**Figure 5.**
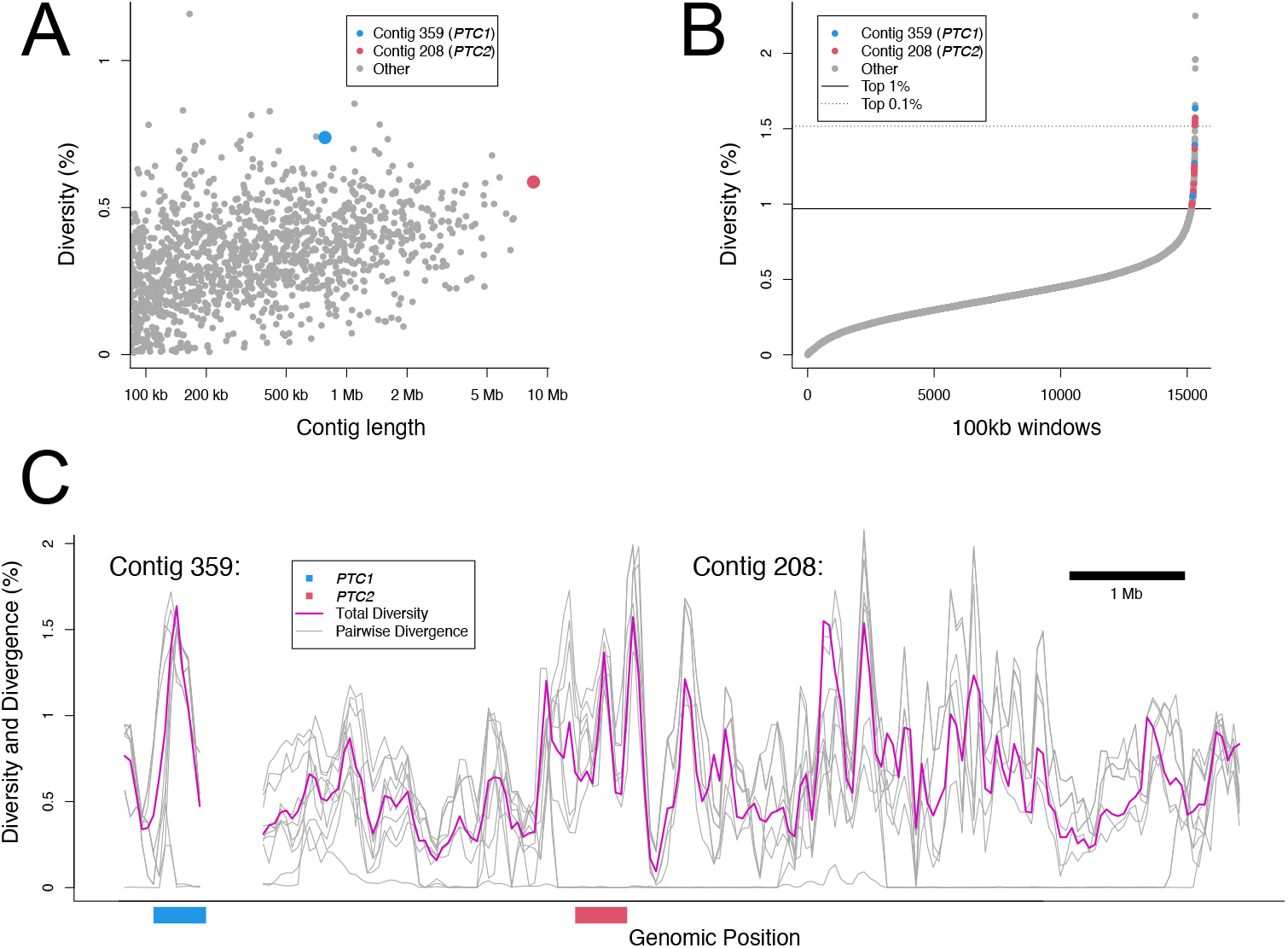
(A) Patterns of nucleotide diversity across genomes of five inbred lines of *Biomphalaria sudanica*, highlighting two polymorphic transmembrane gene clusters that are orthologous to those previously associated with *Schistosoma mansoni* resistance in *B. glabrata* (*PTC1* (40) and *PTC2* (41)). (B) Genome-wide nucleotide diversity across overlapping 100-kb genomic windows (starting at 0-kb and 50-kb intervals) with windows on contigs 359 (*PTC1*) and 208 (*PTC2*) that occur in the top 1% of genome wide nucleotide diversity between inbred lines being colored blue and red, respectively. (C) Genome-wide nucleotide diversity (purple line) and pairwise divergence for each haplotype pair (grey lines) across contigs 359 (*PTC1*) and 208 (*PTC2*). *PTC1* and *PTC2* regions are indicated by the blue and red bars, respectively. Similarly diverse regions span across several megabases of contig 208 in *B. sudanica.* Even in diverse regions, pairwise divergence can be near zero, indicating shared haplotypes.

Characterized genes in diverse regions were categorized based on their annotation descriptions (BLAST results, InterPro, Pfam, DeepTMHMM) as to what predicted role in immune function they may have. We determined 182 (17.4% of shortlist) proteins had no role in immune function (group 1) and 245 (23.4%) proteins had a role (group 2), or 138 (13.2%) a potential role (group 3), in immune function. Of the other shortlisted genes, 81 (7.7%) had an unknown function but contained transmembrane domains (TMDs) (group 4), while the remaining 401 had an unknown function (could not be interpreted due to an absence of information) without containing a TMD (group 5). In performing comparative gene ontology analysis in topGO (72), several GO terms including immune-relevant terms concerning molecular binding and receptor activity classes, CD40 receptor complexes, and defense/inflammatory/immune responses as well as regulation of innate immune responses, were significantly enriched (p<0.05 Fisher’s test) in this subset of 1047 highly diverse genes compared to the full *B. sudanica* genome (see Supplementary Table 18).

Although our novel pipeline was aimed at identifying immune genes under balancing selection in *B. sudanica* that could be due to interactions with any snail pathogen, some of those shortlisted were of relevance to candidate *S. mansoni* immune genes. Twelve of the 245 genes in diverse regions that were suspected to have immune function (group 2) in *B. sudanica* were orthologues to candidates for innate immune genes associated with *S. mansoni* resistance and likely act as PRRs in *B. glabrata*, namely *PTC1* (n=5), *PTC2* (n=6), and *BgTLR* (n=1) (Supplementary Table 17). In addition, four identified VIgLs were identified in highly diverse gene regions (Supplementary Table 17). These were FREP12 (BSUD.12502), and two neighboring FREPs on contig 777, BSUD.20519 (determined as an unknown FREP in N0.HOG0021173, Figure 3) and BSUD.20520 (grouped within FREP12 N0.HOG0000107, Figure 3). In addition, one full CREP (BSUD.13843, Figure 2 and Supplementary Table 14), was also identified within the highly diverse genes/genomic regions analysis (Supplementary Table 17). Several other immune suspected (group 2) proteins contained functional domains of interest, such as Fibrinogen-related domains (FREDs), Fibronectin, C-type lectin (incomplete CREPs) and IgSF domains.

### Pathogen recognition receptor candidates from the highly diverse genes and genomic regions in *Biomphalaria sudanica* genome

From the 1047 hyperdiverse genes identified (see above), a list of 20 proteins that contained at least one transmembrane domain with the highest coding nucleotide diversity between the five *B. sudanica* inbred line genomes were identified as potential PRR candidates (Table 3).

**Table 3.** Top 20 diverse proteins (as determined by coding nucleotide diversity) of *Biomphalaria sudanica* that contain at least 1 transmembrane domain, and thus represent putative pathogen recognition receptors. * Non-truncated BSUD.4885 protein is 405 aa in length as determined by manual alignment.

| Gene (contig/scaffold, Linkage Group) | Nucleotide Diversity % (coding) | Protein length (aa) | Inferred protein-receptor function / functional domains / protein family |
| --- | --- | --- | --- |
| BSUD.20937 (80, LG1) | 5.7 | 453 | Tumor Necrosis Factor (*TNF*) |
| BSUD.4884 (2266, LG10) | 5.3 | 86 | Unknown, linked (same contig) to *RADres* (78). Bacteriocin domain |
| BSUD.4885 (2266, LG10) | 4.8 | 354* | Unknown, linked (same contig) to *RADres* (78). Domains: TMEM154 protein family, Rifin |
| BSUD.3983 (208, LG16) | 4.0 | 817 | Unknown, linked (same contig) to *PTC2* (41). Domains: SHP2-interacting transmembrane adaptor protein, SIT |
| BSUD.4096 (208, LG16) | 3.7 | 270 | Unknown, linked (same contig) to *PTC2* (41). Domains: Viral glycoprotein L |
| BSUD.20268 (7608, LG6) | 3.3 | 75 | Unknown. Domains of unknown function (DUF6768) |
| BSUD.9255 (379, LG11) | 3.3 | 453 | Unknown Domains of unknown function (DUF4781), Death domain |
| BSUD.15077 (580, LG16) | 3.1 | 318 | Cell adhesion molecule. Wnt and fibroblast growth factor (FGF) inhibitory regulator and Protocadherin domain. [SignalP = Membrane] |
| BSUD.4112 (208, LG16) | 2.9 | 341 | Cell adhesion molecule linked (same contig) to *PTC2* (41). Protocadherin domain. [SignalP = Membrane] |
| BSUD.18104 (69, LG15) | 2.8 | 1154 | G-protein coupled receptor. 7 transmembrane receptor (rhodopsin family), Leucine-rich repeat |
| BSUD.3984 (208, LG16) | 2.7 | 324 | Polymorphic transmembrane cluster 2 (*PTC2*) gene 9 (41). [SignalP = Membrane] |
| BSUD.14211 (540, LG5) | 2.7 | 516 | Cell adhesion molecule. F5/8 type C domain. [SignalP = Membrane, Galactose-binding-like domain] |
| BSUD.17807 (6862, LG13) | 2.7 | 504 | Cell adhesion molecule. Ephrin type-A receptor 2 transmembrane and immunoglobulin domain. [SignalP = Membrane] |
| BSUD.14415 (550, LG16) | 2.6 | 483 | Unknown. Wolframin EF-hand domain |
| BSUD.9838 (3979, LG4) | 2.6 | 203 | Transient receptor potential cation channel subfamily M member 2-like. Ion transport domain |
| BSUD.12903 (499, LG16) | 2.6 | 647 | Receptor with C-type lectin and immunoglobulin domain |
| BSUD.23928 (4388, LG10) | 2.5 | 944 | G-protein coupled receptor. 7 transmembrane receptor (rhodopsin family), Low-density lipoprotein receptor domain class A, Leucine-rich repeat |
| BSUD.8884 (359, LG6) | 2.4 | 565 | Guadeloupe Resistance Complex (*GRC/PTC1*) gene 2 (40,79). Fibronectin type III domain, TMEM154 protein family. [SignalP = Membrane] |
| BSUD.4003 (208, LG16) | 2.4 | 473 | Unknown, linked (same contig) to *PTC2* (41). Prodomain subtilisin 2, TMEM154 protein family. [SignalP = Membrane] |
| BSUD.4089 (208, LG16) | 2.3 | 331 | Receptor with C-type lectin domain. [SignalP = Membrane] |

The majority (n=14) of the most diverse immune/PRR genes (Table 3) were clustered on three linkage groups: 6, 10 or 16, which were also the regions of highest diversity overall (Figure 4 and Figure 6). These linkage groups also contain genes/clusters of genes that have previously been inferred to have associations with schistosome resistance mechanisms in *B. glabrata* (*PTC1*, Catalase (*cat*), *BgTLR*, *sod*1*, prx*4, in LG6, *RADres* in LG10 and *PTC2* in LG16, see Figure 6). Direct orthologs to the candidate PRR genes *PTC2* gene 9 (BSUD.3984) and *PTC1* gene 2 (BSUD.8884) were identified in the topmost diverse PRR-like genes in the *B. sudanica* genome (Table 3).

**Figure 6.**
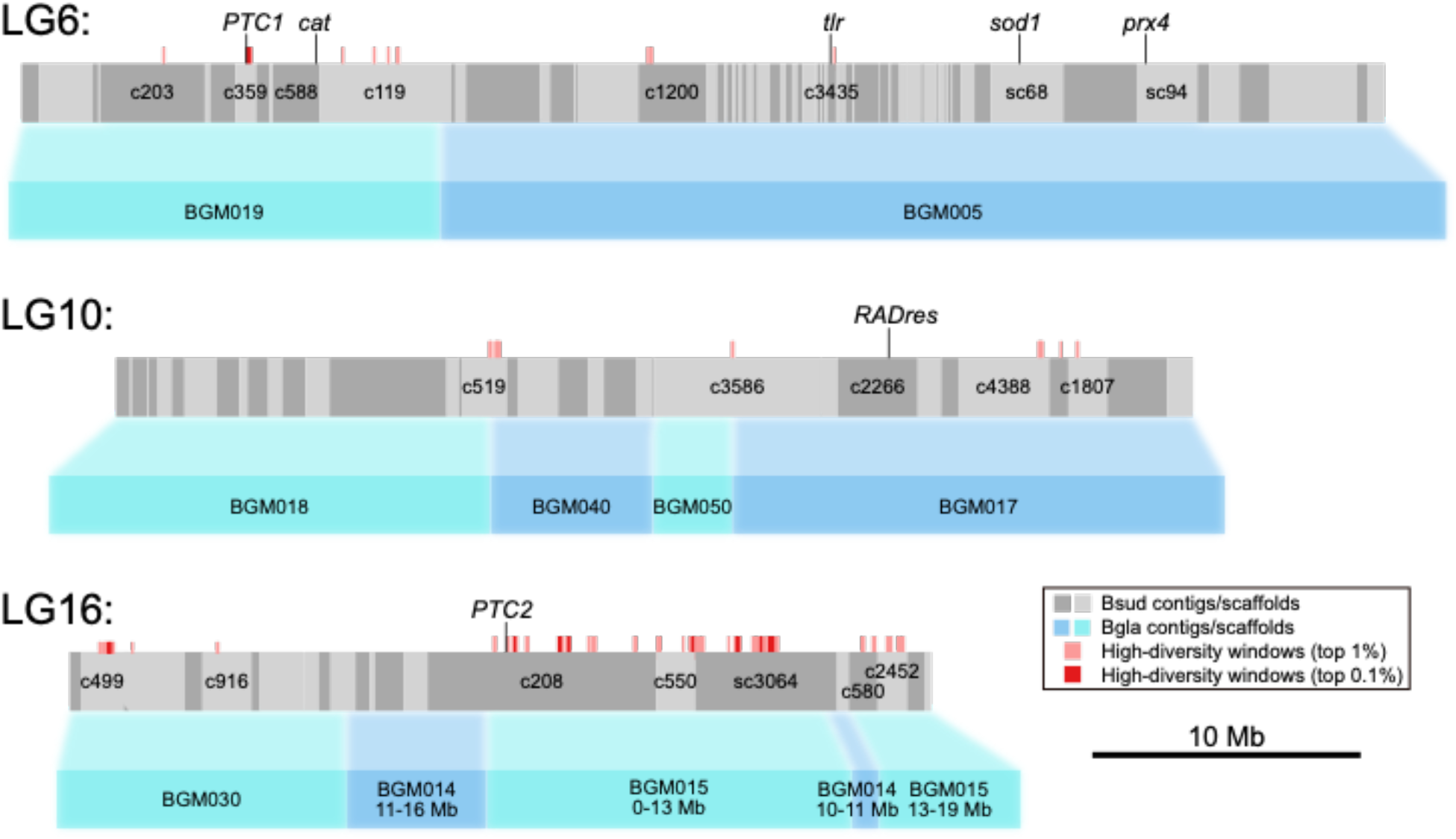
Orthology of *Biomphalaria sudanica* contigs/scaffolds (grey boxes, with different shades of grey representing alternating contigs/scaffolds) to *B. glabrata* scaffolds (blue boxes, (6)) pertaining to linkage groups (LG) inferred from three *B. glabrata* linkage groups, highlighted here because they are notably enriched for both diverse regions/genes and orthologous candidate immune genes from *B. glabrata* (*PTC1*, Catalase (*cat*), *BgTLR*, *sod1, prx4*, *RADres* and *PTC2*), positions of which are shown. Contigs/scaffolds with candidate genes and/or diverse regions (100 kb windows in top 1%, light red boxes, or top 0.1%, dark red boxes) are labeled. Synteny is relatively high between species, except for a large rearrangement on LG16 (see Supplementary Figure 3C).

### Pathogen recognition receptor candidates within Linkage Group 6

The order of genes in LG6 was highly conserved (Supplementary Figure 3A). Within LG6, the hyperdiverse protein, BSUD.8884, was identified as a PRR candidate (Table 3), and is an ortholog of the *GRC*/*PTC1* gene 2 that is tied to schistosome resistance in *B. glabrata* (40). This gene shows a tandem duplication in African *Biomphalaria* species (tandem neighboring gene in *B. sudanica* is BSUD.8885), and this duplication event has occurred independently of a similar duplication seen in some *B. glabrata* haplotypes (Figure 7). A second shortlisted hyperdiverse protein, BSUD.20268 in contig 7608 (Table 3), could not be well described upon comparisons to other domains or proteins, perhaps due to its small (75 aa) size.

**Figure 7.**
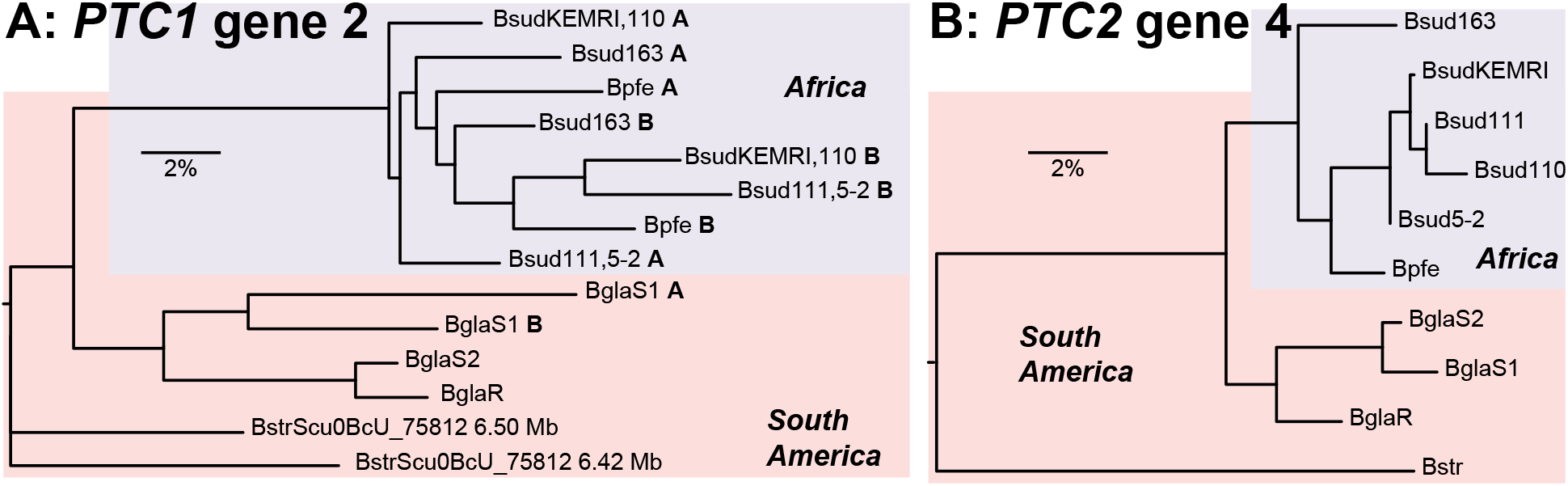
Allelic phylogenies of *Biomphalaria sudanica*, *B. pfeifferi*, and *B. glabrata*, rooted with *B. straminea*. (A) Polymorphic transmembrane cluster 1 (*PTC1*) gene 2 (i.e. *grctm2* (40)) shows a tandem duplication (see A and B duplicates for relevant taxa indicated in bold) in the African species (A=BSUD.8884 and B=BSUD.8885 for *B. sudanica*) that is clearly independent of similar duplications seen in *B. glabrata* haplotype S1 and *B. straminea*. (B) *PTC2* gene 4 (BSUD.3979 in *B. sudanica*) shows distinct haplotypes in each *Biomphalaria* species.

The ortholog to the *BgTLR* gene, BSUD.8256 (see *tlr* Figure 6), is also contained within LG6 in a highly diverse region of the *B. sudanica* genome, although the gene itself is relatively conserved within *B. sudanica* (0.1% coding nucleotide diversity). Additional immune-related genes on this linkage group are listed in Supplementary Table 17.

### Pathogen recognition receptor candidates within Linkage Group 10

Synteny between *B. glabrata* and *B. sudanica* across LG10 was highly conserved (Supplementary Figure 3B). Two proteins coded by neighboring genes located less than 1 Mb from the orthologous site of schistosome resistance marker *RADres1* (78), BSUD.4884 and BSUD.4885 (contig 2266), showed extremely high (∼5%) nucleotide diversity between the *B. sudanica* inbred lines (Table 3). The predicted function of BSUD.4884, an 86 aa protein partially matching a hypothetical protein recorded in *B. glabrata* (GenBank accession KAI8743980), could not be confidently characterized due to an absence of significant matches to orthologous proteins or domains (Supplementary Table 19). Comparisons with the annotated *B. glabrata* genome also revealed that the neighboring protein, BSUD.4885, had been erroneously truncated on the extracellular portion; the manually annotated gene codes for a 405 aa protein. The complete BSUD.4885 protein was predicted to contain both TMEM154 and Rifin domains (see (80)), features shared with some *PTC1* and/or *PTC2* genes (see BSUD.3980, BSUD.8873, BSUD.8874, BSUD.8876 and BSUD.8884, Table 3 and Supplementary Table 17). Remarkably, haplotype lineages of BSUD.4885 in *B. sudanica* also occur in *B. glabrata* (Figure 8), suggesting that either this is a shared polymorphism within the genus that has survived the colonization of Africa, or that there were originally two divergent tandem paralogs but one or the other is independently deleted in every genome since no genome appears to have two copies of this gene.

**Figure 8.**
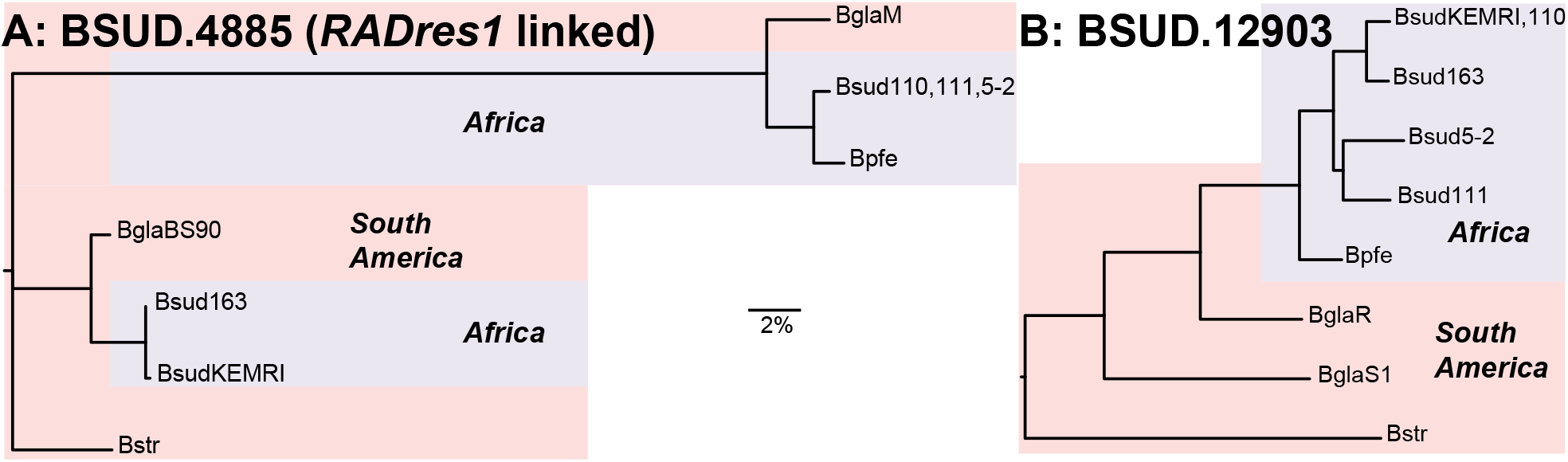
Phylogenetic trees generated using RAxML (81) of exemplar genes showing unusually high diversity in *Biomphalaria sudanica*, alongside orthologous alleles in *B. pfeifferi* and *B. glabrata*, and rooted with *B. straminea*. (A) BSUD.4885 (contig 2266, linkage group (LG) 10) is a gene with exceptionally high diversity, has a protein structure that shows similarities to genes in the polymorphic transmembrane cluster 1 (*PTC1*) (40) and *PTC2* (41), and is closely linked to a genomic marker, *RADres1*, that was previously shown to significantly influence schistosome resistance in *B. glabrata* 13-16-R1 strain (78). Phylogeny shows an apparent trans-species polymorphism of divergent haplotypes in African and South American snails. (B) BSUD.12903 (contig 499, LG16) is another pathogen recognition receptor candidate, in one of the most polymorphic contigs in the *B. sudanica* genome, predicted to contain C-type lectin, immunoglobulin, TMEM154 and alternatively expressed fibronectin III domains (see Figure 10).

### Pathogen recognition receptor candidates within Linkage Group 16

Linkage group 16 contains many of the most diverse genomic regions (Figure 9), such as contig 208, contig 499, contig 550, contig 580, and scaffold 3064, which include highly diverse individual genes (Supplementary Table 17). Six of the PRR candidates within LG16 originate from contig 208 (BSUD.3983, BSUD.4096, BSUD.4112, BSUD.3984, BSUD.4003, BSUD.4089, Table 3), the orthologous region to the *PTC*2 region in *B. glabrata* (41). As the orthologous *PTC2* genes are adjacent to these and many other diverse genes, several of which have single transmembrane domains and/or homology to *PTC2* genes (e.g. BSUD.4003), the resistance-associated *PTC2* region originally described in *B. glabrata* appears to be part of a much wider region of contig 208 (Figure 5C; Figure 6) with distinctive evolutionary and structural features. Furthermore, contigs neighboring contig 208 according to our inferred linkage map also contain single transmembrane domain proteins with homology to *PTC2* genes (BSUD.14425, BSUD.15084 and BSUD.23257 on contig 550, contig 580 and scaffold 3064, respectively).

**Figure 9.**
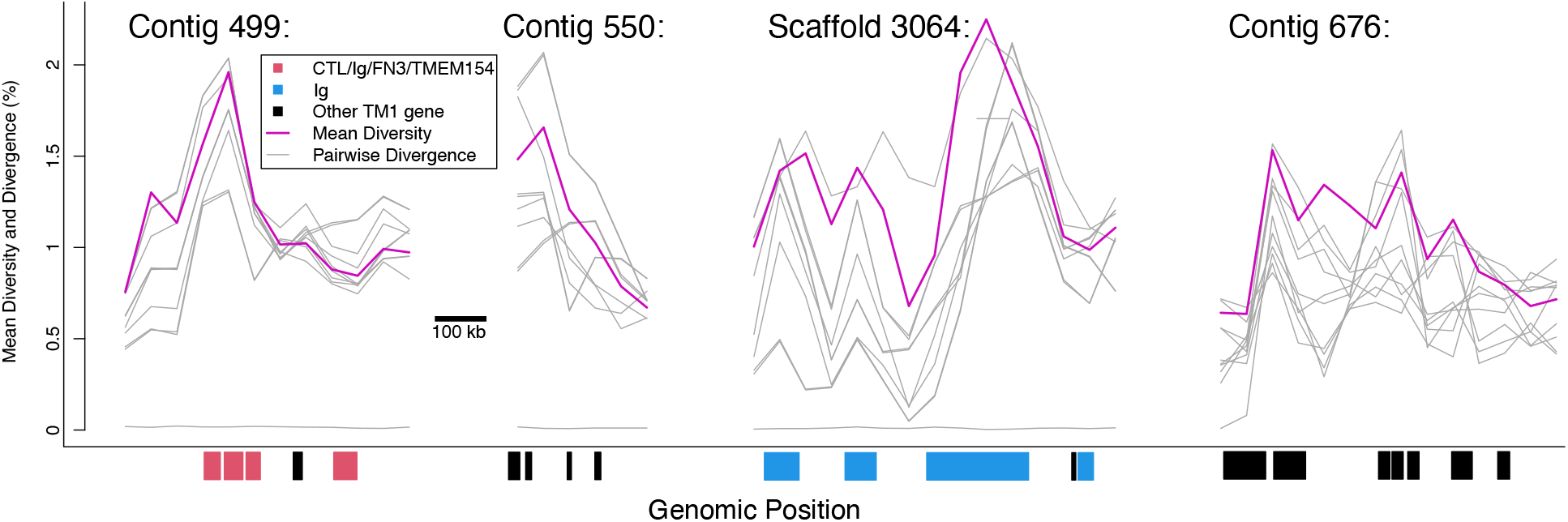
Mean nucleotide diversity (purple line) and pairwise divergence for each haplotype pair (grey lines) across portions of four contigs in LG16 (contig 499 300-850kb, contig 550 0-250 kb, scaffold 3064 2300-3000 kb) and LG5 (contig 676 150-850 kb) showing high diversity and clusters of transmembrane genes. All plotted genes (shown in red, blue and black) encode single-pass transmembrane proteins (TM1); other genes in these regions are not shown. Key functional domains potentially involved in pathogen recognition include C-type lectin (CTL), fibronectin type III (FN3) and immunoglobulin (Ig), and TMEM154, which is a membrane-spanning domain also found in several polymorphic transmembrane cluster 1 (*PTC1*) genes (40). Genes shown in red, including BSUD.12903 (Figure 10) all have at least three of these four functional domains, while genes shown in blue have only Ig.

PRR candidates BSUD.4112 (contig 208) and BSUD.15077 (contig 580) were predicted to be cell adhesion molecules since both contained a domain matching that of a protocadherin (Table 3; Supplementary Table 19). Two other PRR candidates, BSUD.12903 (contig 499) and BSUD.4089 (contig 208) (Table 3), both contain an extracellular C-type Lectin domain known to function in immune responses to pathogens. For BSUD.12903 at least, nonsynonymous diversity substantially exceeds synonymous diversity in the extracellular region of the protein where these predicted functional domains (C-type lectin and IgSF) were present (Figure 10). BSUD.12903 occurs within a cluster of proteins representing one of the most diverse regions of the genome, many of which contain combinations of C-type lectin, IgSF and FN3 domains (Supplementary Table 17). BSUD.4089 encodes a longer isoform (331 aa) with a single transmembrane domain and an extracellular C-type lectin domain, and a shorter isoform (67 aa) with just the secreted peptide signal (confirmed through InterProScan, see Supplementary File 3) and a partial match to the C-type lectin domain, which would result in a truncated domain.

**Figure 10.**
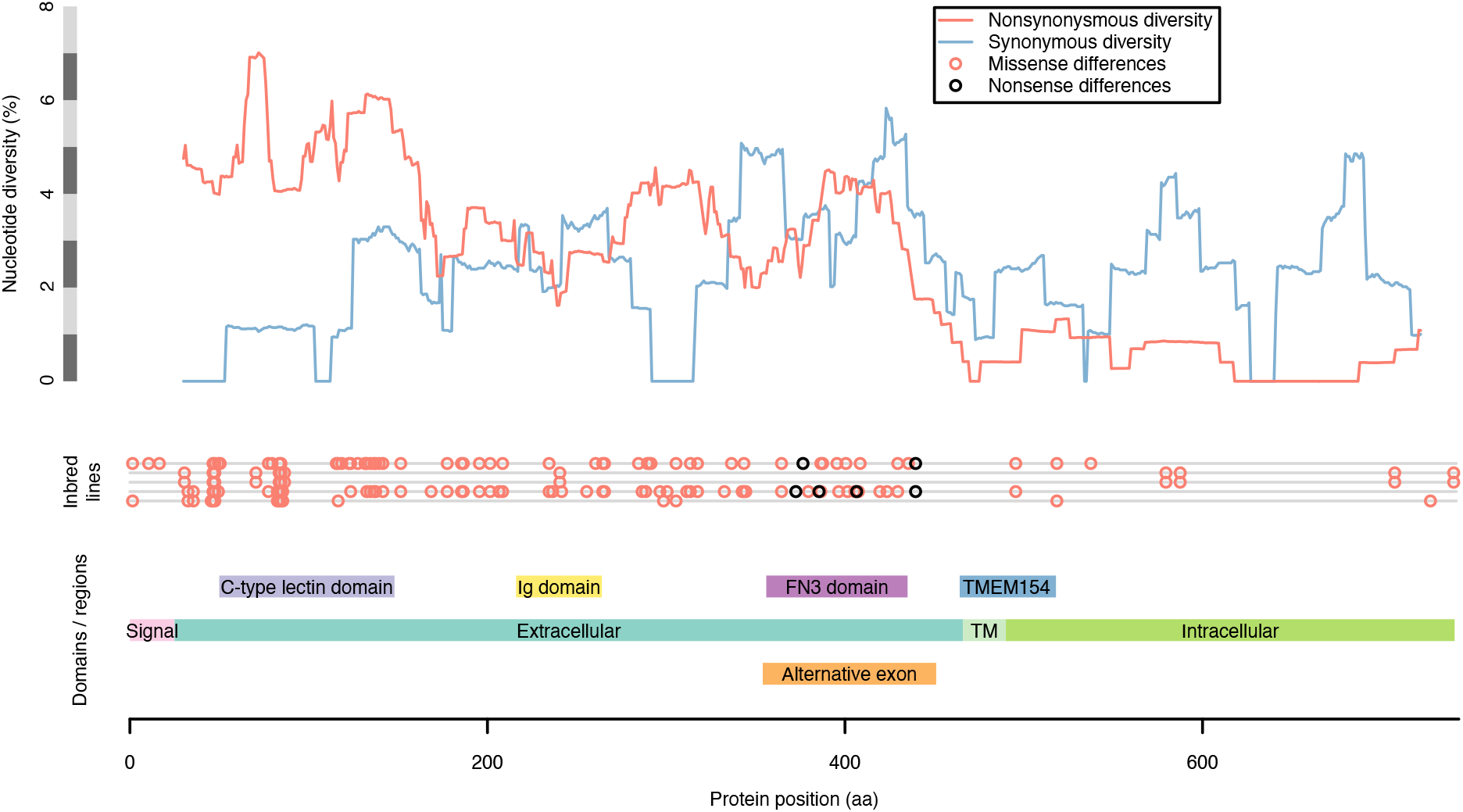
Detailed view of protein BSUD.12903 containing the alternatively expressed exon of 741 aa, which demonstrates that in the extracellular region nonsynonymous diversity greatly exceeds synonymous diversity (shown here in 50 aa sliding windows) in regions where functional domains potentially involved in pathogen recognition are present (C-type lectin, immunoglobulin, and fibronectin III (FN3) domains). Two inbred lines of *Biomphalaria sudanica* have multiple nonsense variants (stop codon or frameshift) in a single exon containing a FN3 domain, which is expressed in the *B. pfeifferi* ortholog (KAK0057508 (8)), suggesting that FN3 may not be expressed in all *B. sudanica* lines. The FN3 exon also occurs in the *B. glabrata* ortholog BGLB024560, where it is also variably either expressed (NCBI Accession XM_056010427) or excluded (NCBI Accession SRX8534561). Both FN3 and/or TMEM154 domains are present in *B. sudanica* genes BSUD.8884, BSUD.8874 and BSUD.8876 that are orthologous to *B. glabrata PTC1* region genes associated with schistosome resistance: *grctm2*, *grctm3* and *grctm4* (40).

A large inversion (∼15 Mb) between *B. sudanica* and *B. glabrata* iM line on LG16 appears to reverse parts of scaffold 3064, contig 208 and contig 550 relative to *B. glabrata* scaffolds BGM014 and BGM015 (Figure 6; Supplementary Figure 3C). BGM014 itself may be incorrectly assembled since it is split across *B. glabrata* linkage groups LG3 and LG16.

## Discussion

### A new genomic resource for vector biology

Long-read PacBio HiFi DNA and RNA sequencing, supplemented by Illumina short-read sequencing of RNA, was performed on an inbred line of *B. sudanica* sensu lato, originally collected from shoreline habitats of the Kisumu region of Lake Victoria. This sequencing approach and bioinformatic pipeline resulted in a well-supported genome annotation, which adds to the growing repository of *Biomphalaria* species genomes. *Biomphalaria sudanica* remains a “neglected vector” despite its established role in transmission of *S. mansoni* in lake and marsh habitats in the African Rift Valley, where schistosomiasis transmission is among the highest in the world. Our analyses focused on characterizing immune genes using a comparative framework with that of the better studied South American vector of schistosomiasis, *B. glabrata*, and another African vector, *B. pfeifferi*, for which genomic resources have recently been developed (8). We also developed a novel bioinformatic pipeline to characterize regions of marked intraspecific diversity as a mechanism to identify novel genes that may be involved in pathogen recognition and are under balancing selection. These analyses and resources will facilitate future work to explore further and uncover additional vector immune defense mechanisms against pathogens such as schistosomes, with direct relevance to public health in Africa where the majority of *S. mansoni* infections occur. The hope is that the scientific community will use these resources to support the development of novel tools for schistosome control, potentially including gene-drive manipulation of the snail vector (82), and development of surveillance tools, as is described as a priority in World Health Organization guidelines (83).

### Comparative analysis of immune genes among species

Based on the average number of substitutions per site, the genomes of *B. sudanica* and *B. pfeifferi* are approximately 5.1% divergent from each other, which is comparable to that of the isolates of *B. glabrata* compared here (between 3.3-8.7%) (see Figure 1). We found that *B. sudanica* possesses orthologous genes to all those previously identified from *B. glabrata* as having a potential function in immunity against schistosomes. However, the functions of these orthologs have yet to be verified for *B. sudanica*.

The number of complete variable IgSF and lectin domain-containing molecules (FREPs and CREPs) in *B. sudanica* was almost identical to that observed in *B. pfeifferi*, with both species having a higher number than observed in *B. glabrata* (Table 1). The largest and most divergent FREP family in *B. sudanica* is FREP12, two genes of which were not identified by the VIgL annotation pipeline but instead were discovered with the pipeline to identify highly diverse immune related proteins and PRRs. Within the African species, some FREP-like gene families were enriched in *B. sudanica* and underrepresented in *B. pfeifferi.* This association is interesting given the observation that *B. pfeifferi* is more susceptible to schistosome infections compared to *B. sudanica* (84,85); however the bulk of enriched FREPs in *B. sudanica* (FREPK1, FREPJ3, FREP5, FREPJ5, FREP12) are not known to play a role in schistosome resistance as based on studies of *B. glabrata* with comparable studies for African species yet to be undertaken. It is also apparent that FREP genes duplicate more readily in *B. sudanica*, resulting in more truncated FREPs than CREPs.

It should be acknowledged that the annotation and nomenclature of FREPs is not trivial. Diversification of FREPs by gene conversion, duplications, gene loss, exon shuffling, and other structural rearrangements, rather than the slow accumulation of mutations, complicates phylogenetic analysis and therefore classification and nomenclature of these genes (43). Due to the complex molecular evolution of FREPs, the *B. sudanica* genome contains a diverse and potentially variable suite of FREP sequences as seen in *B. glabrata* (86), independent of nucleotide diversity at any particular gene. Further investigation into these hypervariable genes is necessary to fully describe the molecular mechanisms that drive their diversity and interactions with pathogens.

Elevated diversity at candidate innate immune gene regions, *PTC1* and *PTC2*, in both *B. sudanica* and *B. glabrata* appears to be independently generated, since the genes remain reciprocally monophyletic (Figure 7), which suggests ongoing selection favoring novel diversity at these loci. Occasional transspecies polymorphisms, including one apparent at BSUD.4885 (Figure 8), are suggestive of a long-term balancing selection, strong enough to overcome what was likely a narrow population bottleneck during the colonization of Africa.

### Intraspecific genomic diversity

Hyperdiverse protein coding genes in *B. sudanica* that were shortlisted as potential PRRs under balancing selection were enriched in molecules that have been associated with various aspects of immune functions in other organisms. The hyperdiverse transmembrane proteins included G-protein coupled receptors (GPCRs), TLRs, cluster of differentiation (CD) and cell adhesion molecules as well as proteins containing functional domains such as C-type lectins, FN3, IgSF, transient receptor potential (TRP) channels and many potentially immune related domains that await further immunological characterization. These often show high extracellular nucleotide non-synonymous diversity which overlaps with functional binding domains (see Figure 10). While we have focused on candidate PPRs with high nucleotide diversity, we do not mean to discount the importance of conserved genes, or the between-paralog diversity found in multi-gene families. In particular, genes with well-studied immune roles in *B. glabrata*, such as FREPs (42) and components of the oxidative burst pathway (87), undoubtably also play essential roles in *B. sudanica* immunity regardless of their allelic diversity.

The most diverse candidate protein (Table 3; BSUD.20937) has a single transmembrane domain and a predicted tumor necrosis factor (TNF) domain. TNFs in invertebrates are far less characterized than those in mammals; however, recent work in mollusks and crustaceans indicate that just like in vertebrates, they have multiple roles in innate immunity and function in activation of antimicrobial peptides, apoptosis and phagocytosis of hemocytes, and activation of immune related enzymes such as lysozyme and phenoloxidase (88–90). We could not detect this gene in two inbred lines (Bs163 and Bs5-2), presumably due to either deletion or extreme sequence divergence, and thus our diversity measurement may even be an underestimate. The high diversity of BSUD.20937 between *B. sudanica* inbred lines, and that of the other hyperdiverse transmembrane genes selected in our bioinformatic pipeline, suggests that these may well be PRR genes under balancing selection driven by variability in ligand binding sites that interact with pathogens.

Regarding genome-wide patterns of diversity, the identified high diversity windows were clustered on three linkage groups (6, 10, and 16), and near candidate immune genes. Linkage group 6 contains a cluster of highly diverse windows as well as several candidate immune genes including *PTC1*, *cat*, *tlr*, *sod1*, and *prx4*. Linkage group 10 also harbors diverse windows as well as resistance region *RADres*. The highest diversity genomic windows occur on linkage group 16, site of the highly diverse gene cluster *PTC*2 which we show is imbedded within a larger region of diversity (Figure 4 and 6) that shows chromosomal rearrangement between *B. sudanica* and *B. glabrata* (Supplementary Figure 3C). Such structural rearrangements may be enriched along with single-nucleotide polymorphisms in hyperdiverse regions like *PTC2* and may even help to maintain diversity if they are segregating within species; however we also note that the 8.5 Mb contig 208 is the largest contig in our Bs111 assembly and thus it afforded the greatest power to observe large rearrangements that may be undetectable elsewhere in the genome. Immune-relevant genes may often be linked within chromosomal regions with distinct evolutionary dynamics, including perhaps the maintenance of elevated adaptive diversity, as also noted for *B. glabrata* (91,92).

### Molecular evolutionary legacy of colonization and adaptation in Africa

*Biomphalaria* originated in South America, but at least one lineage crossed the Atlantic based on previous estimates approximately 1.8 and 5 MYA (8–11) and diversified into the contemporary African species. Such a transcontinental colonization, whenever it did take place, is striking for a freshwater gastropod of limited vagility.

Since the transcontinental colonization of *Biomphalaria* is predicted as sufficiently evolutionarily recent, its history can be explored more easily with the growing collection of *Biomphalaria* genomes. Despite the divergence between *B. glabrata* and the African species being between 10.5-10.8% (Figure 1), thousands of orthologs could be aligned easily across these species and without saturation of neutral sites. These highly orthologous genomes allowed us to examine the expansion and contraction of gene families, with particular interest being paid to the genomic consequences of migration and adaptation of *Biomphalaria* to Africa. Originally, we hypothesized that immune genes of this African *Biomphalaria* ancestor would be substantially expanded as it encountered several new pathogens as it colonized Africa from South America; however, the analysis revealed no significantly expanded immunity-related genes. In fact, the contraction of genes outweighed expansions in the common ancestor of *B. sudanica* and *B. pfeifferi* compared to the South American congeners, suggesting an overall simplification of the genome in *B. sudanica* and *B. pfeifferi*. Perhaps this gene family contraction in African *Biomphalaria* was a maladaptive result of a founder effect, given that the ancestor of African *Biomphalaria* must have descended from a small number of colonizers, and/or perhaps many genes were freely lost without fitness consequences in the absence of the neotropical pathogens to which they were adapted. Despite this proteome simplification, the estimated genome size of *B. sudanica* was ∼73 Mb larger than the genome of outcrossing *B. glabrata*, and also ∼173 Mb larger than that of *B. pfeifferi*, which is hypothesized to have lost genes associated with mating given its reliance on selfing (8). This size discrepancy could be a technical artifact from divergent allelic *B. sudanica* haplotypes getting assembled as separate contigs, perhaps due to the higher heterozygosity in our Bs111 after only three generations of selfing, compared to naturally inbred *B. pfeifferi*, which is a preferentially selfing snail (93,94), and the long-term inbred lines of *B. glabrata*. Alternatively, if the increased genome size of *B. sudanica* is biologically real, it could represent gene duplications and insertions not picked up in our current analysis, which may have led to novel gene functions and expression mechanisms favored in its African habitat. The mechanism could also be selectively neutral: the high repeat content of all *Biomphalaria* genomes indicates that total genome size can vary considerably with little direct effect on the content of coding genes.

As confirmed through our annotation of RNAs, a common feature in the two African species is the presence of tRNA-SeC, demonstrating the ability of *B. sudanica*, like its sister species *B. pfeifferi* (8), to synthesize polypeptides containing selenocysteine (53,54). Although selenoproteins are present in many gastropod lineages (95), tRNA-SeC has not been identified in *B. glabrata* (8), suggesting that the capacity to produce selenocysteinyl has been gained in the African *Biomphalaria* species, or lost across South American lineages of *B. glabrata*. With the growing repository of genome information for species basal to *Biomphalaria*, it will be possible to investigate this further. Selenocysteine-containing proteins, i.e. selenoproteins, are proposed to be involved in a wide range of bioactive processes in other invertebrates (96).

Interestingly, regarding rRNA genes, far fewer were predicted in *B. sudanica* compared to *B. pfeifferi*, although this is likely due to misassembly of the highly repetitive tandem repeats that rRNA genes are generally present in, causing near identical rRNA copies to be collapsed into single copies, rather than biological differences.

### Transcriptomic analysis illuminates unusual mitochondrial transcription processes

Molluscan mitochondrial genomes are unusual in terms of their genomic features and transcriptomic processes (56,58), and the latter has not been explored for African *Biomphalaria*. In *B. sudanica*, mitochondrial genes that were not separated by tRNA genes, namely *nad6*, *nad5*, and *nad1*, were highly represented by polycistronic mRNA transcripts with no clear trimming points. Arrangement and transcription processes of these genes therefore appear similar in other invertebrate and molluscan species (58,97,98). In addition, *nad4l* lacked read coverage, with the few transcripts that were observed being polycistronic with *cob*. Lack of expression in key genes of the respiratory chain, such as the NADH dehydrogenase, is not unheard of within mollusks (99), but regardless the sharp increase on read coverage at the start of *cob*, suggests this is also processed by endonucleases.

Our transcript for *B. sudanica* across mitochondrial genes *atp6* and *atp8*, which in other species form a single mRNA (97), are punctuated by trnN in *B. sudanica* as in *B. glabrata* (59). However, the bulk of the recovered reads belong to the pre-mRNAs. Considering their polycistronic status in other organisms, it is possible that translation could happen before the pre-mRNA is fully resolved, which would add another layer of complexity to molluscan mitochondrial genomes in that these genes are likely separated downstream into proteins through initiation on the ribosome (as reviewed in (58)). Additionally, given that *atp8* has been found under relaxed selective pressures and a high degree of variation in both *Biomphalaria* and *Bulinus* species (55,100), it may play a less important role in the mitochondrial function within these planorbids.

### Conclusions

The species *B. sudanica* is remarkable for two principal reasons: its dynamic evolutionary history involving recent colonization and diversification in Africa, and its tragic impact on public health as a schistosome vector. The genomic data presented here illuminate both aspects. Our observations include expansions and contractions of gene families in African snails, as well as numerous genomic regions of high diversity that contain genes that may play a role in host-pathogen coevolution and their vectorial capacity for parasites such as schistosomes. In combination with increasing genomic resources from other *Biomphalaria* isolates, this work will facilitate an enhanced understanding of the biology of these snails and future mechanisms for curbing transmission of schistosomiasis.

## Methods

### Ethical approval and permitting information

This project was undertaken following approval from the relevant bodies, including Kenya Medical Research Institute (KEMRI) Scientific Review Unit (permit # KEMRI/RES/7/3/1), Kenya’s National Commission for Science, Technology, and Innovation (permit # NACOSTI/P/22/148/39 and NACOSTI/P/15/9609/4270), Kenya Wildlife Service (permit # WRTI-0136-02-22 and # 0004754), and National Environment, Management Authority (permit # NEMA/AGR/159/2022 and # NEMA/AGR/46/2014) (Registration # 0178).

### PacBio whole genome sequencing and assembly of Biomphalaria sudanica line 111

High molecular weight DNA was isolated from the headfoot tissue of a single adult (>8 mm in shell diameter) *Biomphalaria sudanica* snail from the inbred line “111” (Bs111) following the Qiagen Blood & Tissue kit (Qiagen, MD, USA), the only modification to standard kit protocol being the overnight lysis of tissue at 37°C. The inbred line, Bs111, was developed from snails originally collected from the Kisumu region of Lake Victoria, Kenya in 2012 and bred via selfing for 3 generations before the line was expanded through sibling mating and gDNA extracted for sequencing. PacBio high-fidelity (HiFi) circular consensus long-read sequencing (101) was carried out using two SMRT cells of a PacBio Sequel II at the University of Oregon. PacBio ccs reads (raw reads available at NCBI Sequence Read Archive accession: In prep.) were assembled using Flye (102), settings: flye --pacbio-hifi Bsud111_ccs.fasta -g 1g -i 0 -t 16 -o flye/Bsud111. Basic statistics of the genome assembly were calculated in seqkits v0.16.1(103).

### Illumina whole genome sequencing, alignment and genotype calling of four genetic lines of B. sudanica

Short-read Illumina data for four additional *B. sudanica* genetic lines Bs5-2, Bs110, Bs163 and BsKEMRI, all originally collected between 2010-2012 from the Kisumu region of Lake Victoria, Kenya, were also produced. Inbred lines were generated via selfing in isolation for 3 generations, after which, the lines were expanded through sibling mating. The BsKEMRI line snail was not subject to intentional inbreeding but was maintained in the laboratory for ∼10 years (2010–2020) before gDNA was extracted for sequencing. High molecular weight DNA was extracted as described above. Illumina pair-ended reads (150 bp) were generated using the HiSeq 3000 at the Center for Quantitative Life Sciences at Oregon State University to obtain 15-20x coverage for each sample. The FASTQ files had adapters removed with Cutadapt v1.15 (104) and then reads were trimmed with Trimmomatic v0.30 (45) with options: LEADING:20 TRAILING:20 SLIDINGWINDOW:5:20 MINLEN:50, and aligned using BWA version 0.7.12 (105) using command: bwa mem -P -M -t 4, with the PacBio assembly of *Bs*111 (see above). Aligned genome data and raw reads are available in archive NCBI Sequence Read Archive (Accession: In prep.).

Genotypes for each snail genome were called against the Bs111 reference genome in variant call format (vcf) produced using BCFtools v1.9 (106). Basic statistics of the alignments to the genome were calculated using vcftools v0.1.17 (107).

### Nuclear genome annotation of B. sudanica ‘111’ from long- and short-read RNA sequence data

The bioinformatic pipeline for the annotation of *B. sudanica* was performed using a custom script (see: github.com/J-Calvelo/Annotation-Biomphalaria-sudanica). To obtain the best possible genome annotation for *B. sudanica*, combined long- and short-read RNA-seq data was used. RNA for long- and short-read sequencing was obtained from multiple samples of pooled Bs111 snails during two developmental stages (juvenile and adult) or under different stressors (heat stress or exposure to *S. mansoni* NMRI strain from the NIH-NIAID Schistosomiasis Resource Centre (108)). Full details of samples used, and RNA extraction protocol and pooling are contained in the Supplementary Material (Supplementary File 6). The equal mass pool of eight samples was processed for sequencing using the standard IsoSeq protocol (PacBio protocol: PN 101-763-800 v2). The NEBNext Single Cell/Low Input cDNA Synthesis and Amplification Module kit (New England BioLabs Inc. MA, USA) was used following manufacturers protocols for cDNA synthesis and amplification from 300 ng of RNA, and the library was generated using the SMRTbell Express Template Prep Kit 2.0 (Pacific Biosciences, CA, USA). A Pronex bead purification (Promega, WI, USA) was performed on the amplified cDNA product following a size selection of =2kb transcripts. The final cDNA library was sequenced on one SMRT cell on the PacBio Sequel II at GC3F, University of Oregon.

Short-read Illumina sequences of RNA were obtained from two samples, one of a single unchallenged Bs111 and the second a pool of three Bs111 adult snails that had been challenged to *S. mansoni* miracidia 24 hours prior (Supplementary File 6). Library preparation and directional mRNA sequencing were performed at Novogene Corporation Inc. (CA, USA). Library preparation was performed using the NEBNext Ultra II Directional RNA Library Prep Kit (New England BioLabs Inc. MA, USA) for strand-specific Illumina libraries. Libraries were then sequenced on the NovaSeq 6000 platform (Illumina, CA, USA) and 150 bp pair-ended reads were generated to provide ∼12 G of sequence data.

### Bioinformatic analysis of long- and short-read RNA-seq data

Quality of the long-read data was first assessed with longQC software (44). Then, reads were processed with Lima (github.com/PacificBiosciences/barcoding), with the options “--css and --dump-clips”, and isoseq3 refine tool (github.com/PacificBiosciences/IsoSeq), with the option “- -require-polya”, to recover all complete sequenced transcripts (both 5’ and 3’ adapters and poly-A tail should be present to consider a transcript as complete). Complete transcripts were then mapped to the assembled genome with minimap2 (46,47) with options “-ax splice:hq -uf”. In parallel, short-read quality was verified using FASTQC software (109) and Illumina’s adapter sequences were removed along with low-quality bases using Trimommatic (45), with options “ILLUMINACLIP:TruSeq3-PE-2.fa:2:30:10 SLIDINGWINDOW:5:20 MINLEN:25”. The cleaned reads were then mapped to the genome using STAR (48) with options -- genomeChrBinNbits 13 --genomeSAindexNbases 13, as suggested by the authors for the genome assembly. Alignment quality was measured with Bamtools v2.5.2 (110). Transcripts were then characterized using both types of reads with StringTie2 v2.2.1 in “--mix” mode (49) plus the “-- conservative” option. Transcriptome completeness was evaluated with BUSCO (52) using Mollusca as the reference. Nuclear coding sequences longer than 150 bases (50 amino acids) were predicted with the utility tool TransDecoder.Predict v5.5 of the Trinity platform (50), with the option --single_best_only, making use of homology information from UniProt’s Swiss-Prot (111) and Pfam v35 protein domains (112). Similarity searches were done with BLASTp, with the “- evalue 1e-5” parameter (113) and hmmscan program with default parameters (114), for both databases, respectively.

### Functional annotation

Predicted proteins determined by nuclear coding sequences of TransDecoder.Predict v5.5 (50) were functionally annotated using eggNOG-mapper v2.0 (database downloaded 7/22/2022 (70,71)) for Gene Ontology (GO Terms) assignation based on curated orthogroups, and InterProScan v5.56-89.0 (51) for protein identification based on domain prediction.

### Repeat identification and masking of nuclear genome

Repetitive elements were annotated and masked using Earl Grey v 1.2 (115) pipeline. In short, repeat elements were identified with RepeatMasker v4.1.2 (116) with the “-norna, -nolow, -s” option and using Dfam v3.6 (117) curated database for the group Eumetazoa as reference. Then, RepeatModeler v2.0.3 (118), RECON v1.08 (119) and RepeatScout v1.0.5 (120) generated a *de-novo* repeat library that was refined by a “BLAST, Extract and Extend” process to combine fragmented detections into a single repeat candidate. These repeats were finally used by a second run of RepeatMasker v4.1.2 (116) to identify novel repeat elements. Summarizing plots were generated with ggplot2 (121).

### Non-coding RNA annotation

Transfer RNAs (tRNAs) and ribosomal genes (rRNA) were predicted using tRNAscan-SE v2.0.9 (122) and Barrnap v0.9 (github.com/tseemann/barrnap), respectively, options for both configured for eukaryotes.

### Mitochondrial Genome Annotation

The contig encoding the mitochondrial genome was identified by BLAST searches of the genome assembly (-perc_identity 80) (113) against the known mitochondrial protein coding genes from *B. glabrata* (NC_005439.1 (59)). One retrieved, the DNA sequence (contig 7934) independently annotated the retrieved DNA sequence with MITOS2 (60) web server (mitos2.bioinf.uni-leipzig.de/index.py: accessed July 2022). Genes not automatically annotated by mitos2 (*nad4l* and trnK, see Results) were localized with BLASTn (113) searches with respective *B. glabrata* mitochondrial genes (BLASTn with options -task blastn and -task blastn-short for *nad4l* and trnK, respectively).

RNA PacBio reads were mapped to the mitochondrial genome following the same procedure as for the nuclear genome annotation (see above). Transcript alignments to each mitochondrial gene were inspected on Integrative Genomics Viewer (IGV) (123) to manually improve gene annotation making the assumptions that a) translation started on the first viable start codon and b) stop codons were complete unless read coverage suggested a premature RNA end with a truncated stop codon, similar to the assumptions described previously (58). Reads belonging to intermediary pre-mRNA were counted with htseq-count (124) using the options “-- nonunique=all --samout” and custom scripts (see: github.com/J-Calvelo/Annotation-Biomphalaria-sudanica). Using the pre-mRNA read data, a schematic representation of the post-transcriptional trimming/modification processes of the primary transcript was then manually generated using Inkscape (inkscape.org).

In line with other already published mitochondrial genomes for *B. sudanica* (NCBI RefSeq: NC_038060.1 (55)) and other *Biomphalaria* species (NCBI RefSeq: NC_038061.1 and NC_038059.1, GenBank Accession: MG431965.1 (55)) the start coordinate for the mitochondrial genome was set to the first codon of the ND5 gene (Supplementary Figure 4). However, for manual inspection of read alignments on IGV, a custom origin set to position 7222 (between two tRNA molecules, tRNA-s1 and tRNA-s2, lacking both gene predictions and read coverage from the PacBio RNA sequence dataset) was used.

### Gene family analysis and evolutionary position of B. sudanica

A general study of the evolutionary dynamics (copy number and selection pressure) of *Biomphalaria* protein families was carried out based on the longest proteins predicted for each gene, in combination with 3 *B. glabrata* strains: BB02 (NCBI RefSeq: GCF_000457365.2 (5)) iBS90 and iM (GenBank Accession: GCA_025434165.1 and GCA_025434175.1 (6)), and the species *B. pfeifferi* (NCBI BioProject Accession: GCA_030265305.1 (8)), *B. straminea* (GenBank Accession: GCA_021533235.1 (7)), with the planorbid snail *Bulinus truncatus* (GenBank Accession: GCA_021962125.1 (125)), and the Plakobranchidae *Elysia marginata* (GenBank Accession: GCA_019649035.1 (126)) as an outgroup. With the goal of homogenizing criteria and avoid discrepancies between the reported mRNA sequence and their proteins, protein sequences taken from other works were predicted from the reported cDNA using getorfs from the EMBOSS v6.6.0.0 package (127). The longest ORF among all isoforms (min size 150 bases) of each gene were selected, ties were broken by picking the most upstream candidates for each gene. Orthology relationships were estimated in HOGs with Orthofinder v2.5.4 (67) and treated as putative protein families. A species tree was generated in Orthofinder using the Species Tree from All Genes (STAG) algorithm (67,74), to confirm that the topology of the tree matches those previously reported for these *Biomphalaria* species (9,128).

Significant expansions/contractions above the background were identified with CAFE 5 (68). The Enriched GO terms among significantly expanded/contracted HOGs from CAFE 5 were determined using the topGO R package v2.48.0 (72), with the Weight01 algorithm, node size=10 and statistical test of Fisher (significant p-value ≤ 0.05). To this end, GO Terms were assigned to each species’ genes as described above, and those annotations were assigned to each HOG. Summary plots of gene lists were produced in REVIGO to allow further interpretation (73).

### Cellular location of proteins: secreted, mitochondrial translocated and transmembrane proteins

Location signals for exportation or mitochondrial translocation of proteins were predicted with SignalP v6.0 (61) and TargetP v2.0 (62), respectively. Isoforms with negative results for both tools were additionally analyzed with SecretomeP v1.0 (63) to identify additional secreted proteins based on their biochemical characteristics, that is, proteins exported through an alternative route or with location signals missed due to annotation errors. For all three analyses only hits with a probability/score above 0.95 were considered significant. Lastly, TMDs were predicted with DeepTMHMM v1.0.13 (129).

### VIgL (FREP/CREP/GREP) identification and analysis

Based on the presence or absence of known domains and features of FREPs, CREPs and GREPs, as reviewed previously (43), these VIgLs were identified following these criteria: 1) Evidence of secretion (see section: *Cellular location of proteins*), 2) presence of IgSF domains, which were identified using hmmsearch v3.3.2 (75) using the domain profiles generated previously (43), and 3) evidence of their respective 3’ terminal domain in the InterPro annotation: FBD (IPR002181, IPR014716, IPR020837, IPR036056), C-lectin (IDs: IPR001304, IPR016186 IPR016187, IPR018378) or Galectin (IPR001079, IPR015533, IPR044156, IPR000922, IPR043159). For a summary of the expected hits for each family, see Dheilly et al. (76). Lastly, a BLAST search between *B. sudanica* transcriptome and the *B. glabrata* FREP, CREP, and GREP genes reported by Lu et al. (43) and an additional 17 reference sequences (13 FREP and 4 CREP from NCBI (Supplementary Table 15) were performed.

Full FREP, CREP, and GREP candidates were defined as genes with a SP, at least one IGsF, and their respective 3’ terminal signature domain (FBD, C-lectin, or Galectin domain respectively). Classification into subfamilies was carried out according to their phylogenetic relationship with the reference sequences. For each candidate gene the longest isoform reported for each gene and reported as a full candidate for each family were aligned in MAFFT v7.310 (130) and positions with more than 20% gaps removed with trimal v1.4.rev22 (131) (options -gt 0.8 -st 0). Then their phylogenetic relationships were estimated by maximum likelihood with IQ-TREE v.2.2.0.3 (132). The best substitution model was selected by ModelFinder (133) using the Bayesian Information Criterion, by setting “-m MFP” as part of IQ-TREE options. The DNA substitution models VT+I+G4 and JTT+F+R6 models were selected as the best fitting for the CREP and FREP sequence alignments, respectively. Node support was estimated with 1000 replicates of non-parametric bootstrap using IQ-TREE options “-b 1000”. Trees were re-rooted at the midpoint and inspected in iTOL v6 (134).

### Identification of previously identified schistosome resistance genes

Several candidate immune genes have been identified in *B. glabrata* that could be involved in the response of snails to *S. mansoni*, inferring resistance (Supplementary Table 2). To test whether polymorphisms were shared within *B. sudanica* and between species of *Biomphalaria* in some of these candidate PRRs and immune genes that show the strongest support (BSUD.12903, *PTC1*, *PTC2*, *RADres*), phylogenies were generated using RAxML with -m GTRCAT (Stamatakis, 2006).

### Assessment of highly diverse genes and genome regions for novel pathogen recognition receptors

Intraspecific genomic diversity between the five *B. sudanica* inbred lines was determined by interrogating polymorphisms between the 5 inbred line vcf file (see above) using statistical measures in vcftools v0.1.17 (107). High-diversity genes were determined by calculating nucleotide diversity across both the entire gene (including noncoding regions such as introns and UTRs) and in coding regions only, determined by the Bs111 nuclear genome annotation, of which the top 1% were selected for further analysis (Supplementary Figure 5). In addition, we identified genes occurring in or near diverse windows, as follows. We calculated nucleotide diversity across the genome in sliding windows of 10 kb (starting every 2.5 kb), 30 kb, (starting every 7.5 kb) or 100 kb (starting every 25 kb), and identified the top 0.1% of 10 kb windows, top 0.3% of 30 kb windows, and top 1% of 100 kb windows. We then identified all genes that occur within (or overlap partially with) 100 kb of the midpoint of any of these diverse windows.

For all unique genes identified following these criteria, the largest protein-coding amino acid sequence for each gene was summarized from the annotation (Supplementary File 7). Where available (those with matches), transcripts of coding regions from each gene (transcript determined from Bs111 RNAseq data) were matched with their annotated UniProt description (uniprot.org/, database as of May 25^th^, 2022) and InterPro v5.56-89.0 description (51) (Supplementary Table 19 and Supplementary Table 20). All amino acid sequences were further characterized by searching for orthologous proteins using BLASTp (113) on the NCBI protein database (ncbi.nlm.nih.gov/protein, database as of October 1^st^, 2022) (Supplementary Table 21). An e-value cut-off of 1e-50 was used for amino acid sequence matches. The proteins producing the most significant alignments (based on the lowest e-value and highest percentage identity from BLASTp), and those predicted using UniProt and Pfam to the query amino acid sequence were recorded and used to determine each peptide’s function/biological process and key protein domains if this information was available and informative.

Based on the annotated protein, functional domain presence and structure (i.e. presence of TMDs), each protein was then placed in immune and non-immune gene related categories, such as in similar analyses (135,136), and placed under the broader groups of 1. non-immune function suspected, 2. immune-related function, 3. potentially immune-related function, 4. unknown protein function but containing TMD(s), 5. unknown protein function because sufficient information could not be obtained and without TMD.

To create a shortlist (n=20) of candidate PRRs under balancing selection, proteins with the highest nucleotide diversity, and categorized as either in group 2, 3 or 4 that also contained at least one transmembrane domain were shortlisted. The genes and in some cases the genomic regions surrounding these were assessed in more detail, including assessing positions of functional domains and intra/extracellular regions relative to nucleotide diversity (Supplementary Figure 5).

To determine if any enriched GO terms, particularly those associated with immunity, were amongst the proteins identified as potential PRRs, topGO v2.48.0 (72) in R v4.3.1 (137) with the Weight01 algorithm, node size=10 and statistical test of Fisher (significant p-value ≤ 0.05) was used. The test was performed by comparing the *B. sudanica* whole genome gene family composition to three separate lists of the highly diverse genes identified: 1) all 1047 highly diverse genes identified in the *B. sudanica* genome (as listed Supplementary Table 17), 2) 245 genes classified as immune suspected genes from the highly diverse genes (group 2 in Supplementary Table 17), and 3) 242 genes shortlisted as the protein containing a transmembrane domain (TMD) and categorized in group 2, 3 (potential role in innate immunity) or 4 (an unknown function but contained TMD(s)) (Supplementary Table 17).

### Inferring linkage groups and synteny with *Biomphalaria glabrata* linkage groups

We compared synteny and orthology with *B. glabrata* using the iM assembly (6). We defined orthologous genes as reciprocal best BLASTp hits of protein sequences. We defined orthologous contigs/scaffolds as those sharing the most orthologous genes. For *B. sudanica* contigs/scaffolds that were orthologous to scaffolds mapped in the 18 *B. glabrata* iM LGs, we assigned them to 18 LGs corresponding to the linkage groups and ordered them to mirror the orthologous order. A single iM scaffold (BGM014) is split between linkage groups, so we considered the distinctly-mapped portions of it separately. For linkage groups of particular interest, we generated dot plots of sequence similarity based on our previous approach (138). We used these to refine the order of contigs/scaffolds, to identify contigs/scaffolds with little sequence similarity despite possessing nominally orthologous genes, and to characterize inversions and other chromosomal rearrangements.

## Declarations

The authors have nothing to declare.

## Ethics approval and consent to participate

Not applicable

## Consent for publication

Not applicable

## Availability of data and materials

The datasets generated and/or analyzed during the current study are available in the NCBI BioProject repository, accession number In prep.

The bioinformatic pipeline for the genome annotation is available at github.com/J-Calvelo/Annotation-Biomphalaria-sudanica.

## Authors’ contributions

TP, JCal., JAT, SRB and MLS collected, analyzed and interpreted the majority of the data contributing to this manuscript. TP, JCal., JAT, AI and MLS were major contributors to writing the initial manuscript. JCal. developed the bioinformatic pipeline for the annotation of the genome. JAT developed and conceptualized the bioinformatic pipelines for the analysis and description of hyperdiverse genes under balancing selection. RB and JCay. collected and analyzed data regarding the hyperdiverse immune genes in the genome. SRB and MSB generated and assembled/aligned whole genome sequence data from the inbred lines and assisted in generating transcriptome sequence data. MLS, TP, JMS, assisted in the maintenance and development of inbred snail lines. FGH, GO, FR, KA, BM, MOdh. and MOdi. assisted in manuscript preparation and editing. MOdi assisted in project planning. ESL, MRL and LL assisted in the comparative work with African *Biomphalaria* species, and LL assisted with the FREPs identification pipeline and analysis. MLS and MOdi. were responsible for funding acquisition to complete this project. All authors reviewed and approved the final manuscript.

## Acknowledgements

The NMRI schistosomes were provided by the NIAID Schistosomiasis Resource Center of the Biomedical Research Institute (Rockville, MD) through NIH-NIAID Contract HHSN272201700014I. We also acknowledge Hannah Tavalire for her work in developing the inbred lines of *B. sudanica*.

## Supplementary Material

### Supplementary Figures

**Supplementary Figure 1.**
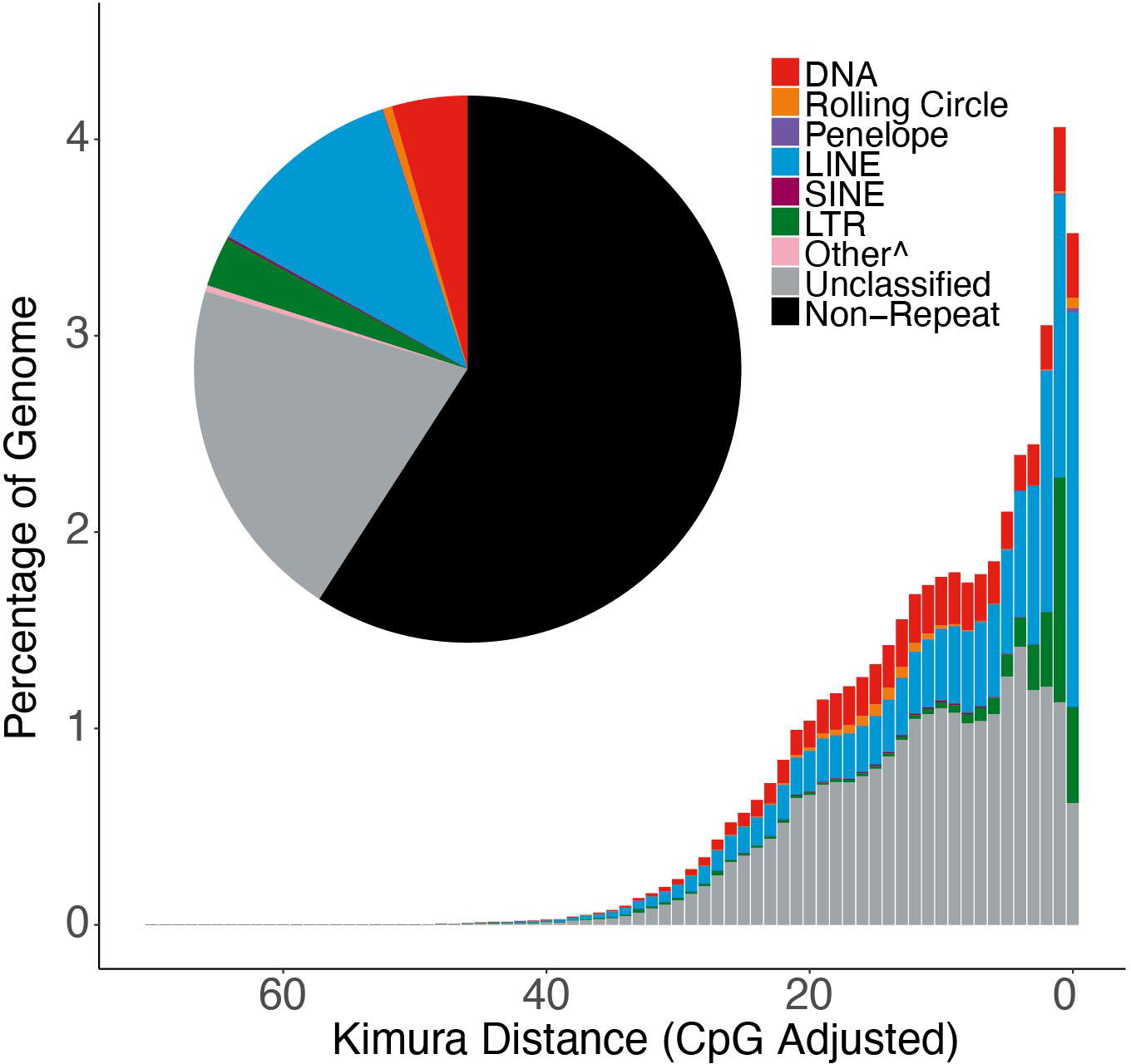
Repetitive elements in the *Biomphalaria sudanica* genome. ^ “Other” indicates a Simple Repeat, Microsatellite, RNA (.jpg file).

**Supplementary Figure 2.**
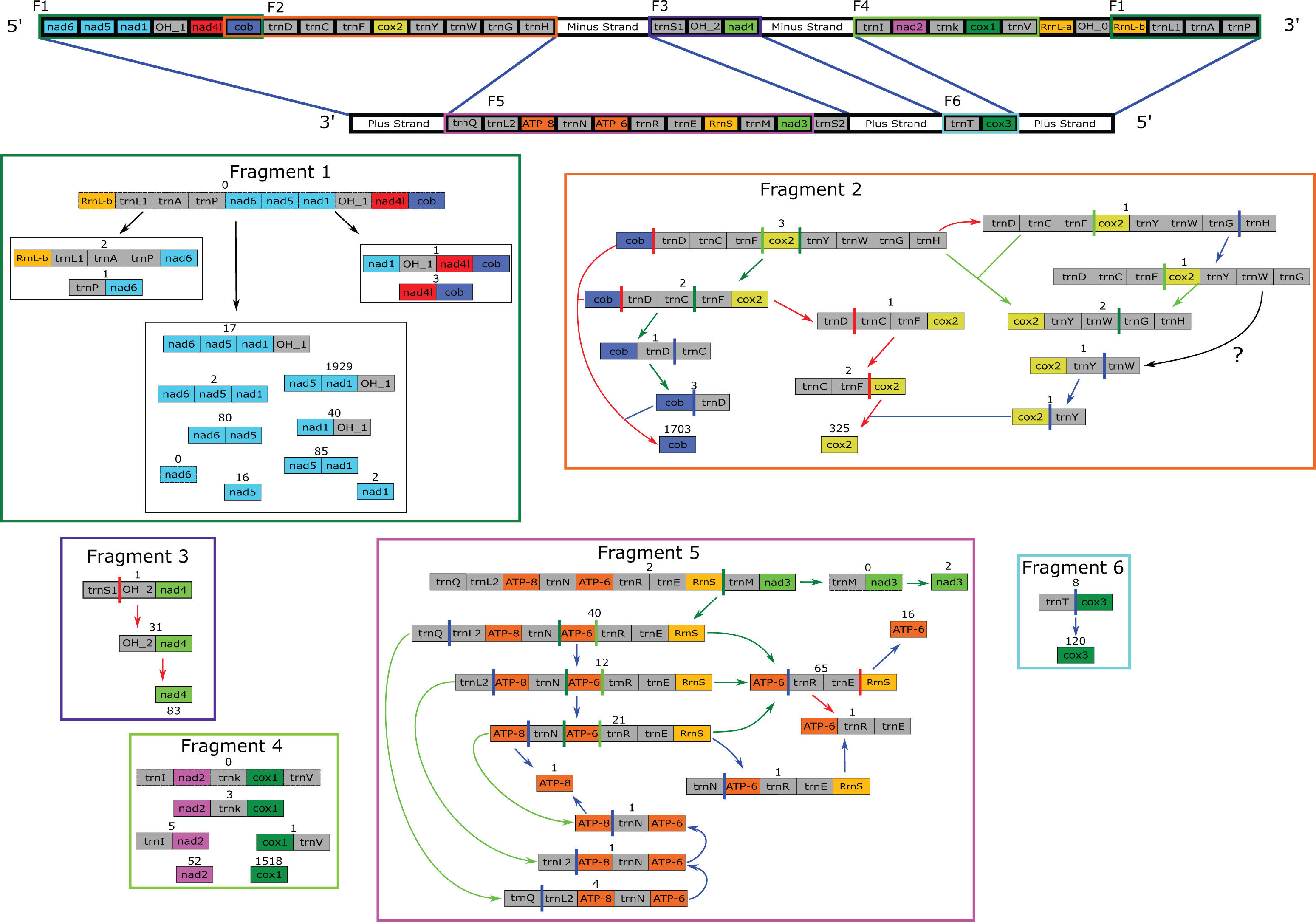
Schematic presentation of the mitochondrial gene trimming process for the transcripts, showing regions of primary transcription (Fragment 1-6) on the plus and minus strand, and the trimming processes of these primary transcripts into pre-mRNA. Numbers above transcripts represent the RNA sequence depth from aligned PacBio IsoSeq ccs data (.pdf file).

**Supplementary Figure 3.**
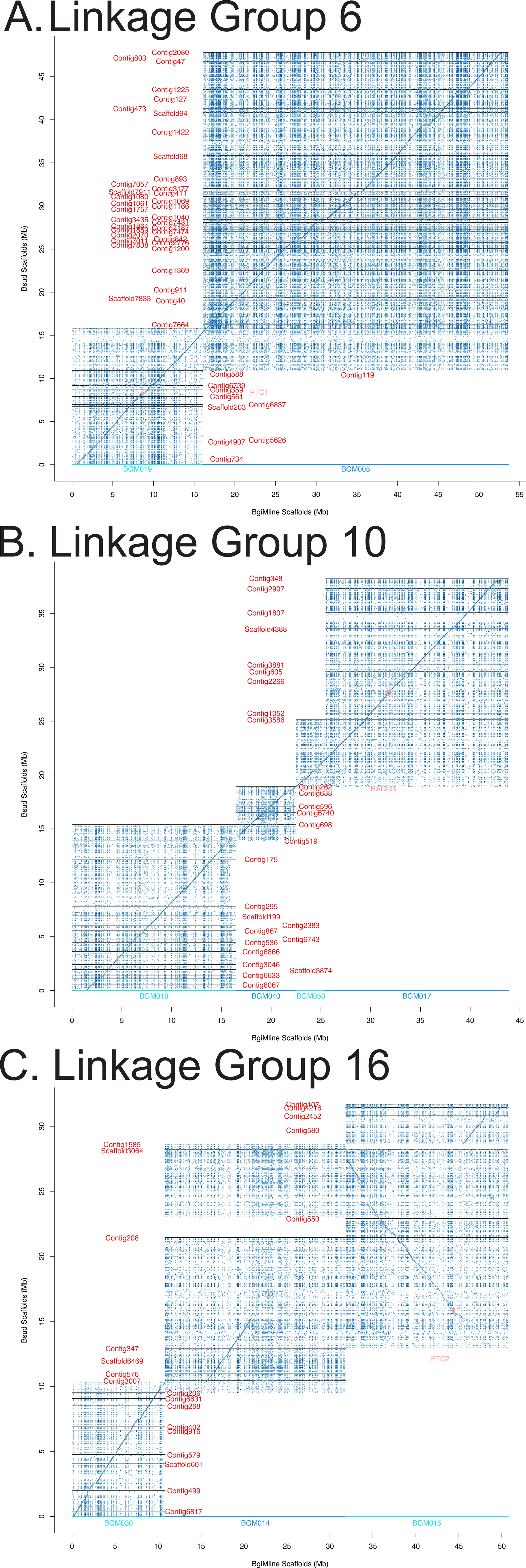
Dot plots of *Biomphalaria sudanica* linkage groups 6 (A), 10 (B) and 16 (C), composed of multiple scaffolds determined using the *B. glabrata* iM line linkage map (Bu et al., 2022). Dots represent 600bp segments; dark blue is ≥97.5% sequence similarity, light blue is ≥90% sequence similarity (.pdf file).

**Supplementary Figure 4.**
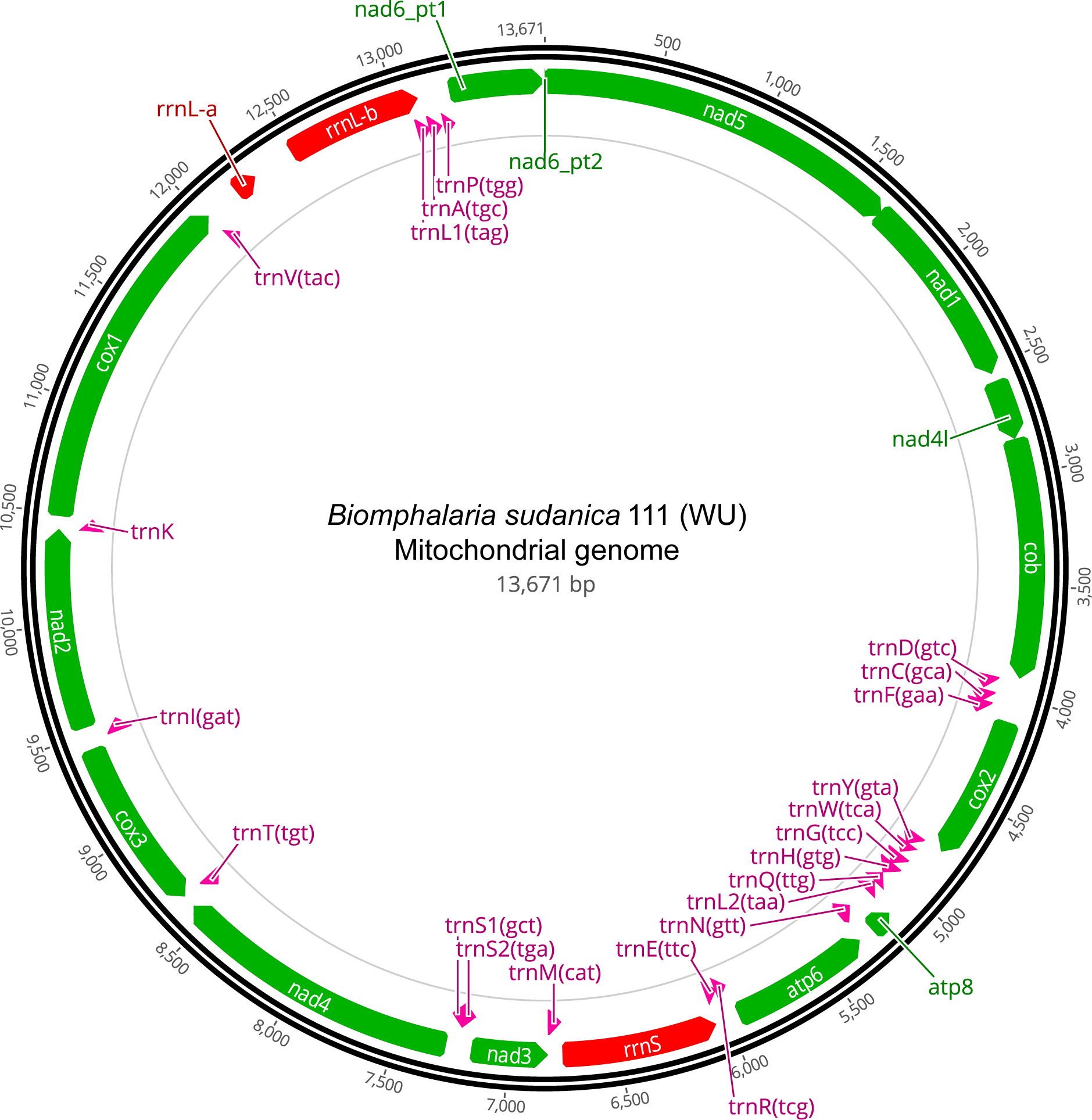
Mitochondrial genome of *Biomphalaria sudanica* with point of origin set to the start of the *nad5* gene (.pdf file).

**Supplementary Figure 5.**
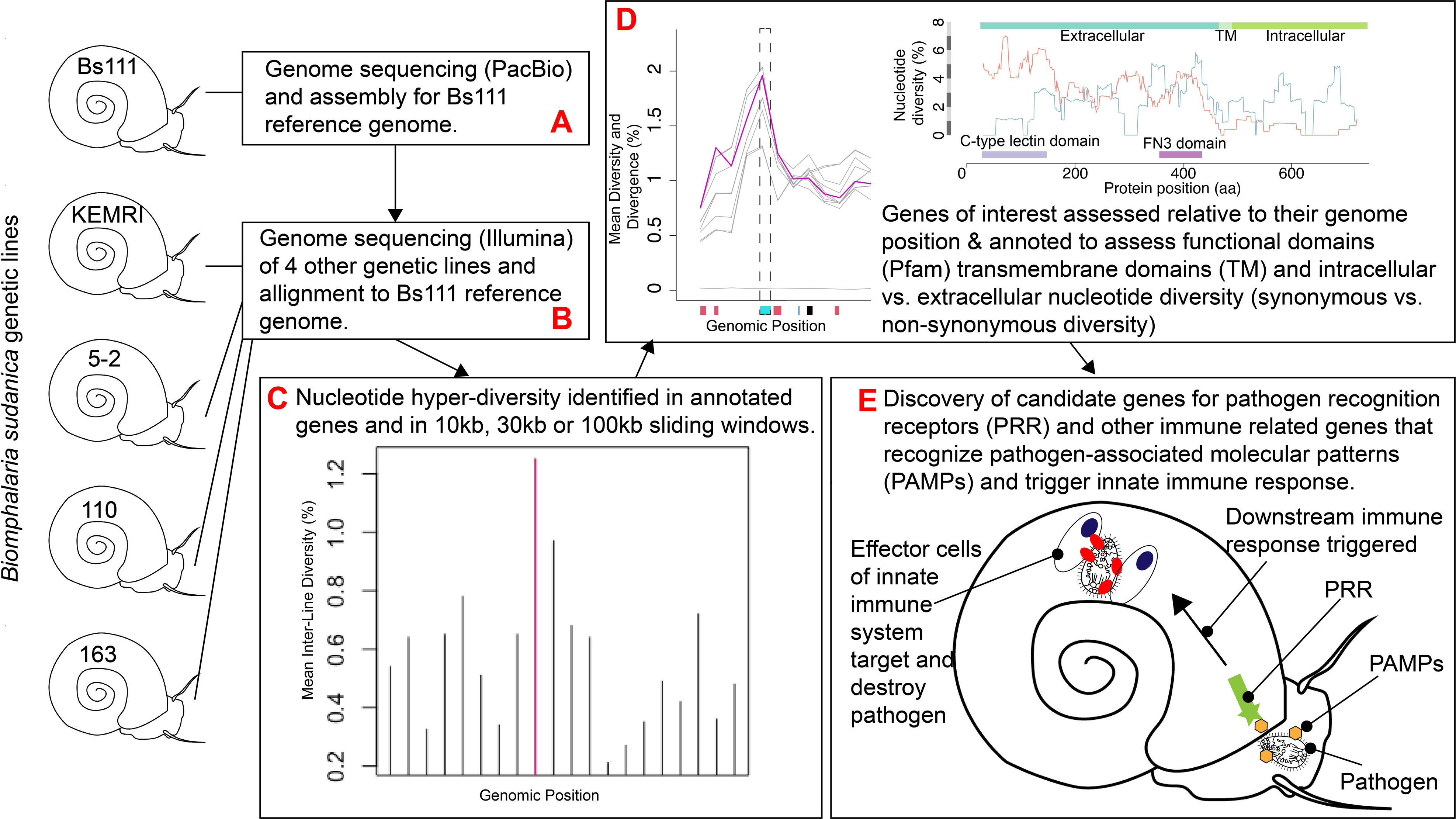
Schematic overview of the methods employed to delimit pathogen recognition receptors (PRR) and other immune genes of *Biomphalaria sudanica* under balancing selection, determined through the analysis of high intraspecific genetic diversity regions, potentially relevant to the resistance and susceptibility of this species to *Schistosoma mansoni*.

