## Supplementary File 5 for "The genome and transcriptome of the snail *Biomphalaria sudanica s.l.*: Immune gene diversification and highly polymorphic genomic regions in an important African vector of *Schistosoma mansoni*"

Species tree generated in Orthofinder using the Species Tree of All Genes (STAG) algorithm, as shown in Figure 1 on the main text. Node support values represent the bipartition proportions in each of the individual species tree estimates. Branch lengths (rounded up to 3d.p.) represent the average number of substitutions per site across all the individual trees inferred from each gene family. The number of (significant/total) gene families expanded (blue) and contracted (red) of Biomphalaria species, and outgroups Elysia marginata and Bulinus truncatus as determined in Café are shown for each internal and terminal node (<0> to <13>).

The following pages show node by node the REVIGO summaries of the GO terms enriched among the protein families with significant Expansion and Contractions. Results for the outgroup species and nodes are not included.

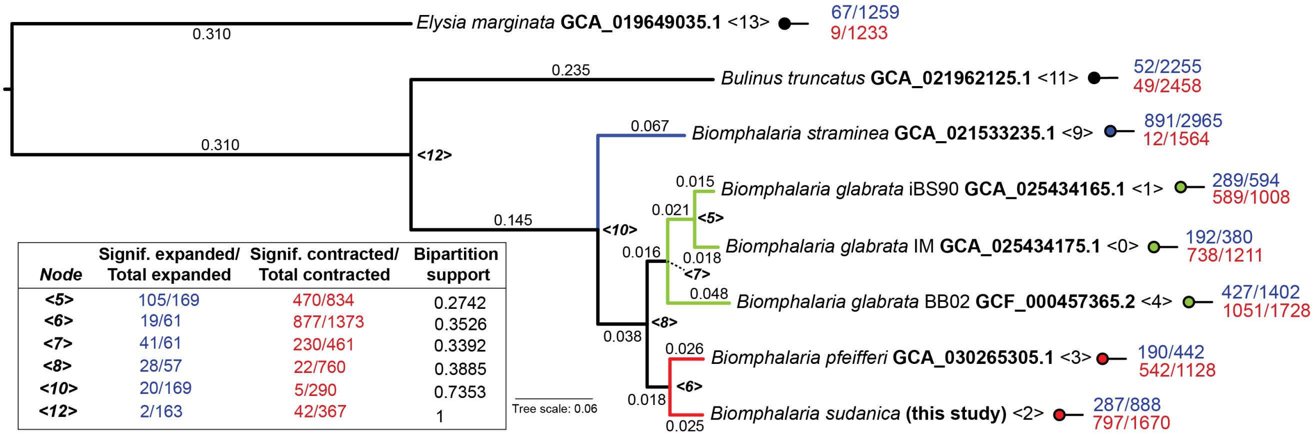


| Node | Signif. expanded/ Total expanded | Signif. contracted/ Total contracted | Bipartition support |
|---|---|---|---|
| <5> | 105/169 | 470/834 | 0.2742 |
| <6> | 19/61 | 877/1373 | 0.3526 |
| <7> | 41/61 | 230/461 | 0.3392 |
| <8> | 28/57 | 22/760 | 0.3885 |
| <10> | 20/169 | 5/290 | 0.7353 |
| <12> | 2/163 | 42/367 | 1 |

Nodes with REVIGO results:
Node 0: Species *B. glabrata*, strain IM (GCA_025434175.1).
Node 1: Species *B. glabrata*, strain iBS90 (GCA_025434165.1).
Node 2: Species *B. sudanica* (this study).
Node 3: Species *B. pfeifferi* (GCA_030265305.1).
Node 4: Species *B. glabrata*, strain BB02 (GCF_000457362.2).
Node 5: Common ancestor between *B. glabrata* strains iBS90 and IM.
Node 6: Common ancestor between the African Biomphalaria species.
Node 7: Common ancestor between the three *B. glabrata* strains.
Node 8: Common ancestor between the three *B. glabrata* strains and the African species.
Node 9: Species *B. straminea* (GCF_021533235.1).
Node 10: Common ancestor of all Biomphalaria species.

**Node 0**: *Biomphalaria glabrata* IM GCA_025434175.1
Protein Family Expansion

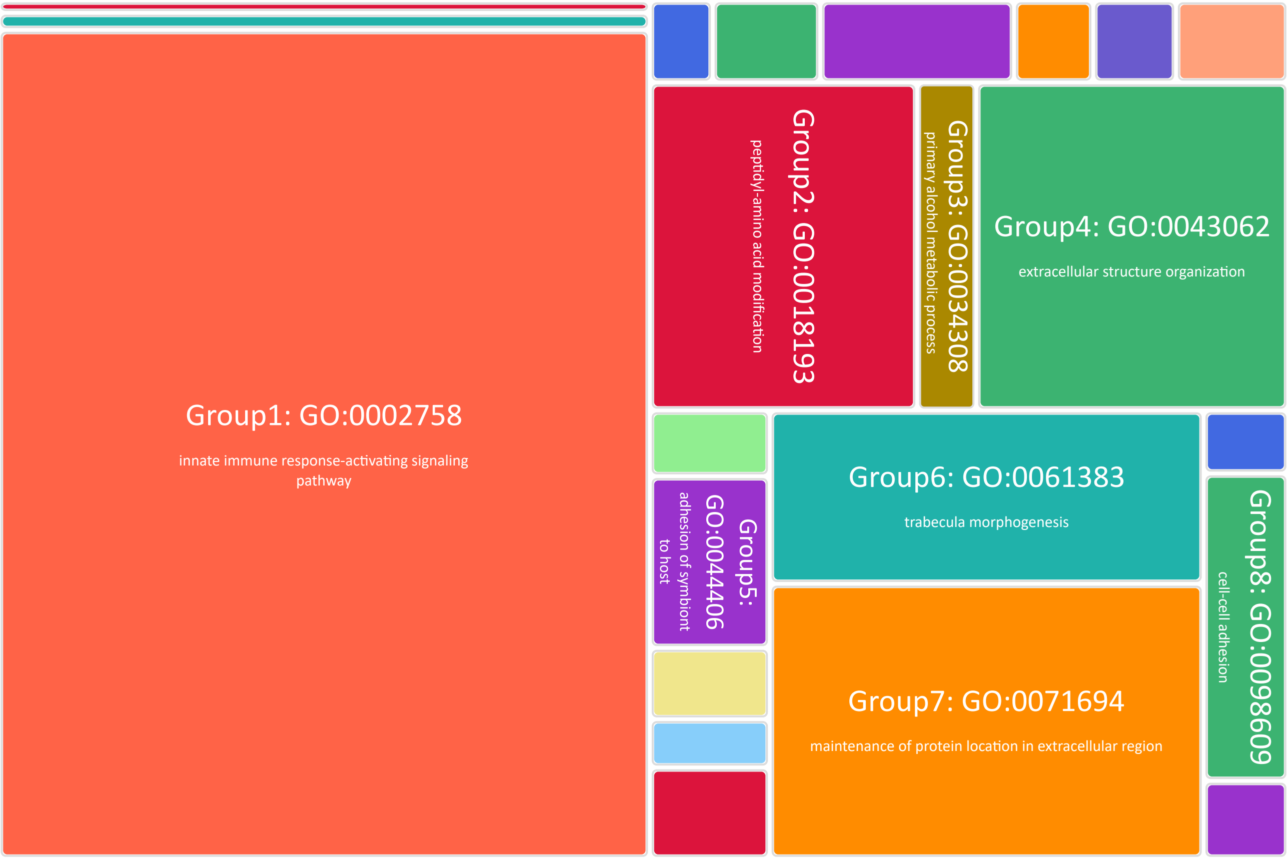


**Node 0**: *Biomphalaria glabrata* IM GCA_025434175.1
Protein Family Contraction

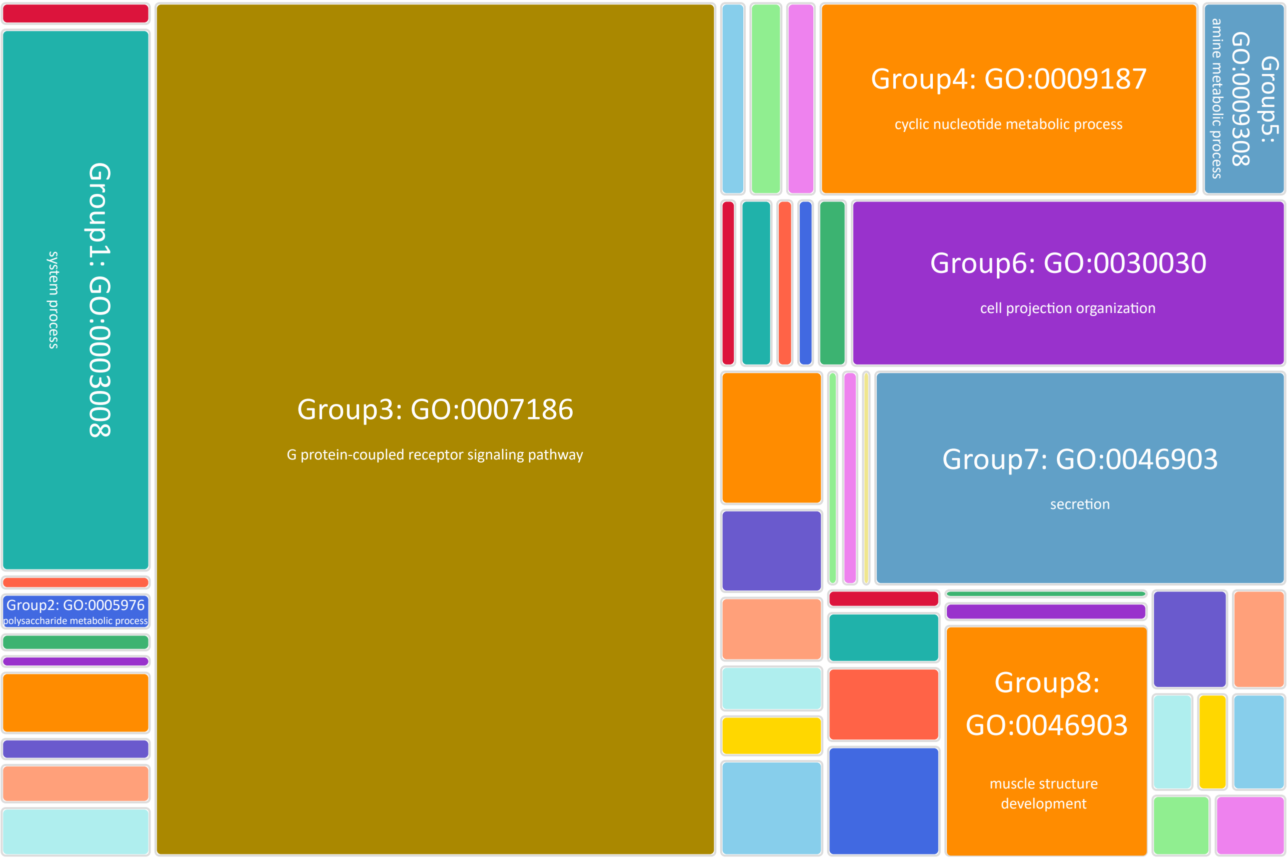

**Node 1**: *Biomphalaria glabrata* iBS90 GCA_025434165.1
Protein Family Expansion

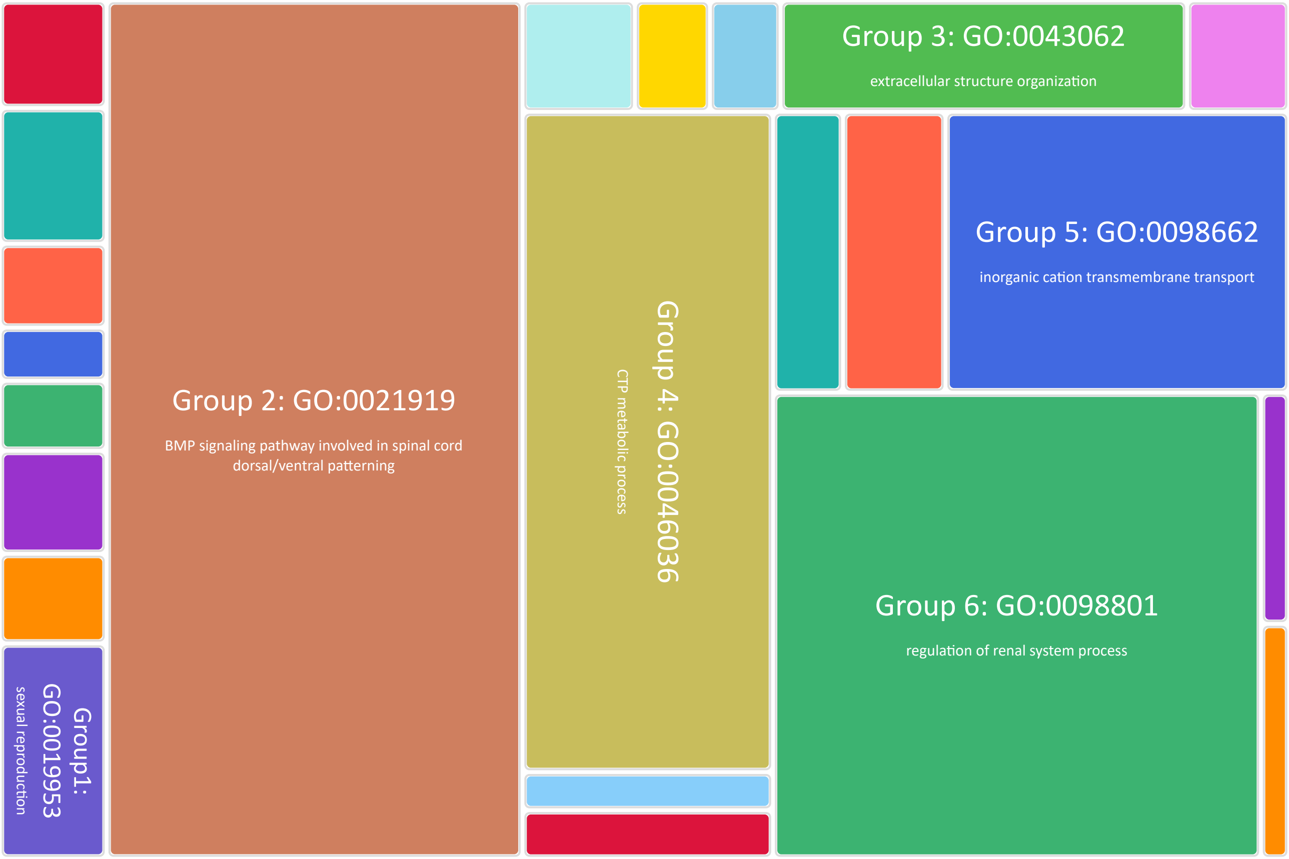


**Node 1**: *Biomphalaria glabrata* iBS90 GCA_025434165.1
Protein Family Contraction

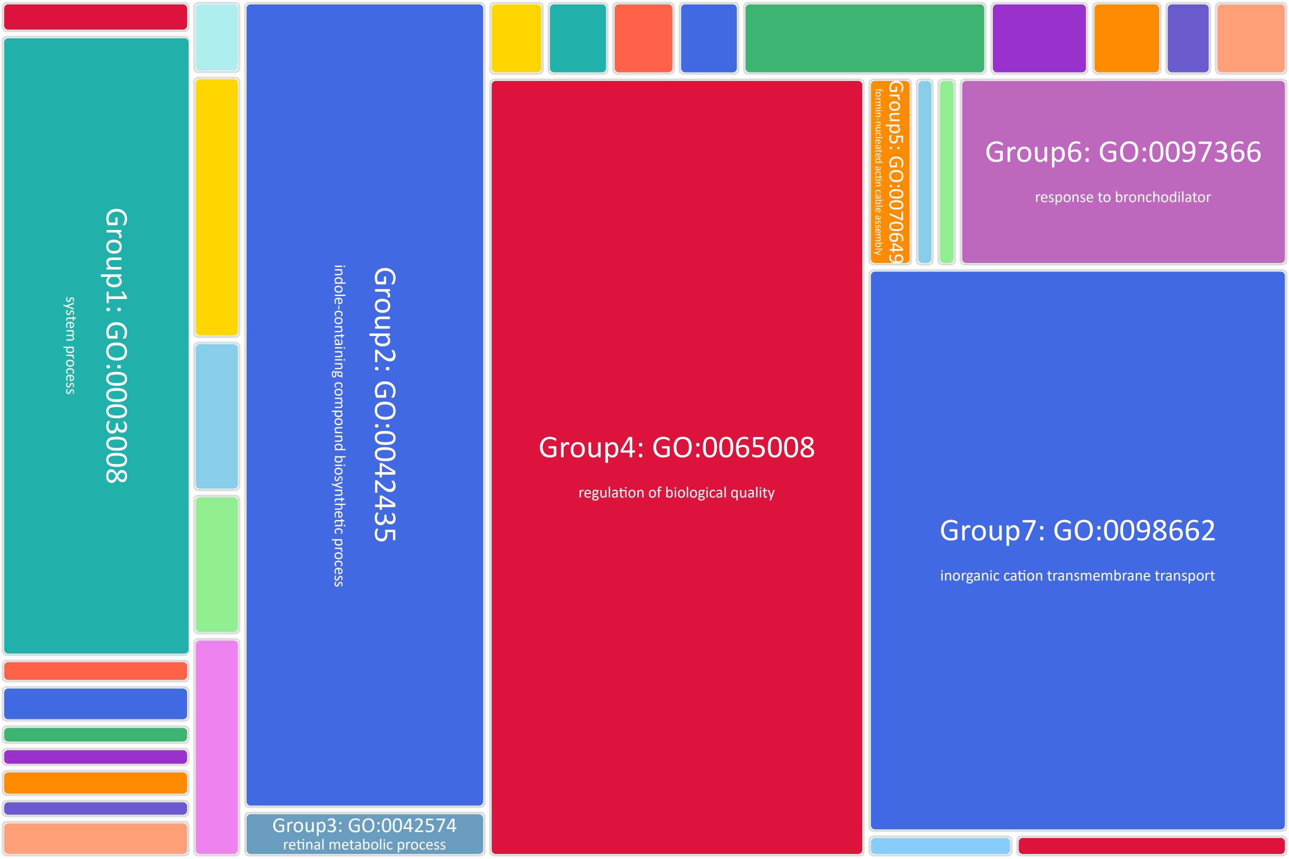

**Node 2**: *Biomphalaria sudanica*
Protein Family Expansion

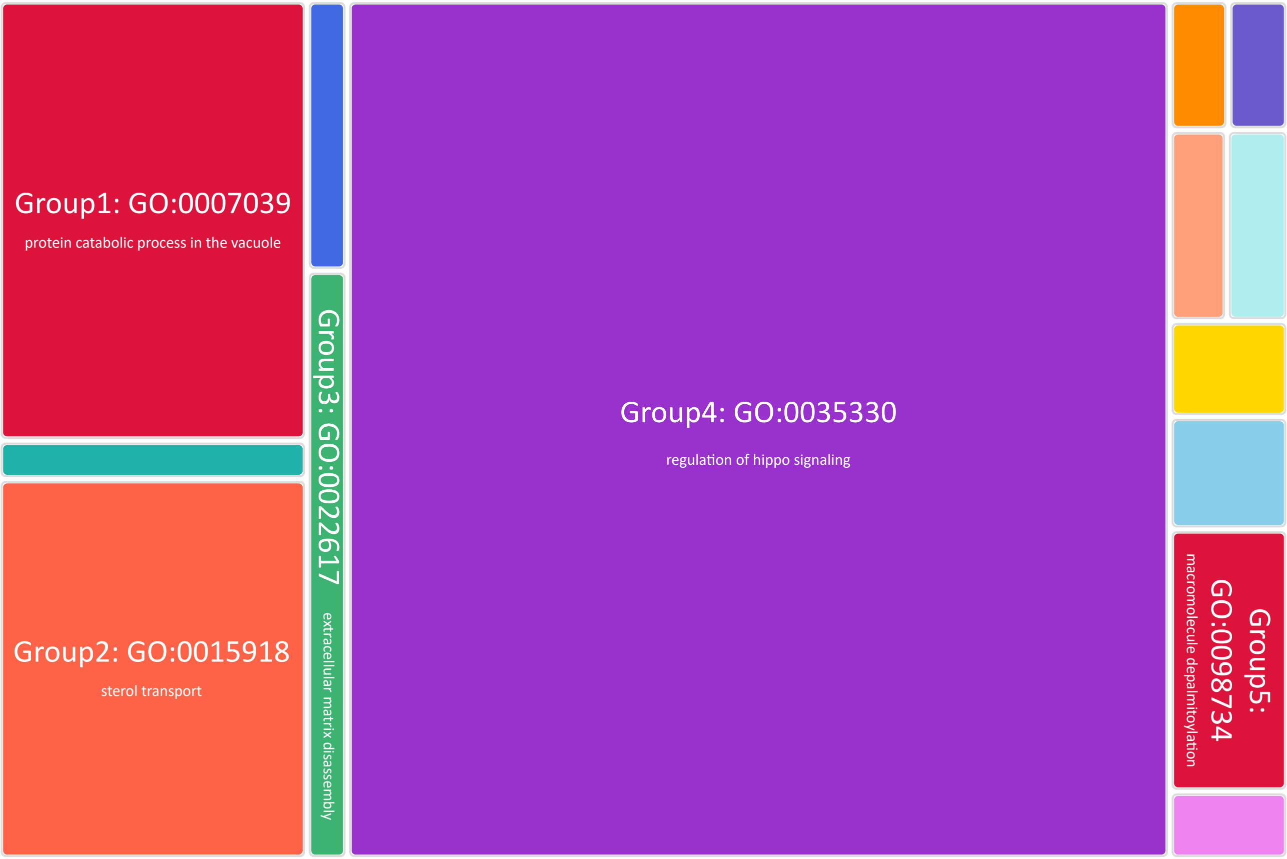


**Node 2**: *Biomphalaria sudanica*
Protein Family Contraction

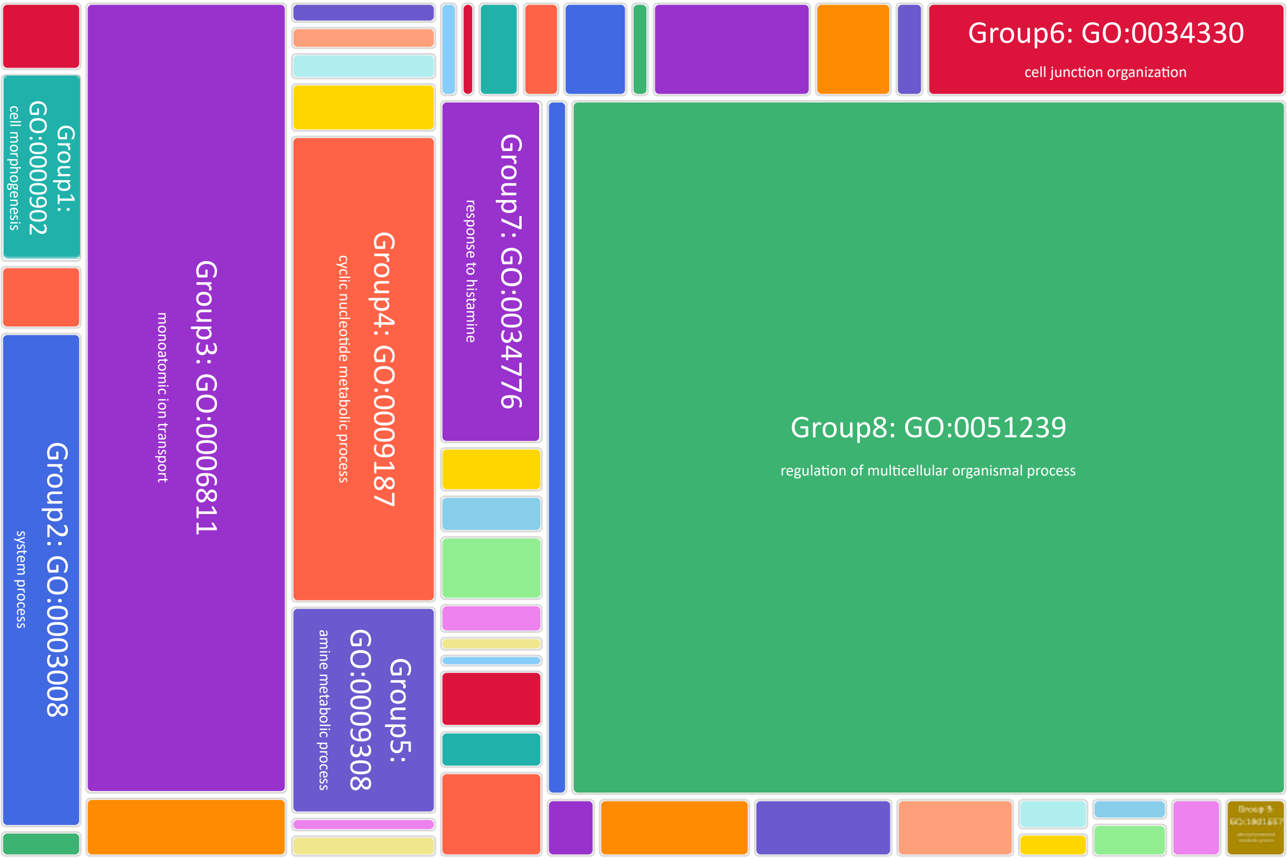

### **Node 3**: *Biomphalaria pfeifferi* GCA_030265305.1
### Protein Family Expansion

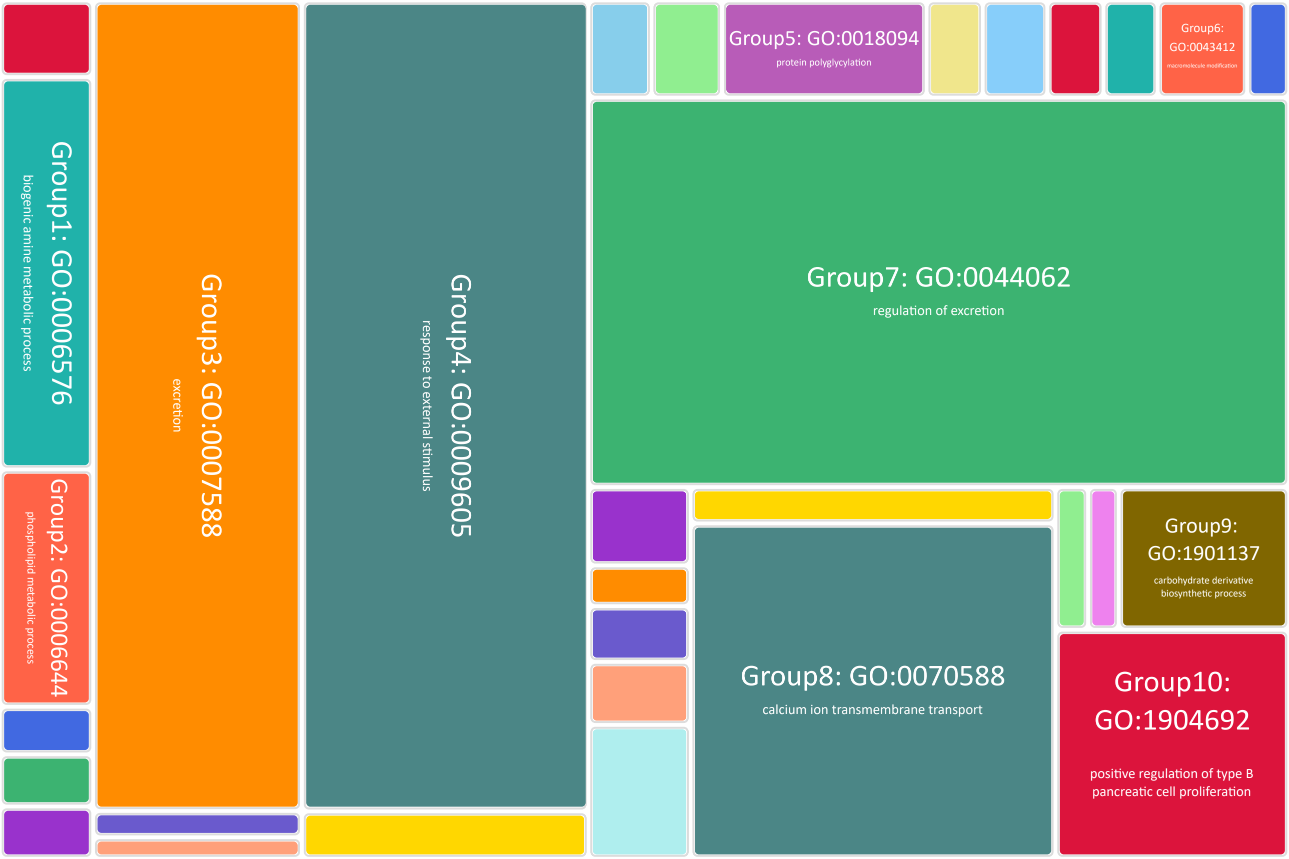


### **Node 3**: *Biomphalaria pfeifferi* GCA_030265305.1
### Protein Family Contraction

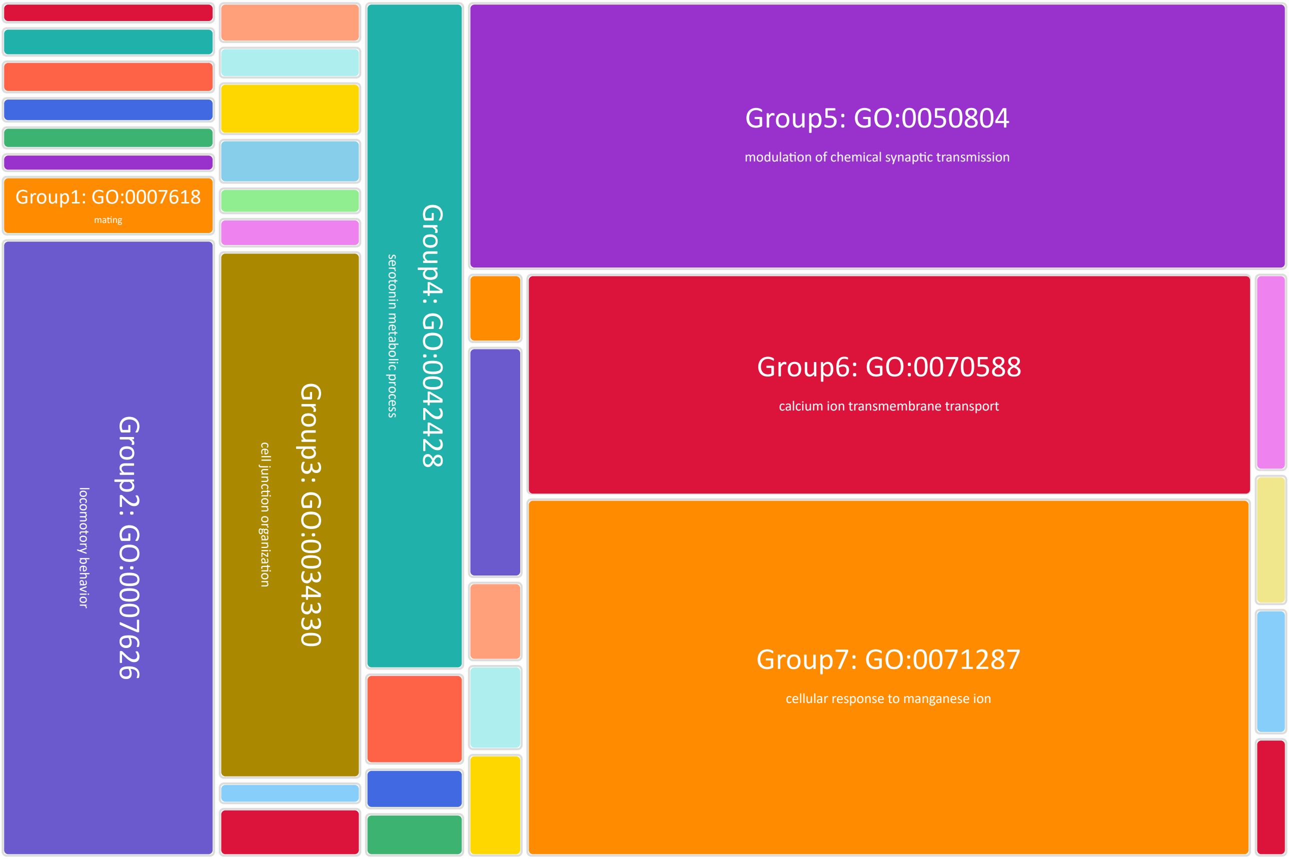

**Node 4**: *Biomphalaria glabrata* BB02 GCF_000457362.2
Protein Family Expansion

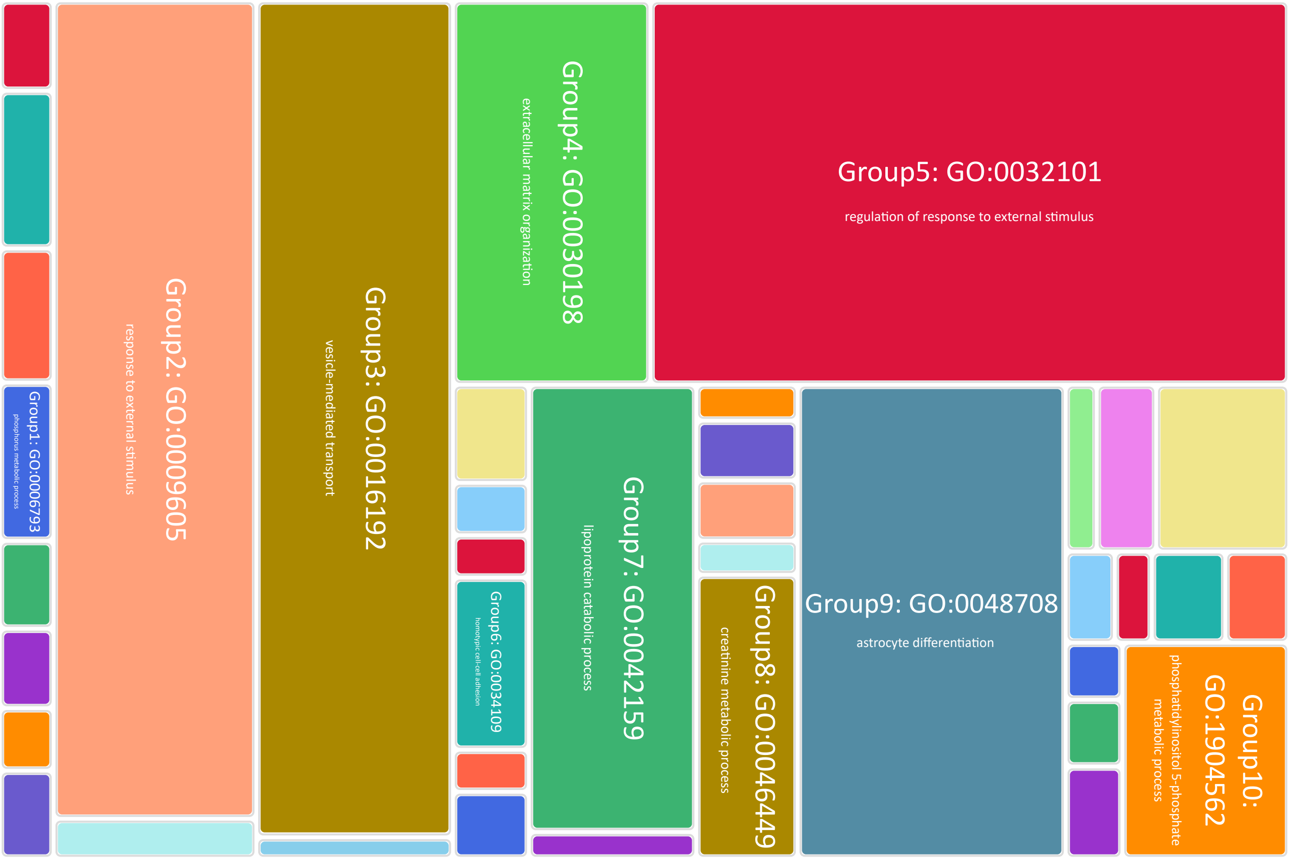


**Node 4**: *Biomphalaria glabrata* BB02 GCF_000457362.2
Protein Family Contraction

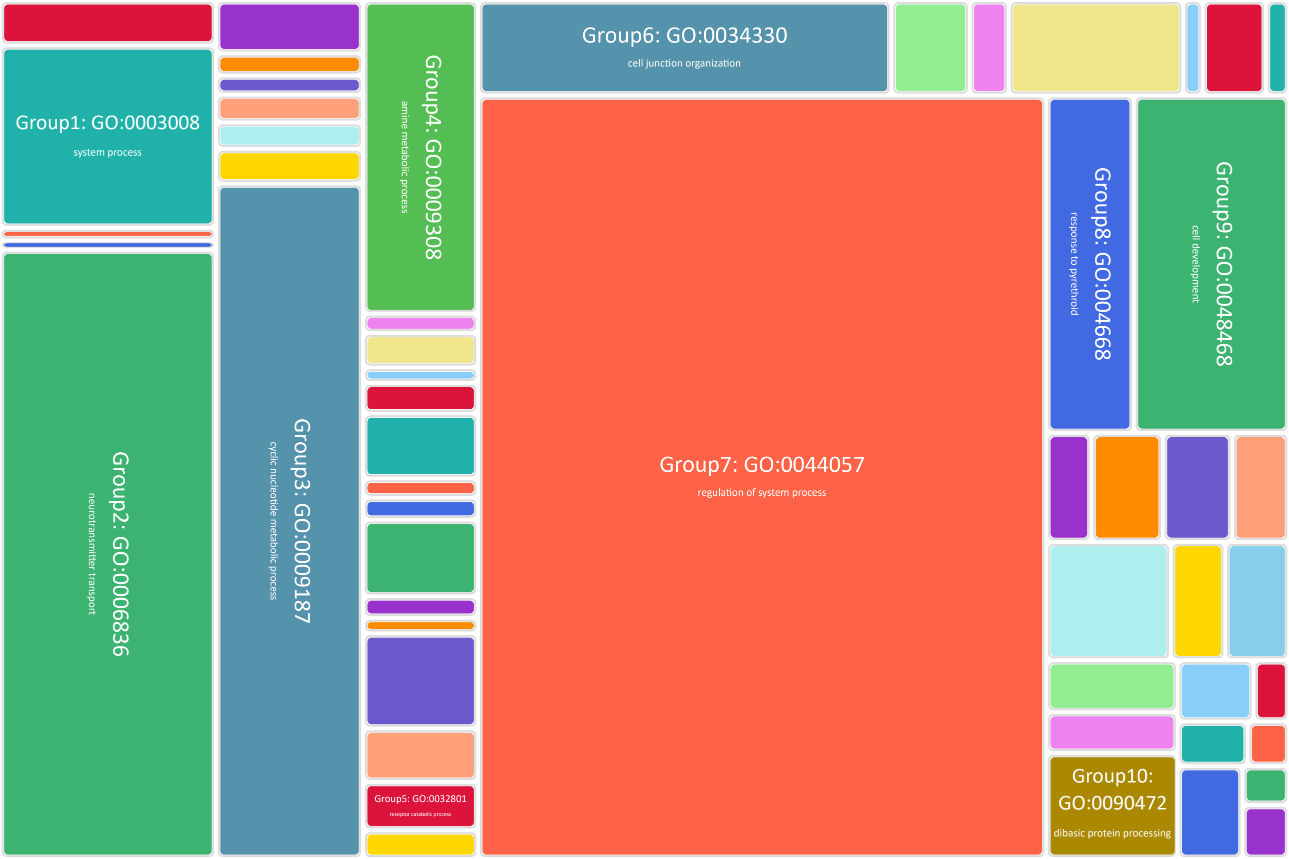

### **Node 5**: *Biomphalaria glabrata* iBS90 x IM ancestor Protein Family Expansion

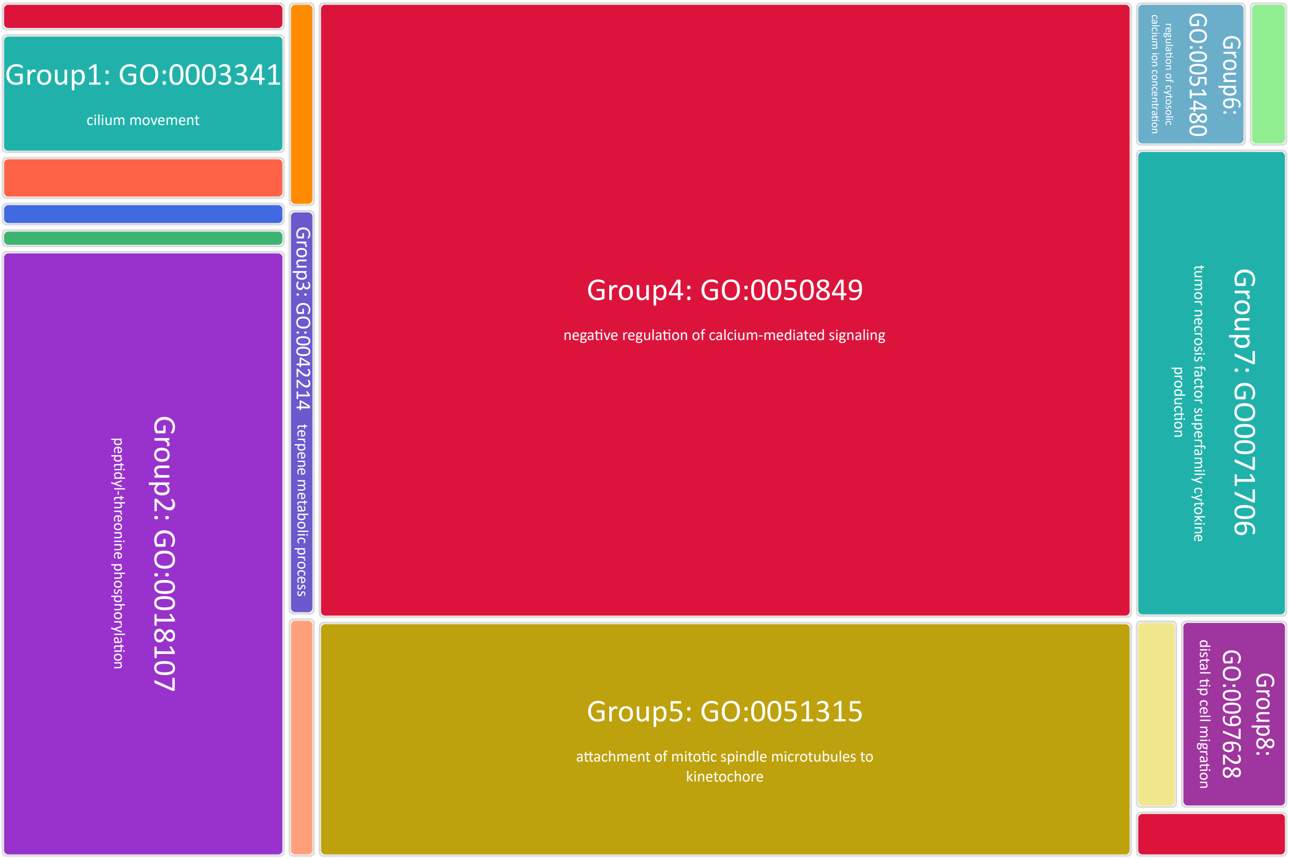


### **Node 5**: *Biomphalaria glabrata* iBS90 x IM ancestor Protein Family Contraction

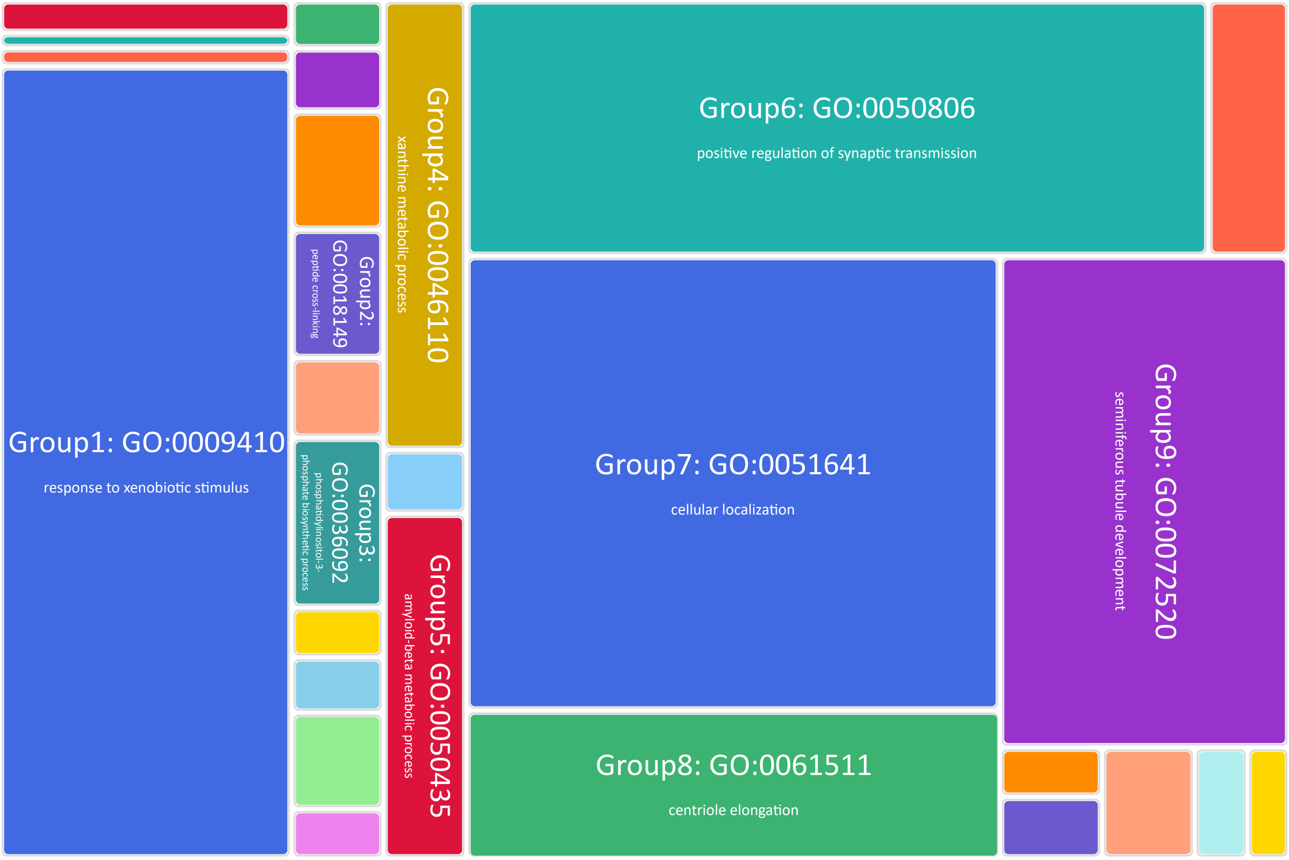

**Node 6**: African *Biomphalaria sp.*
Protein Family Expansion

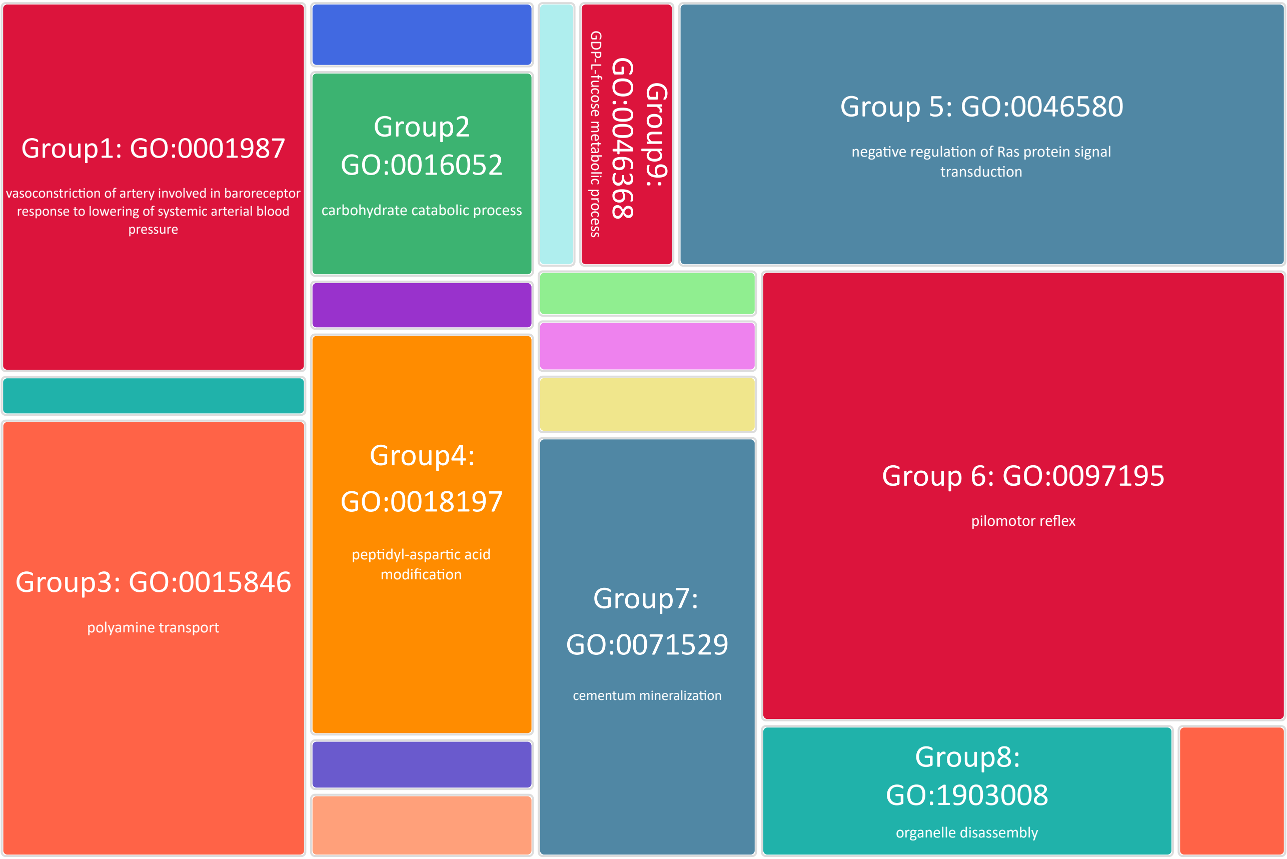


**Node 6**: African *Biomphalaria sp.*
Protein Family Contraction

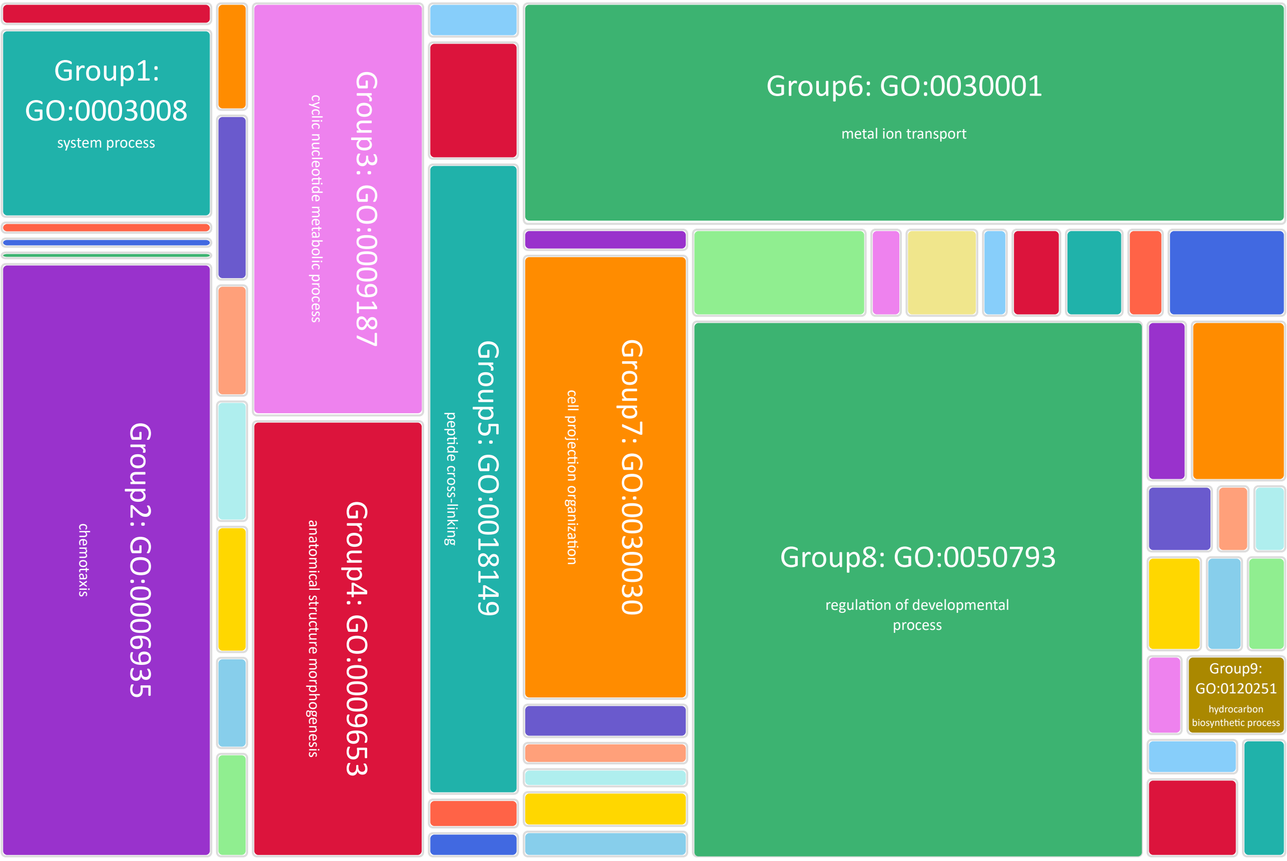

### **Node 7**: *Biomphalaria glabrata* strains ancestor Protein Family Expansion

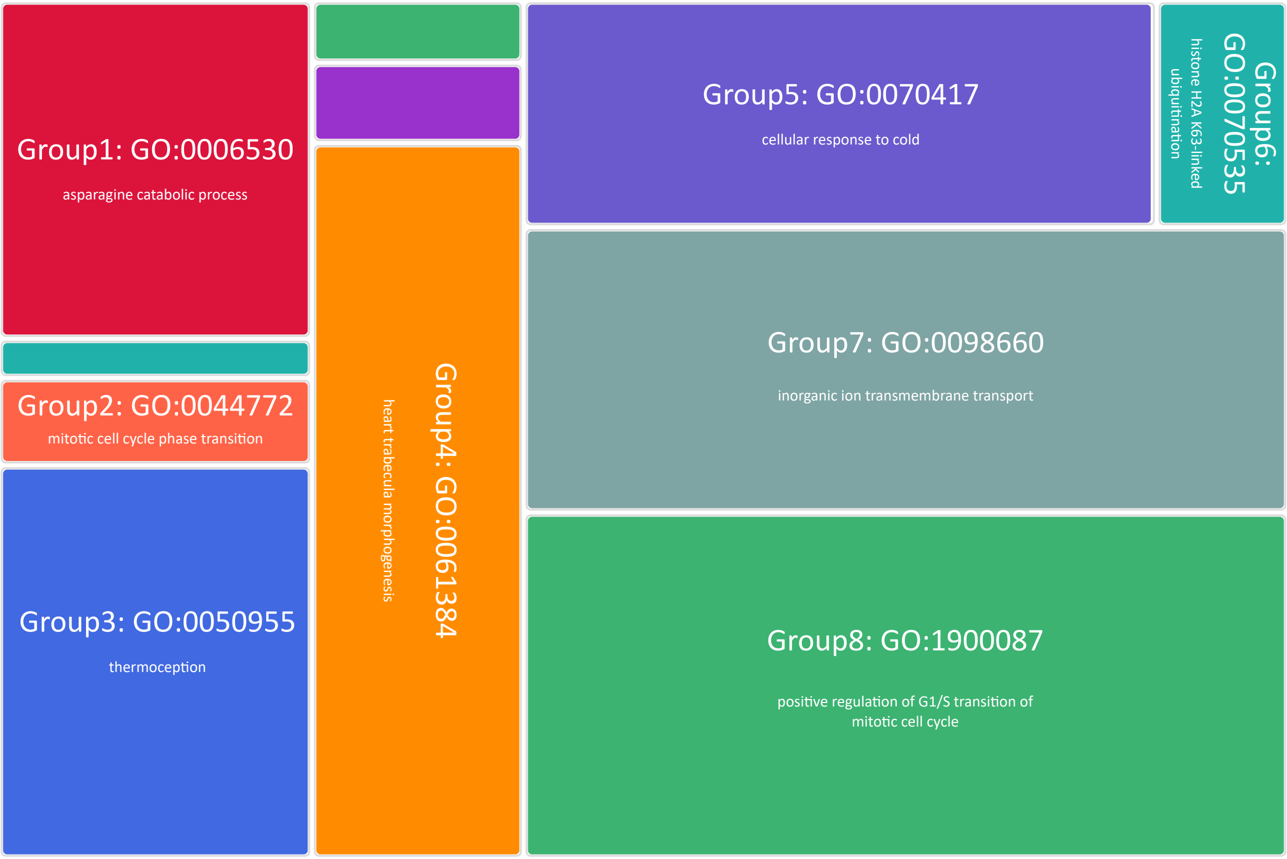


### **Node 7**: *Biomphalaria glabrata* strains ancestor Protein Family Contraction

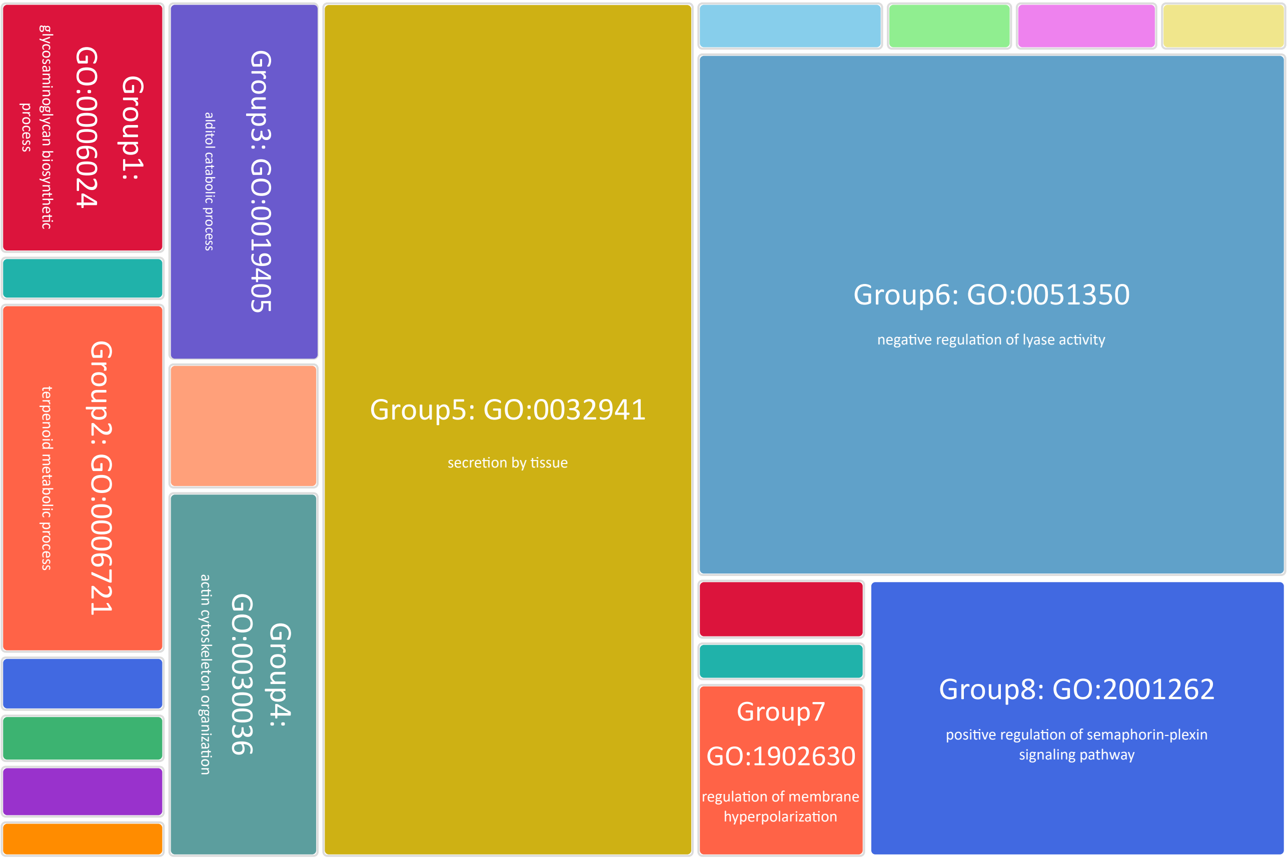

### **Node 8**: *Biomphalaria glabrata* x African species ancestors Protein Family Expansion

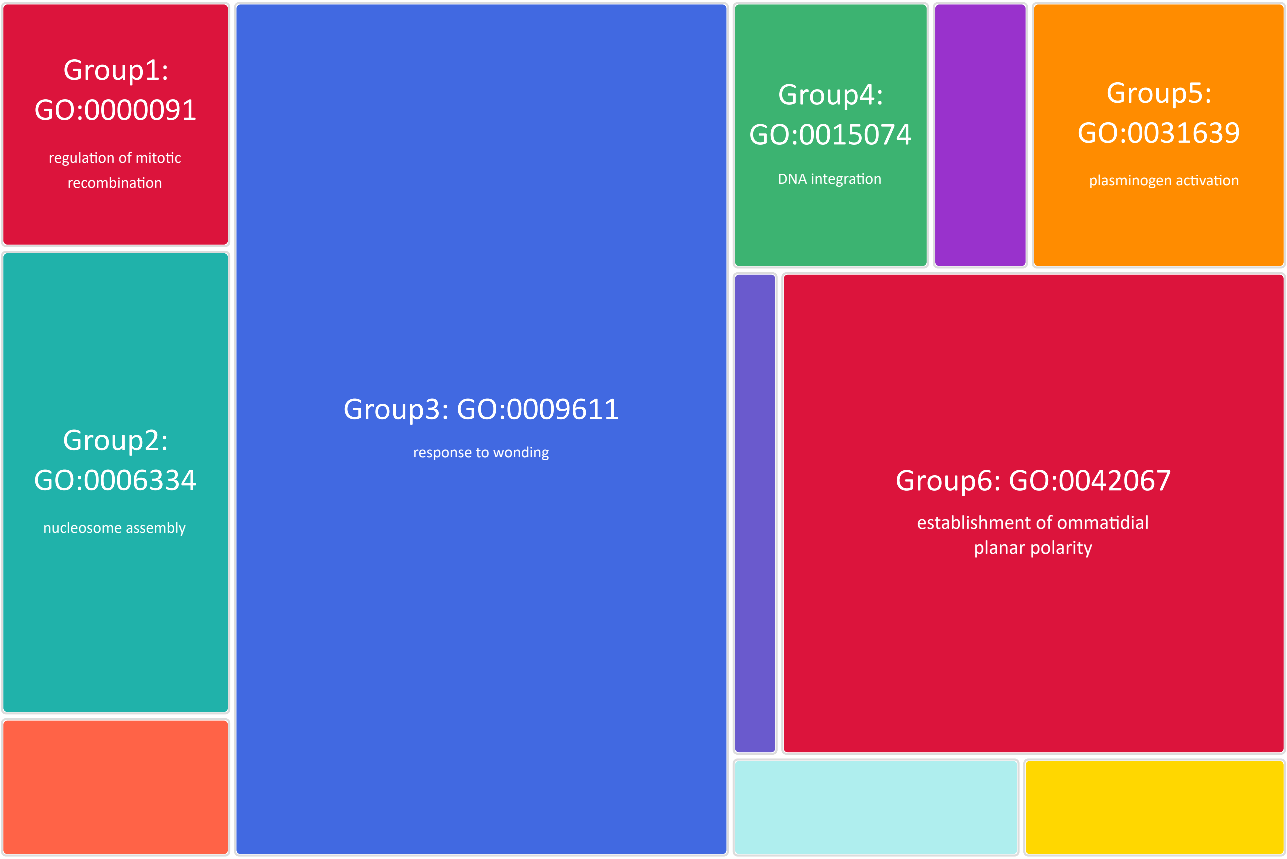


### **Node 8**: *Biomphalaria glabrata* x African species ancestors Protein Family Contraction

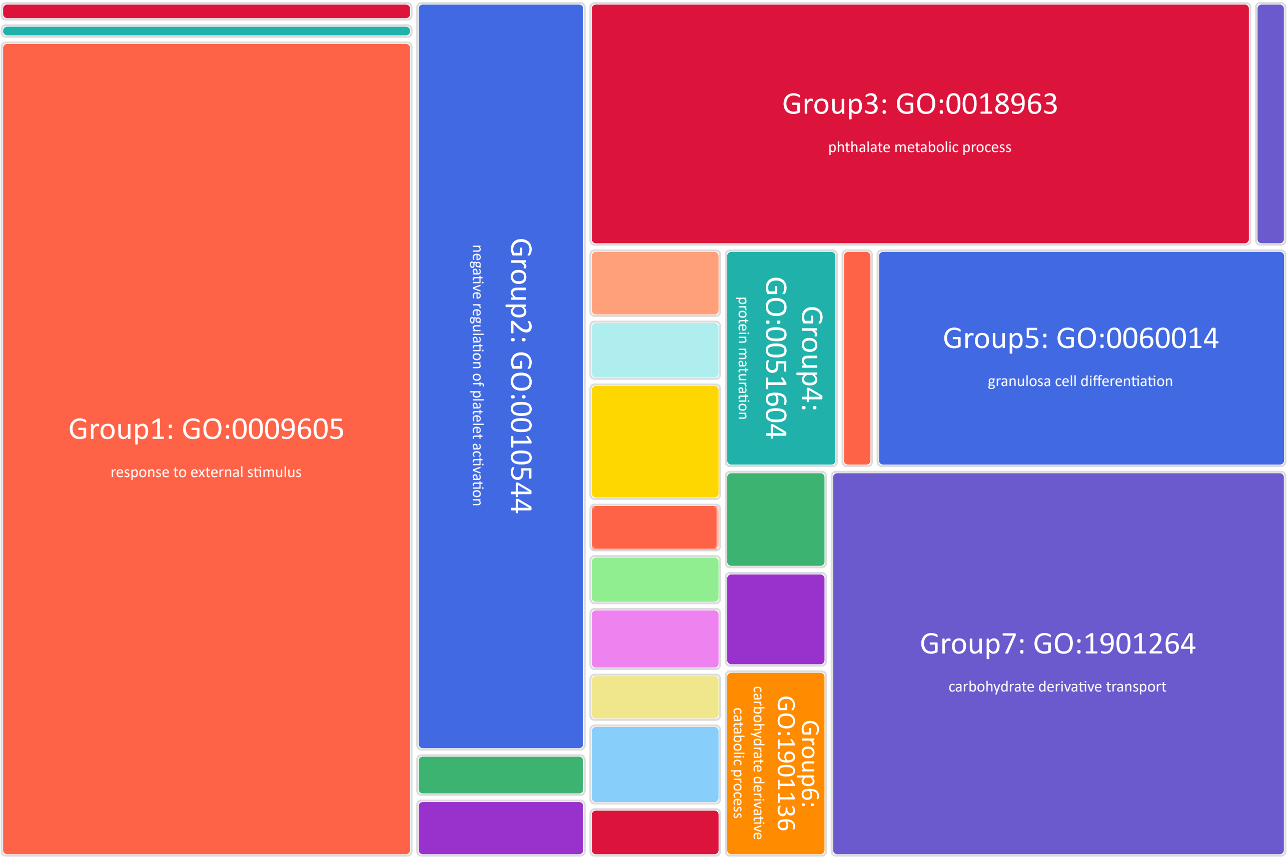

### **Node 9**: *Biomphalaria straminea* GCF_021533235.1 Protein Family Expansion

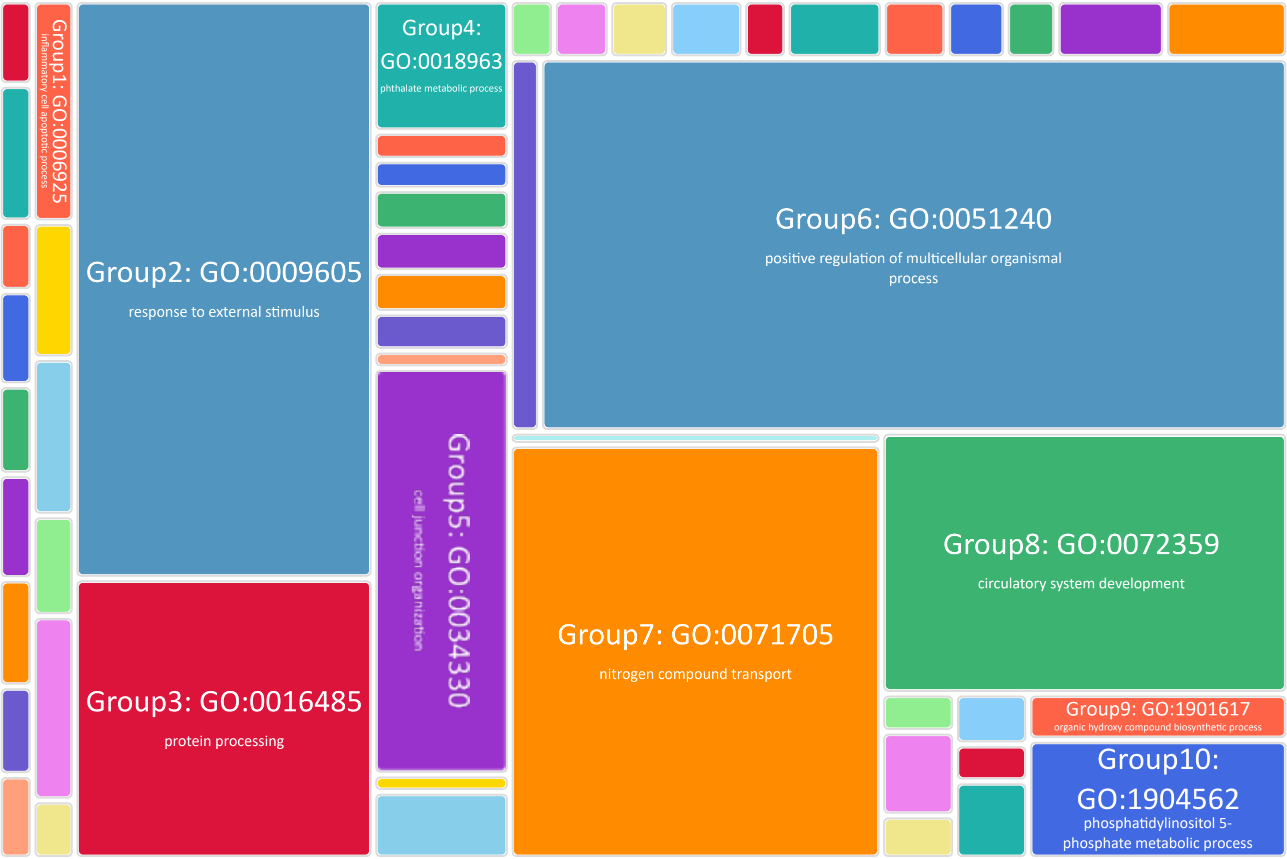


### **Node 9**: *Biomphalaria straminea* GCF_021533235.1 Protein Family Contraction

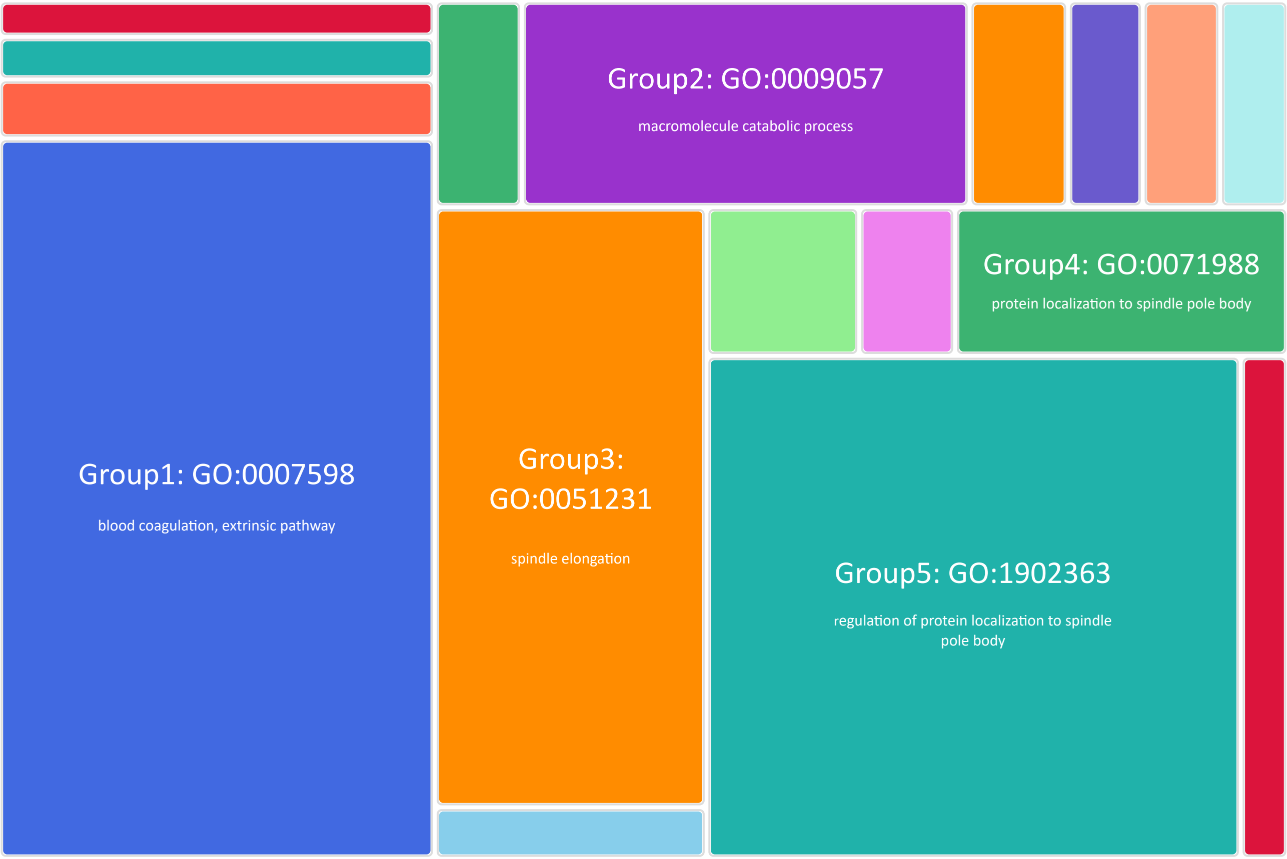

**Node 10**: Biomphalaria common ancestor
Protein Family Expansion

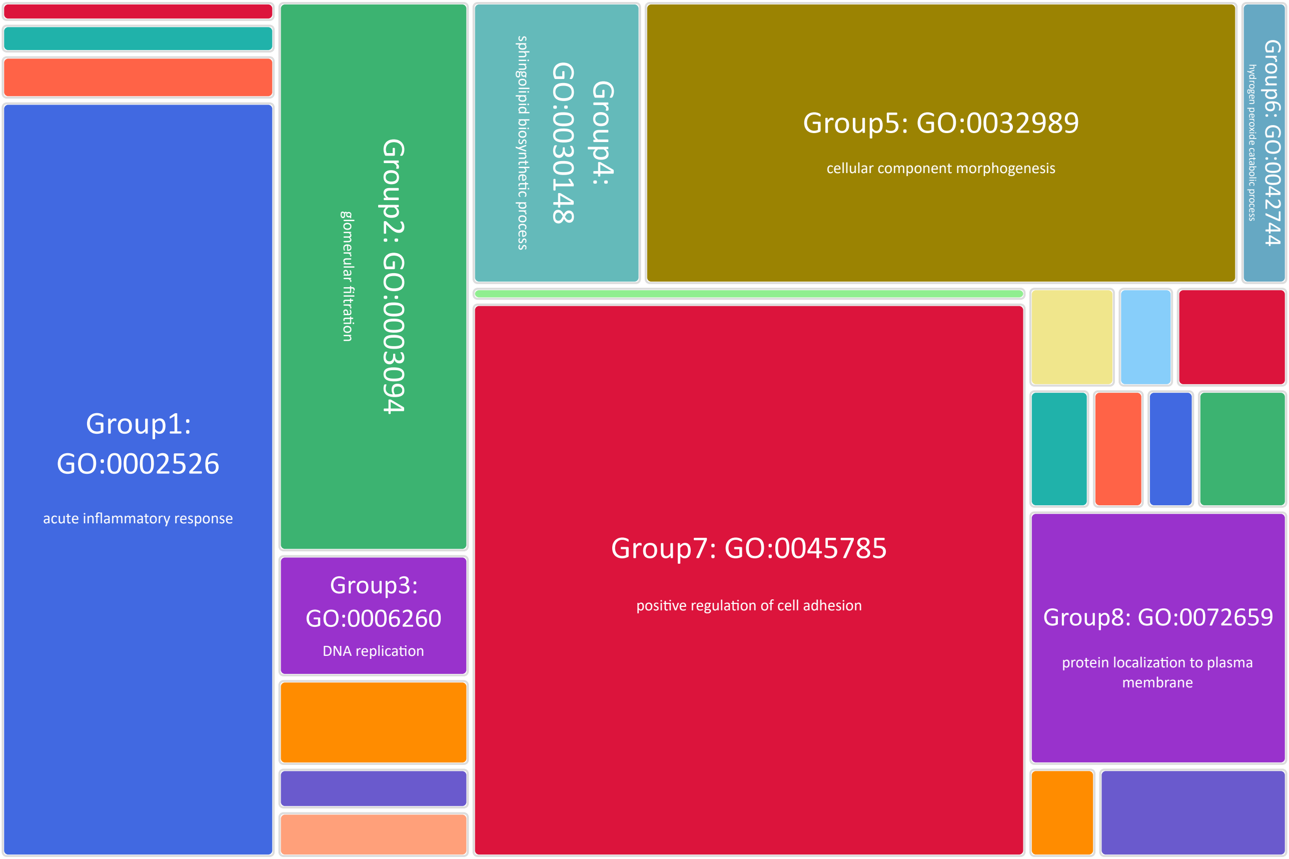


**Node 10**: Biomphalaria common ancestor
Protein Family Contraction

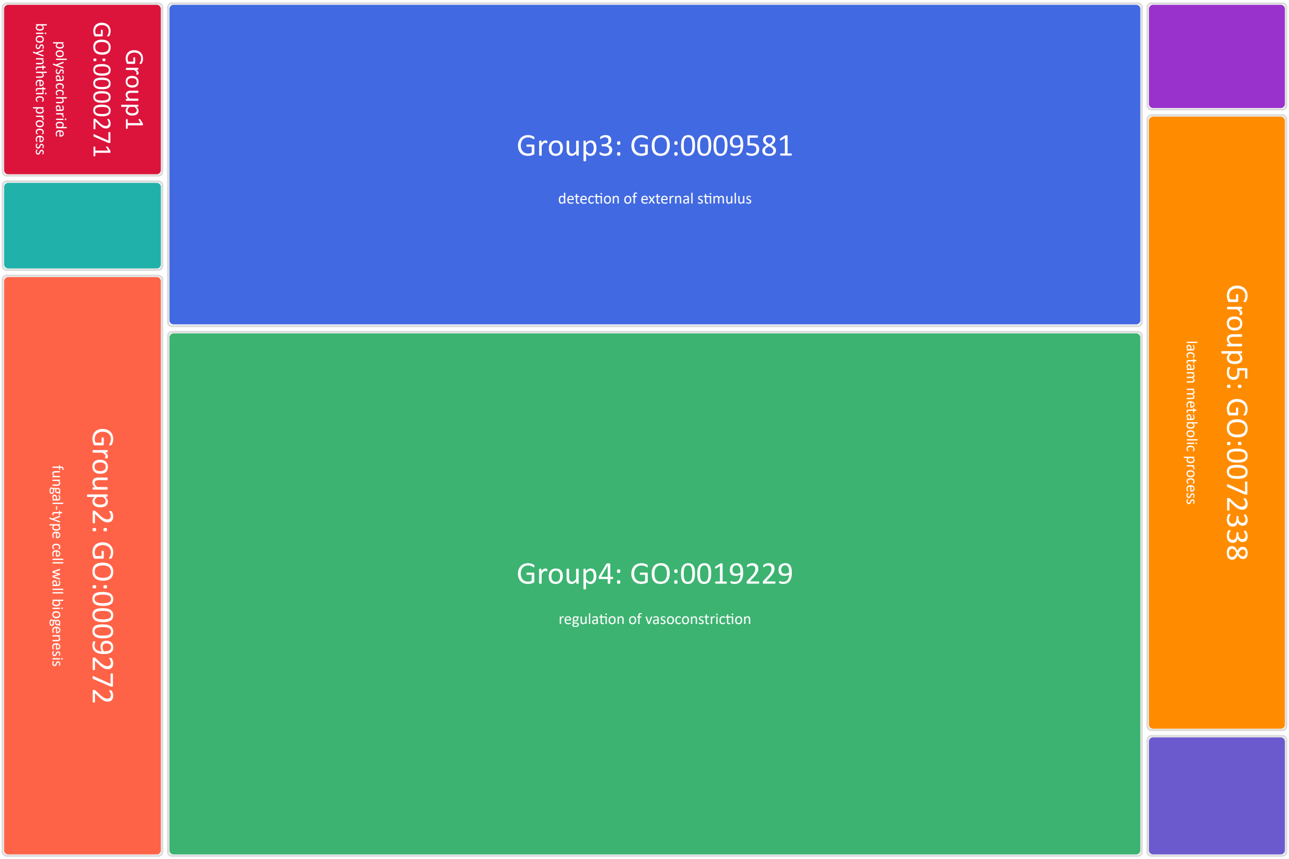
