## Supplementary File 6 for "The genome and transcriptome of the snail *Biomphalaria sudanica s.l.*: Immune gene diversification and highly polymorphic genomic regions in an important African vector of *Schistosoma mansoni*"

RNA was extracted from *Biomphalaria sudanica* inbred line “111”. RNA for long and short read sequencing was obtained from 8 and 2 RNA extractions, respectively (see Table A). To capture potentially differentially expressed genes during *B. sudanica* life stages and possible environmental stressors, RNA extracted from the 10 samples included both individual and multiple snails (see Table A) representing 18 juvenile snails sized 4-5 and 12 adult snails >8 mm in size. Of these, we exposed 21 (12 juvenile 4-5mm and nine adult >8mm) to 10 *Schistosoma mansoni* miracidia from the inbred line population supplied by NIH-NIAID Schistosomiasis Resource Centre (Lewis et al. 2008). Snails were placed in 12-well ELISA plates filled with 2 ml of snail conditioned water and challenged by adding 10 miracidia to each well. After 24 hours, six juvenile (s8, Table A) and six adult (b7 and b8, Table A) snails were sacrificed as detailed below. The remaining six juvenile and three adult snails were moved from ELISA plates and into two respective tanks containing 8 liters of snail conditioned water for the following 48 hours (as detailed below) until reaching the 72 hour post-exposure time point where snails were sacrificed.

| Table A. Biomphalaria sudanica samples used for PacBio and Illumina whole genome sequencing | | | |
| --- | --- | --- | --- |
| **Sample ID** | **Snail size** | **Number of snails** | **Snail treatment group** |
| **PacBio Sequel II** | |  |  |
| s1 | 4-5mm | 3 | NA |
| s4 | 4-5mm | 3 | NA |
| s8 | 4-5mm | 6 | 24hrs post challenged w/ BRI-NMRI *S. mansoni* (D=10) |
| s9 | 4-5mm | 6 | 72hrs post challenged w/ BRI-NMRI *S. mansoni* (D=10) |
| b5 | >8mm | 1 | NA |
| b7 | >8mm | 3 | 24hrs post challenged w/ BRI-NMRI *S. mansoni* (D=10) |
| b9 | >8mm | 3 | 72hrs post challenged w/ BRI-NMRI *S. mansoni* (D=10) |
| b11 | >8mm | 1 | Heat stressed (6hrs @ 32C) |
| **Illumina sequencing** | |  |  |
| b2 | >8mm | 1 | NA |
| b8 | >8mm | 3 | 24hrs post challenged w/ BRI-NMRI *S. mansoni* (D=10) |

A single adult snail (b11, Table A) was exposed to a heat stress, which was achieved by placing the snail in a single well of a 12 well ELISA plate containing 2 ml of snail conditioned water, and placed in an incubator at 32ºC for six hours, following which time it was sacrificed as detailed below. Six juvenile and two adult snails were removed directly from breeding tanks and were not exposed to schistosomes or heat stress.

To sacrifice snail(s) for both individual (Table A: b5, b11, b2) and pooled (Table A: s1, s4, s8, s9, b7, b9, b8) samples, snails were removed directly from breeding tanks, or 12-well ELISA plates for exposure groups (as detailed above) and placed directly in liquid nitrogen to flash freeze for at least 20 seconds. Samples destined for PacBio sequencing were pulverized using a sterile pestle and mortar and 1 ml of TRIzol® (Ambion, Carlsbad CA, US) or those destined for Illumina sequencing bead-rupted (2.8mm ceramic beads, (Omni International, Kennesaw GA, US)) in 500 µl of TRIzol® (Ambion, Carlsbad CA, US) using the Omni Beadrupter 12 (Omni International, Kennesaw GA, US) (speed = 6.00, cycle time = 45 seconds, number of cycles = 1). Samples were placed immediately on dry ice to transfer to storage at -80ºC until final RNA extraction.

RNA extraction followed the TRIzolTM reagent (Invitrogen^TM^, ThermoFisher Scientific, MA USA) manufacturers instructions. Isolation and purification was performed using the Qiagen RNeasy Mini kit (Qiagen, MD, USA) followed by an ethanol-sodium acetate precipitation to further purify and concentrate total RNA. Precipitated RNA was eluted in RNase free water.

**Quantification**

Total RNA was quantified using Qubit RNA Broad Range (BR) Assay Kit and RNA integrity checked using the Qubit RNA IQ Assay kit using the Qubit 4 Flourometer (Invitrogen^TM^, ThermoFisher Scientific, MA USA) and 2100 Bioanalyzer (Agilent, CA, USA). Purity and presence of contaminants in RNA samples were assessed by measuring 260/230 and 260/280 absorbance values on a Nanodrop to check that samples sat within the expected range for pure RNA, ~1.8-2.2 and ~2.0, respectively.
